# FunctionaL Assigning Sequence Homing (FLASH) maps phenotype to sequence with deep and machine learning

**DOI:** 10.64898/2026.04.04.715981

**Authors:** Daniel J. Cotter, Marie-Claire Harrison, Arjun Rustagi, Peter L. Wang, Marek Kokot, Allison F. Carey, Sebastian Deorowicz, Julia Salzman

## Abstract

Genome-wide association studies (GWAS) map genetic variation to a reference genome and correlate variants to phenotypes. Yet, GWAS and similar procedures have limitations, including an inability to predict phenotype on variants never seen during the discovery phase and difficulty integrating structural variants. Deep and machine learning alternatives have not been successful at consistent prediction of resistance phenotypes (Hu et al. 2024). Here, we introduce FLASH: a new interpretable, statistically-based deep learning framework that operates directly on raw sequencing reads. In over 35,000 isolates of bacteria, fungi and viruses, FLASH achieves uniformly high accuracy on independent test data, including on variation never seen in training, meeting or exceeding bespoke state of the art methods. FLASH identifies canonical drug targets ab initio and new pan-species predictors of virulence, including those lacking annotation and those only partially aligned to NCBI reference databases. Further, FLASH can predict phenotypes beyond the possibility of GWAS, such as bacterial host range of phage, a task that to our knowledge is impossible today. FLASH is simple to run, highly efficient and constitutes a new approach for predicting gene function and phenotype across the tree of life. It is especially valuable when bioethical concerns and the vast genetic complexity of pathogenic microbes limit the feasibility of experimental validation.

## Introduction

GWAS has been successful at creating bacterial genotype-phenotype maps (Yang et al. 2025; Power et al. 2017; Uffelmann et al. 2021) but it also has significant limitations. First, it assigns variants at specific positions and in general does not treat structural variation such as insertion and deletions or gene presence-absence in a unified framework, an especially acute problem in microbes such as *E. coli* where genomes are extremely plastic (Braz et al. 2020). Second, it is an associative procedure; it does not perform phenotype prediction, and cannot rank the contribution of variation in distinct loci, mainly because its statistical tests operate at each position independently. Third, it is not possible to associate phenotype with new sequences or genomes where variants have not been previously observed; Fourth, GWAS is not an option for predicting complex microbial phenotypes (e.g. host range of phage, of emerging clinical and environmental importance (Sanmukh et al. 2012)). We note that k-mer approaches to GWAS have been performed, but still suffer from the majority of issues above: for example, GWAS of ampicillin resistance in E. coli identified SNP variants in a known drug target but functional assignment of variants beyond SNPs were not discussed (Rahman et al. 2018); similar studies were later conducted in plants (Voichek and Weigel 2020). Empirically, GWAS has only enjoyed partial success for bacteria and predictive power, and methods that can provide pan-bacterial, reproducibly strong, interpretable prediction of resistance remain out of reach (Wiatrak et al. 2024). To our knowledge, GWAS has not been successful in fungi or viruses.

Massive experimental or parallel genetic screens can be used to interrogate genotype-phenotype relationships as an alternative to GWAS, but can be limited in measuring phenotypes in genetically tractable systems and or systems that are manipulatable in the laboratory– bioethical concerns, especially around gain of function, limit the feasibility and or ethics of executing these experiments. Existing computational pipelines often require knowledge of gene annotations to assess the importance of a given prediction. These pipelines can often have highly customized statistical, computational and experimental steps, or all of the above.

Deep learning approaches also have limitations; they can learn representations in the data that can be used for zero-shot prediction and for fine tuning (Wei et al. 2021), but they are computationally intensive and do not readily generalize between domains of life (e.g. between microbes and eukaryotes) and do not reliably re-identify canonical resistance markers (Hu et al. 2024). Further, they are trained on reference genomes; therefore, they necessarily cannot attribute phenotype to missing sequences. Sequences missing from these genomes are “out of distribution” (Yang et al. 2021) meaning functions for them are poorly predicted if at all. Further, it remains a major open question in the field whether deep learning models indeed improve inference over more classical machine learning analyses (Ching et al. 2018; Sapoval et al. 2022; Wysocka et al. 2023). To our knowledge, no deep learning approach takes as input raw reads and produces phenotype predictions with sequence attribution (i.e. feature importance at nucleotide resolution).

In principle, direct inference of both phenotypes and of the genetic variants which cause them should be possible from raw sequencing reads. Genetic data in nature samples multitudes of sequence variants and explores a highly rich space of genetic changes representing repeated selection on potentially highly divergent genomes. This process creates diverse genetic variation where causal interactions will be consistent, whereas variants unrelated to the phenotype under selection will wash out. Millions of sequenced isolates exist from model and nonmodel organisms, including viruses, bacteria and fungi—all of great importance to public health (e.g. pathogen adaptation and resistance to antimicrobial drugs). Not all of these species have reference genomes and many can have highly plastic, rapidly evolving DNA. In principle, the signal in these natural sequences should reveal mechanisms of resistance.

Here, we show the power of this idea, introducing a new paradigm for jointly linking sequence and phenotype called *FunctionaL Assigning Sequence Homing* (FLASH). FLASH takes raw sequencing data, bypassing genome assemblies and alignment to identify sequence features driving biological phenotypes. Phenotype is abstract; examples include, but are not limited to lineage, resistance phenotypes, viral host range or cell type. Sequence is also abstract: a cluster of sequences within a specified edit distance (rigorously defined) that could be considered as an analogue of a multiple sequence alignment. FLASH dually predicts significant and functionally important biological sequence clusters and uses them to develop a model of how their variation—or sequence change—modifies a phenotype. It can be seen as identifying genotype from phenotype, flipping the conventional approach.

FLASH identifies parsimonious sets of sequence variants—which we refer to as features or clusters because they can be thought of as clusters of gene-like sequence—that predict a phenotype. By design, FLASH predicts functional attributes correlated with variation in a sequence. The simplest example is a mutation that confers resistance, but this variation is much more general, and can include many mutations, as well as insertions, deletions or structural variants. FLASH takes as input the output of our previously developed reference-free, statistical feature extractor SPLASH (Chaung et al. 2023; Kokot et al. 2024; Baharav et al. 2024; Henderson et al. 2024; Meyer et al. 2024; Dehghannasiri et al. 2024, 2025), but there is no explicit dependence or requirement that SPLASH be used. A significant limitation of SPLASH is that it does not predict which of the potentially millions of k-mers it identifies contribute to phenotypic difference. We developed FLASH to fill this gap.

FLASH is very simple to run, modular, and highly efficient. It completely bypasses reference genomes or alignment and achieves blind ‘universal’ phenotypic prediction accuracy that matches or exceeds custom bespoke studies that we chose based on the existence of metadata from public data. We analyzed raw sequencing data from >35,000 isolates from 9 different studies of bacteria, 4 studies of fungi, and 1 study of H5N1, an RNA virus (each study processed from raw reads to predictions and annotations completed in <24 hours including studies with thousands of isolates) to distill interpretable features that predict phenotype.

Microbes have polygenic phenotypes, and predicting them from genomic sequencing can be difficult, partly due to their complex and highly plastic genomes (Braz et al. 2020; Sakoparnig et al. 2021). Canonical targets of widely-used antimicrobials (eg. macrolides, or beta lactams) are known for many drugs. However, resistance to even the oldest—such as penicillin—can be due to multiple mechanisms beyond target modification (Elshobary et al. 2025). Methods to identify genes synergistic to canonical resistance are urgently needed, both to predict treatment response and design new drugs. Further, factors driving virulence are similarly difficult to predict, such as group B Streptococcus, where stratifying patients and monitoring disease outbreaks are essential. Fungal pathogens are emerging and evolving rapidly and have been understudied, and thus present similar problems. Finally, as new therapies such as phage therapy emerge, computational methods are needed to predict molecular interactions that include phage host range.

Unlike black box deep learning or many machine learning techniques, FLASH provides transparent, interpretable results, assigning a phenotype prediction based on embedding sequences of a parsimonious set of features chosen by the model. We present the results of running FLASH with a pipeline and parameter regime, blind to any sequence metadata in eukaryotes, prokaryotes and virus. FLASH predicts multiple antibiotic resistance phenotypes in diverse species of bacteria, drug susceptibility in fungi, and host range in the avium influenza virus H5N1, all without previous knowledge of the input data or a reference genome. Further, FLASH associates coding and noncoding sequences of unknown function to clinical resistance phenotypes, and identifies new sequence variation missing from the available NCBI reference genomes, including in a fungal gene where we identify structural plasticity (highly elevated polymorphisms due to copy number changes, or repeat expansion and contraction) associated with virulence. We show that FLASH can be trained on raw read data and used to predict on different data inputs such as previously assembled genomes. Finally, FLASH performs tasks that, to our knowledge, are not available by any method: it predicts host range of phage, and can nominate new pan-species predictors of resistance in bacteria. FLASH is available publicly at <u>github.com/djcotter/FLASH-smk/</u>.

## Results

FLASH is an interpretable, scalable machine and deep-learning method that operates on k-mer (nucleotide sequences of length k) representations of each sample to jointly analyze sample phenotypes and genetic features (Fig. 1). It has three key steps. First, it performs “clustering” to identify similar k-mer sequences, called anchors, which precede significant downstream variation. Second, FLASH identifies the most abundant, or representative, sequence variant per sample per k-mer cluster, which is used for prediction; if no such sequence exists, it is encoded as “N..N”. Third, FLASH maps these variants for each sample to numbers via embedding each sequence in our pre-trained nucleotide language model (Methods). FLASH uses this sample representation to perform a) unsupervised clustering on the underlying structure of the data or b) supervised dual prediction of phenotypes and their features, fitting models on a balanced training set, the FLASH analogue of fine tuning. FLASH computes accuracy on at least 20% of held out test data (Methods).

**Figure 1A.**
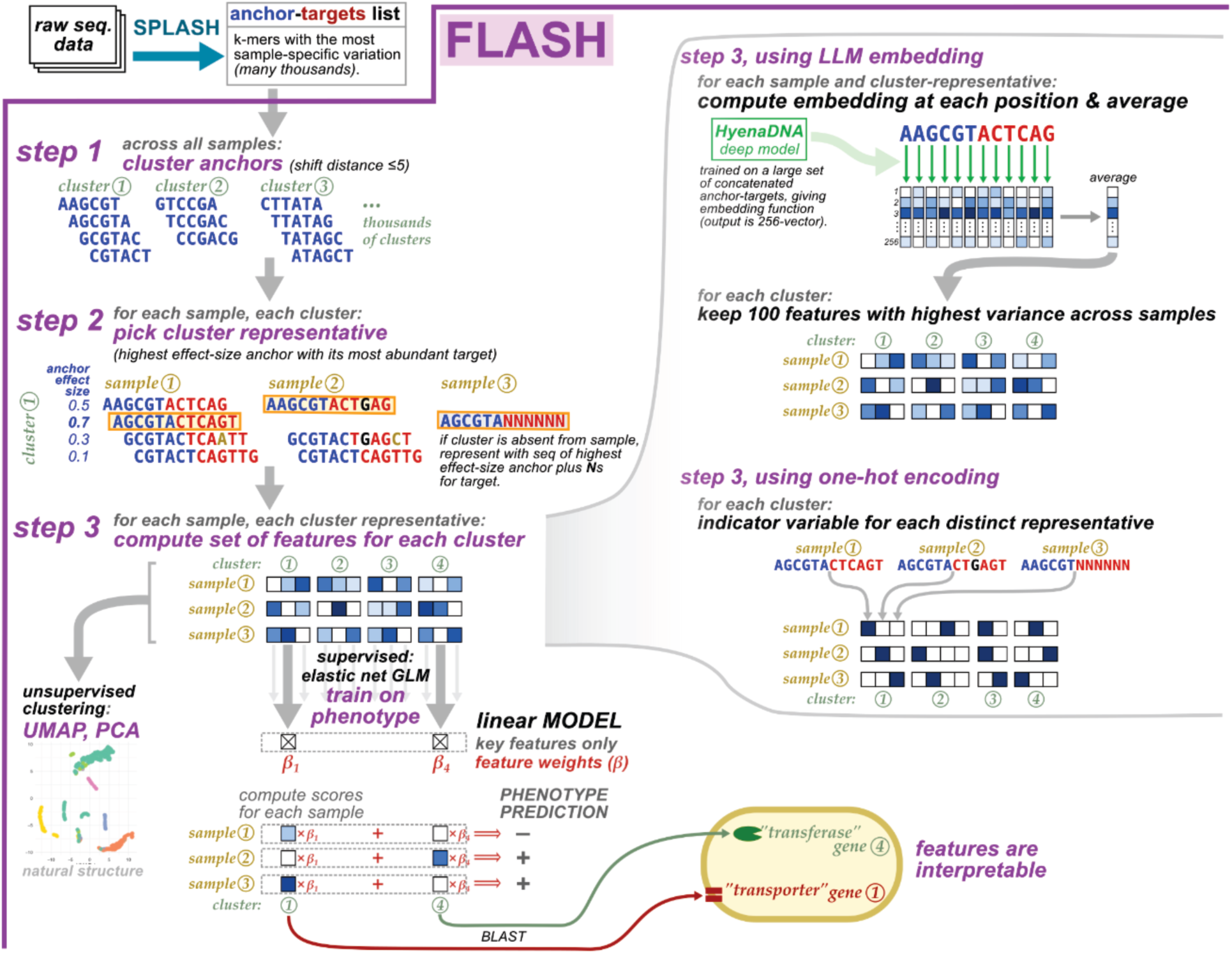
Schematic of FLASH. Includes running SPLASH to featurize inputs, followed by step 1: clustering of anchors; step 2: parsing each sample data into a representation of the clusters via the most abundant target for each anchor; step 3: training a supervised model to predict phenotype and dually identifying clusters that drive that prediction, or unsupervised clustering on the most variable embeddings. Right: Step 3 can take the form of either embedding sequences in a language model and averaging the embedding dimensions to use as predictors (right, top) or by one-hot encoding the presence of each feature (right, bottom).

For interpretation, clusters chosen by the FLASH model are assigned gene identity via protein and nucleotide homology searches with BLAST (Methods). Variants in a cluster may have no hits to 0, 1 or many entries in these databases and may or may not be annotated; their matches may or may not be perfect. When some sequences within clusters map to a reference but others diverge substantially from a reference, these sequences may be de-orphanized through their link to a cluster and an associated phenotype. This is a key to the interpretability of FLASH, and to providing prediction of how genetic variation impacts phenotype. These annotation results are output as both figures and summary tables (Fig. 1B).

**Figure 1B.**
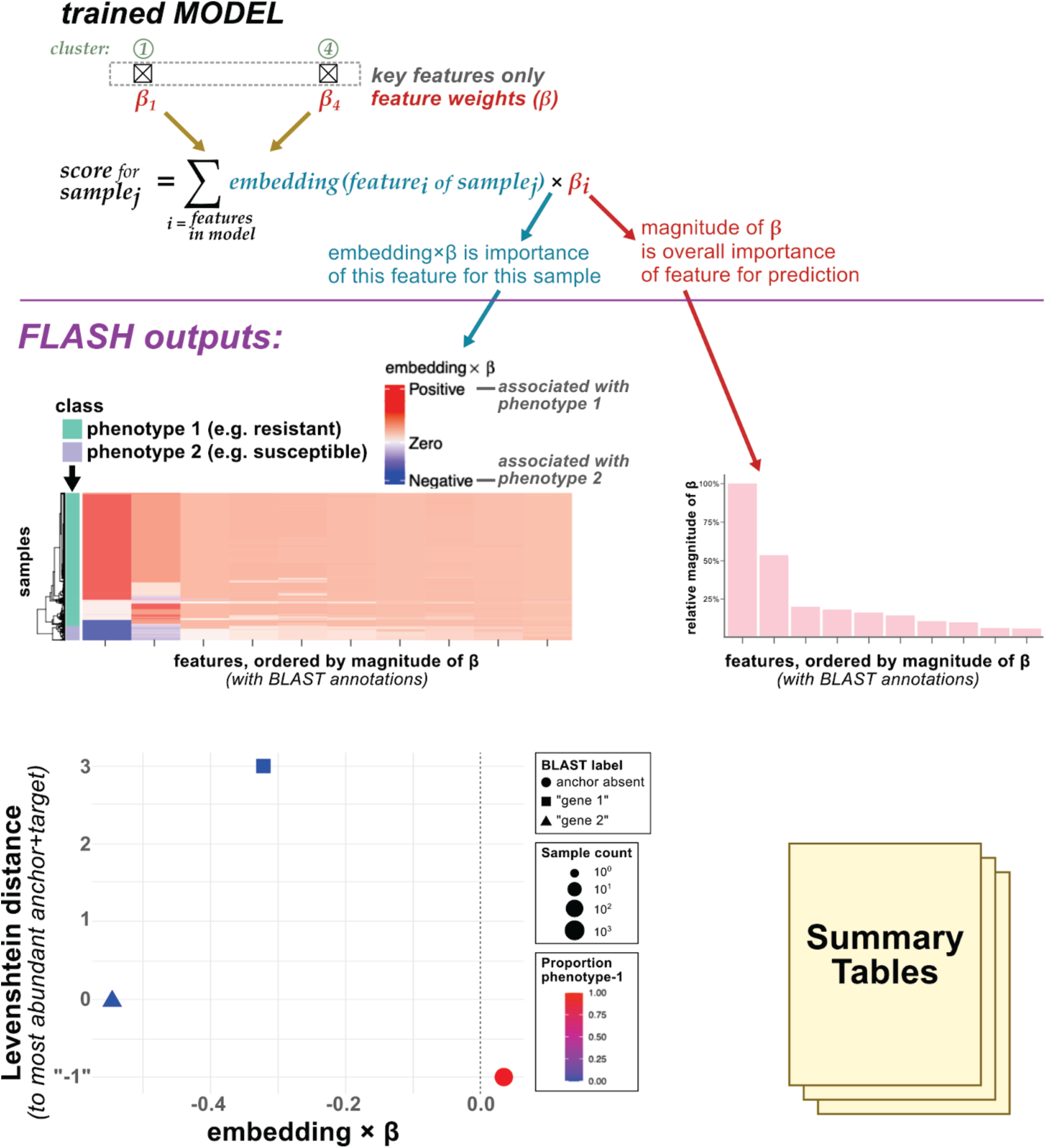
FLASH produces interpretative outputs for the user. FLASH pipeline takes input from the user and produces graphical outputs including heatmaps of per-sample, per-cluster importance or annotations and graphical displays of how sequences vary per metadata category. Heatmaps illustrate the effect that the embeddings have on the prediction of the model with metadata annotated via color to the left. Barplots illustrate the magnitude of the effect of each nonzero coefficient on the model. Scatter plots show visually each separate variant in a cluster. “-1” indicates an anchor is absent. Labels indicating the Identity (I) and coverage (C) of each variant to a BLAST hit appear next to points with imperfect hits. Summary tables containing BLAST hits, prediction accuracies, and the magnitude of each nonzero coefficient cluster on the prediction are also reported.

A user has many parameter tuning options to restrict input features in multiple ways such as by changing the cluster assignment or removing features that are missing across some number of samples in the dataset, should they desire. For example, there are 64% of features that are missing in at least 20% of samples in *E. faecium*. Our results here were based on one parameter regime, and a kmer size which we did not optimize (see methods). FLASH can be run on a variety of datasets ranging from bacterial to fungi to viruses. We explore datasets of each type here (Fig. 1C; Supp. Table S1).

**Figure 1C.**
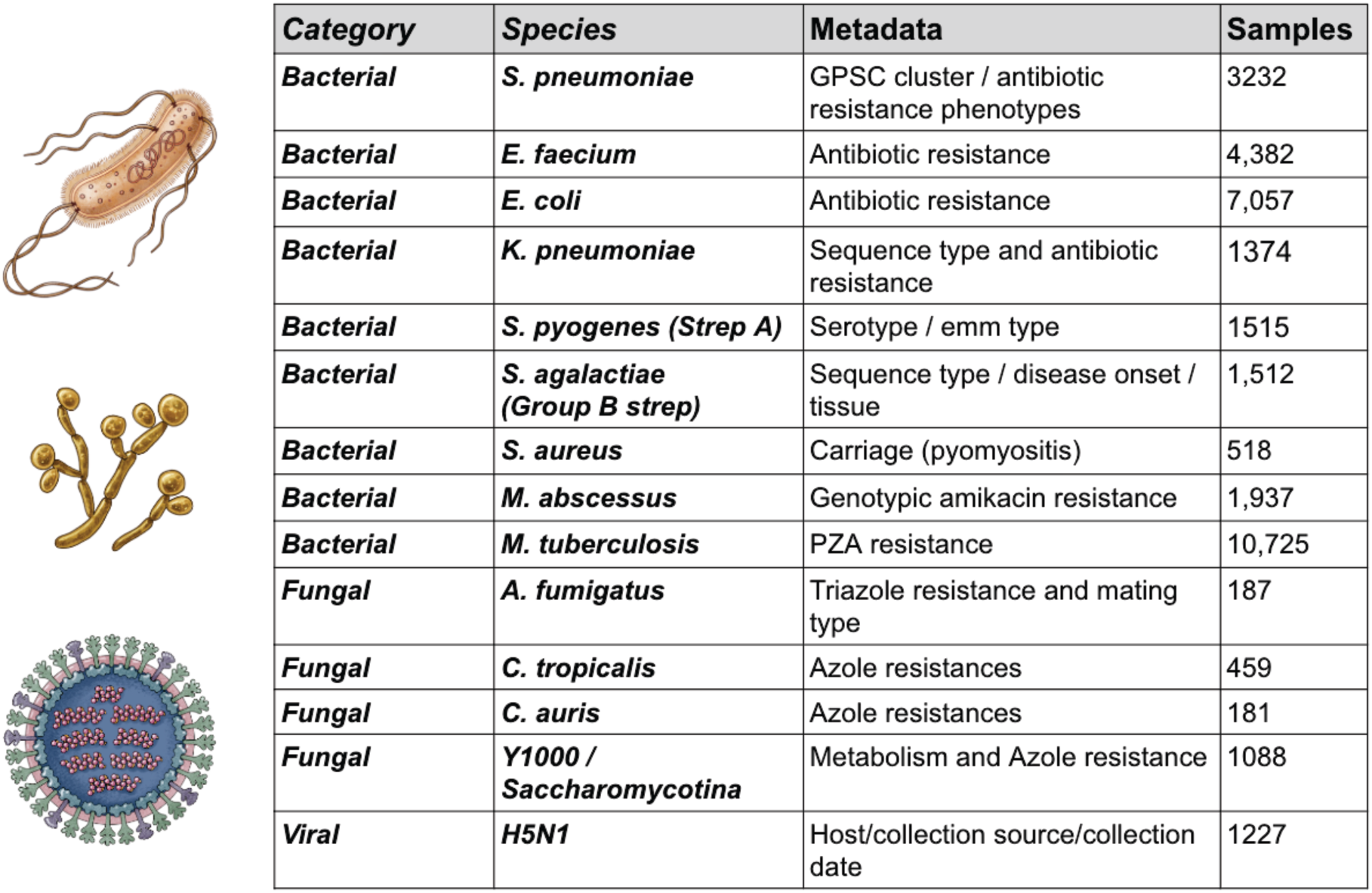
Size of data and scope of metadata. Datasets, species and number of isolates analyzed in this study.

### A unified FLASH model predicts resistance phenotypes as well or better than bespoke methods

FLASH runs agnostic to input data, and therefore without species-specific tuning. In our tests, it meets or exceeds accuracy of bespoke benchmarking studies, many reporting accuracies on data used in training. In 4,382 isolates of *E. faecium*, FLASH predicts antimicrobial phenotypes with equal or higher accuracy to a state of the art pipeline (Coll et al. 2024) (Fig. 2A, left, Supp. Table S2). The accuracy of FLASH predictions is 3.4% increased on average compared to the original study. In 1051 fungal species across the *Sacchoromycotina* subphylum, FLASH also predicts antifungal phenotypes with a mean increase in accuracy of 17% compared to the original study based on recent machine-learning predictions tailored to fungal lineages, whereas FLASH is agnostic to them (Harrison et al. 2025) (Fig. 2A, right). FLASH accuracy on blind test data of *M. tuberculosis* was 92% compared to 79.5% in a recent study when predicting pyrazinamide resistance ((Carter et al. 2024); Fig. 3). FLASH is also highly efficient and easy to use: one pipeline command analyzes this set of ∼10,000 *M. tuberculosis* samples (Kim et al. 2023) from raw read input in 4.1 hours (Supp. Table S3).

**Figure 2A.**
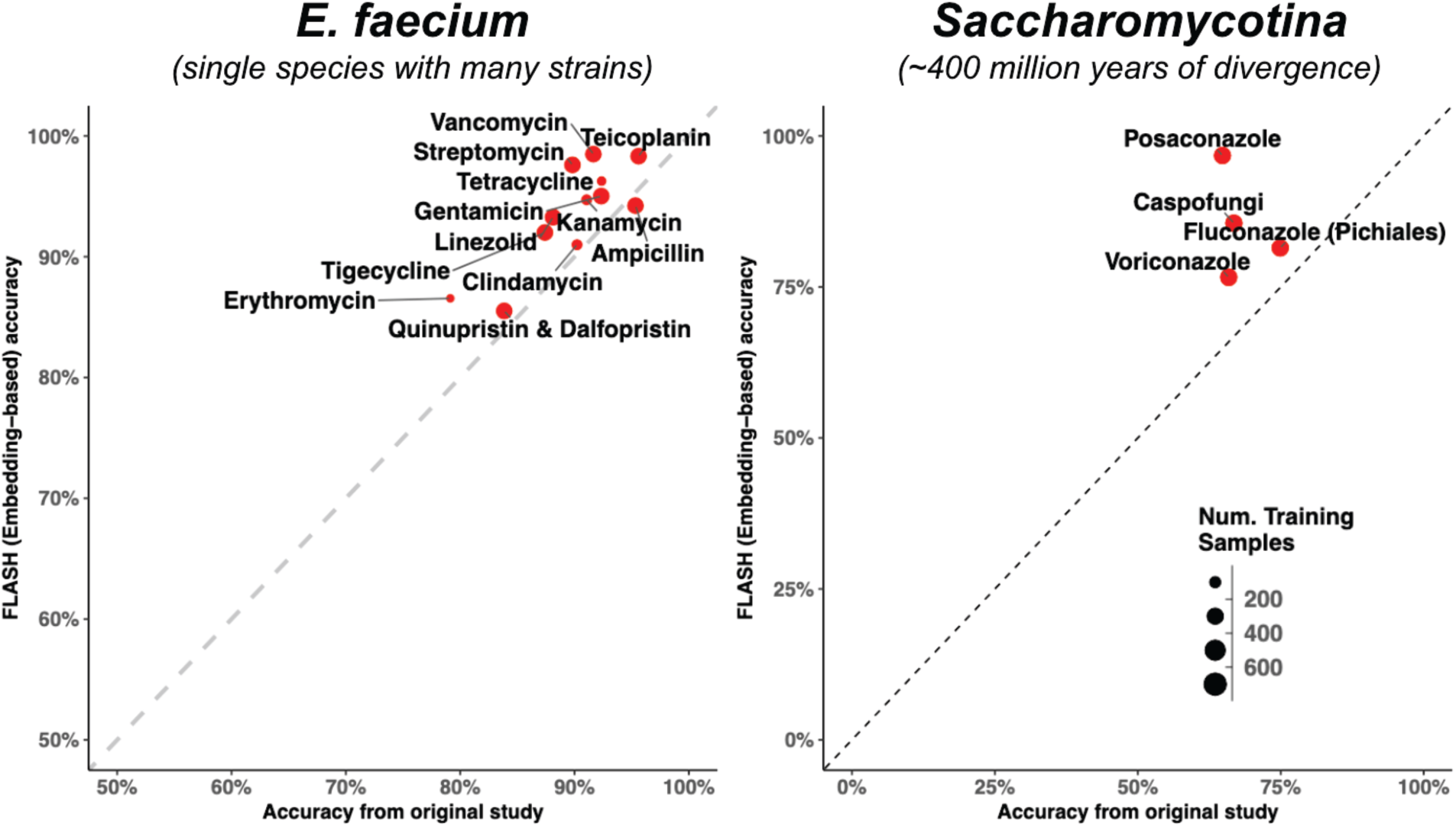
FLASH accuracy performs comparably to bespoke studies predicting phenotype both in individual species and across 400 million years of evolutionary history. Accuracy reported for *E. faecium* (left) calculated using embeddings on a model trained on balanced classes composed of 80% of the smallest class. Test accuracies are compared to accuracies calculated by the original paper which we believe to be based on training data, hence a conservative comparison to blind test accuracy (Coll et al. 2024). FLASH accuracies have, on average, a 3.4% mean increase from the original study. Saccharomycotina accuracy (right) is reported for predictions made for Y1000 (Harrison et. al., 2025) data using blind test data fitting a model based on nucleotide embeddings. Antifungal resistance predictions have a mean increase of 17.0% using FLASH when compared to the original study. The size of the data points reflects the number of samples used to train each model. We exclude datasets with less than 200 samples due to variances in estimates of the accuracy.

### FLASH obtains high zero-shot accuracy in never-before-seen datasets and sequences, including assembled genomes

FLASH predictions are functions of sequence embeddings in a custom-trained model (Methods, Supp, Table S4). Accuracy with embeddings exceeds that of One Hot Encoding (OHE) (Fig. 2B&C). Beyond improved accuracy, embeddings enable FLASH to generalize GWAS, as the model can be used to perform prediction on sequences never seen in training. To demonstrate this, we deliberately trained the *E. faecium* model on a single study to constrict the amount of target variation seen during training. We trained the model on only isolates from this study, and tested on the remaining studies. A substantial fraction, ∼25%, of features per cluster were never seen by the model during training. Yet, accuracy on the held out test data with the remaining targets did not change (Fig. 2D).

**Fig 2B:**
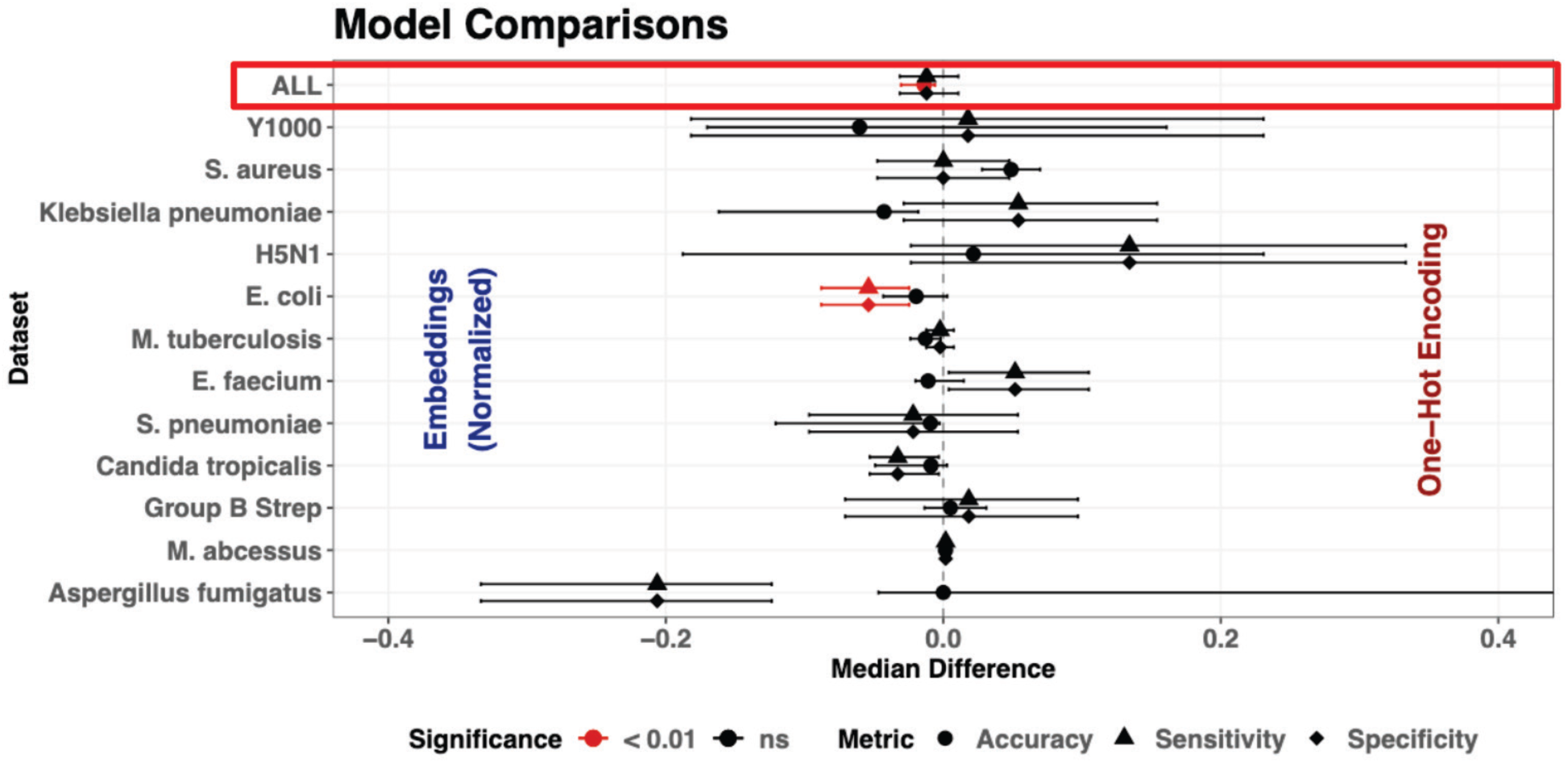
Embedding models perform better than one hot encoding on average. Comparison of the median difference in accuracies using a two-sided Wilcoxon signed rank test between the nucleotide-embeddings-based models and the One-hot-encoded (OHE) models only comparing datasets with binary classes via a Wilcoxon Rank-sum test. Medians are reported as the pseudomedian with appropriate confidence intervals. Across most datasets, accuracy (indicated by the circle) for OHE predictions is not significantly different from accuracy for embedding-based predictions. Summing across all datasets (red box), the embedding-based model has 0.9% higher median accuracy than the OHE model (p<0.01). Strep A is excluded as we exclude all phenotypes from this plot with > 2 classes to better compare accuracies across datasets. Wide error bars exist in datasets with fewer samples where p-values cannot be accurately calculated.

**Figure 2C.**
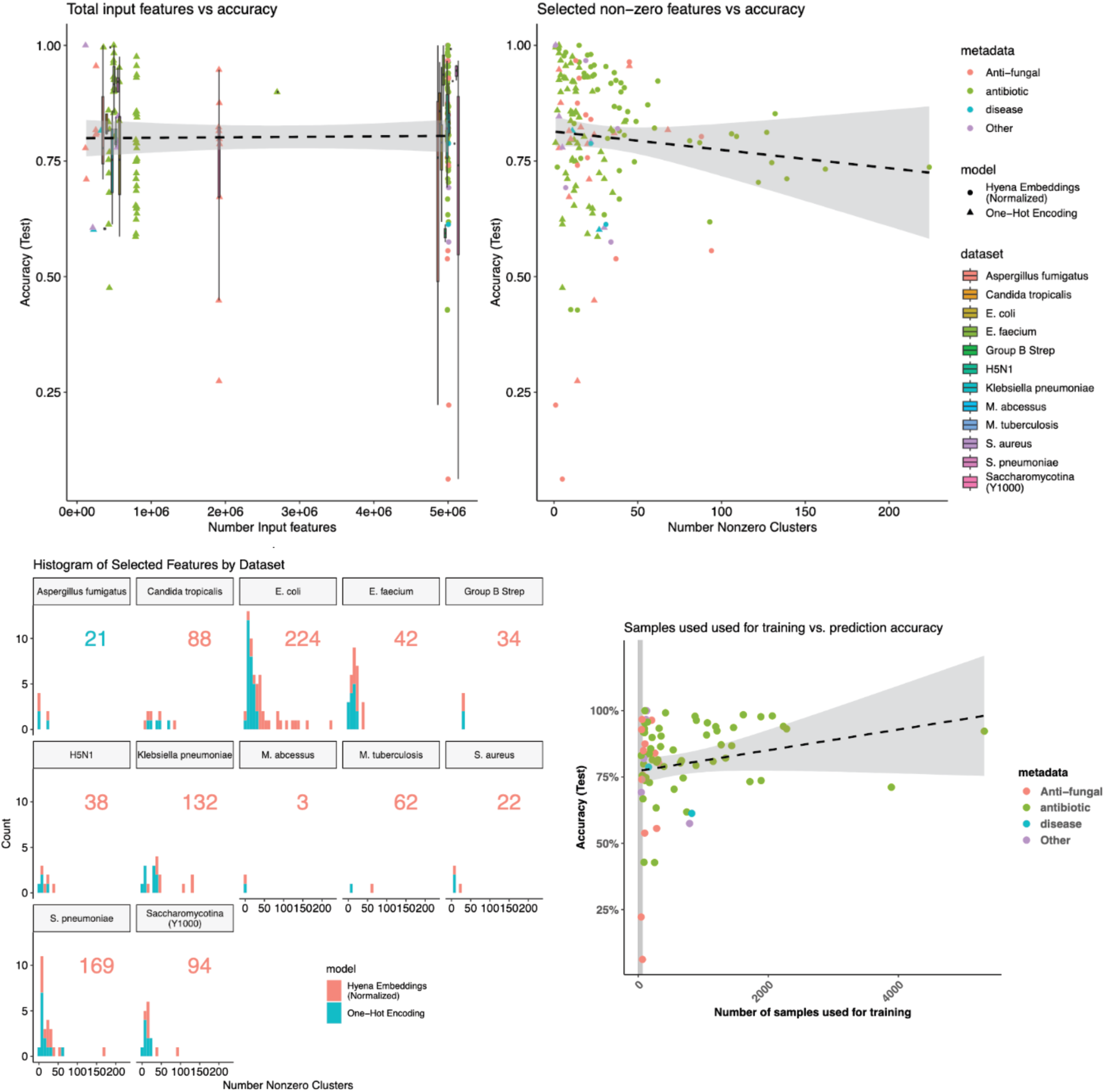
Number of input features and number of selected features do not affect accuracy. Using only accuracies calculated for metadata categories with binary phenotypes (on the y axis), accuracy marginally increased with the number of nonzero features selected by the model (top). Most models and most datasets have comparable numbers of selected features. Bottom left is the histogram of the number of selected features for the prediction by model for each dataset. The maximum number of features selected for any of the metadata categories is indicated in the upper right corner of each plot. Very low prediction accuracies <25% are in models trained on very few samples (bottom right). An increase in accuracy is associated with an increase in the number of samples used for training, both for binary phenotypes and phenotypes with greater than two classes. The grey box indicates tests with few samples <=50. The regression line does not include these accuracies.

**Fig 2D.**
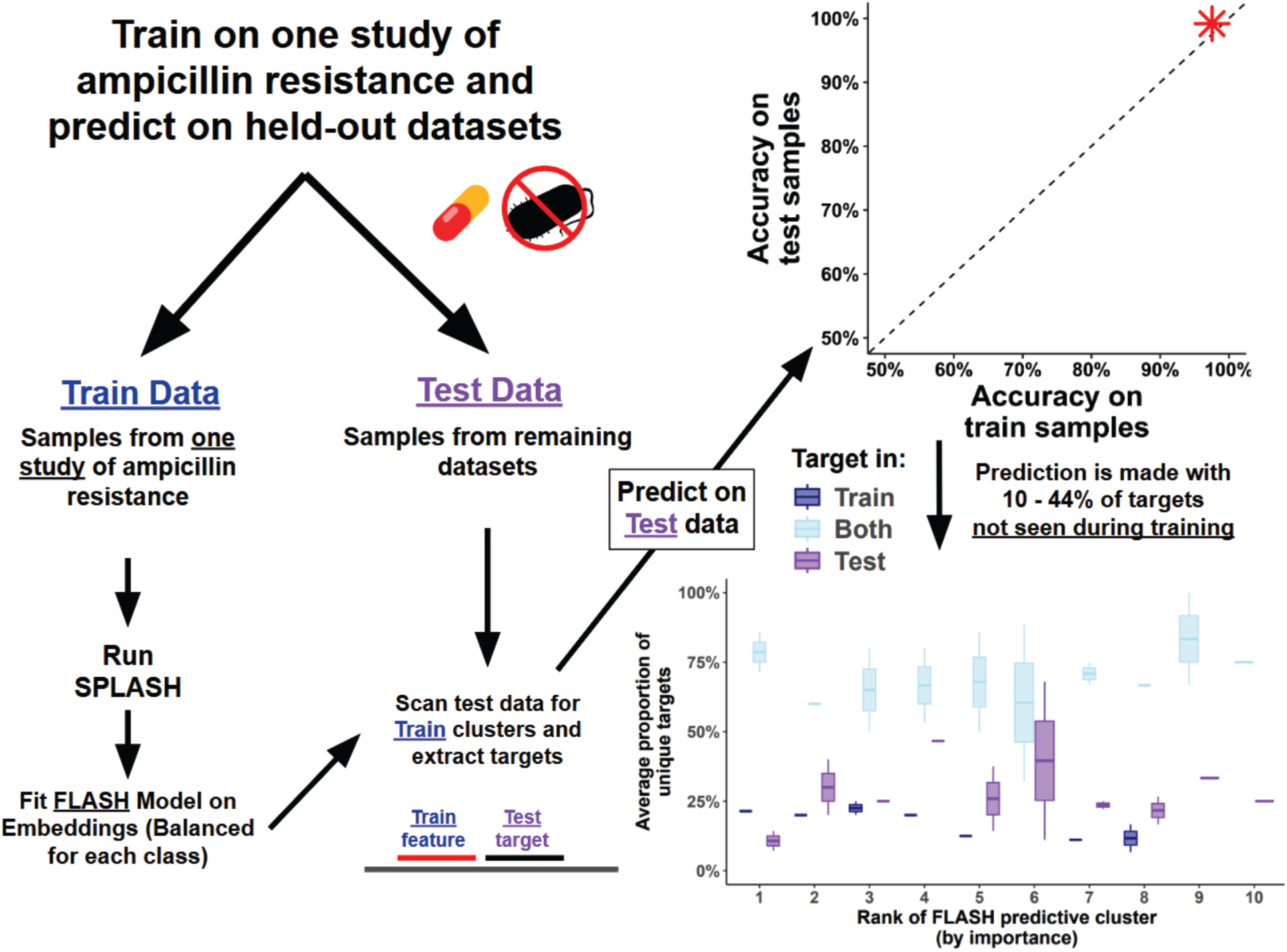
Generality of FLASH. FLASH models predict on never-before-seen variation after artificially splitting the test and train data. FLASH predictions can be made in held-out sets of samples including other raw fastqs (left) where we subset the entire *E. faecium* dataset to a train and test set based on the selecting one study from a list of studies that were compiled of *E. faecium* by (Coll et al. 2024). SPLASH is run first only on the training data to discover anchors that are best for this set only. Then the FLASH model is trained. Accuracy (red asterisk; top right) is reported on held out samples from the initial trained model (x-axis) and tested on the remaining samples from the other studies which never had SPLASH run on them (y-axis). 10% to 40% of the targets used for prediction in the top 10 predictive clusters were never seen by FLASH prior to prediction (Bottom right (ampicillin resistance)).

We further tested if FLASH models could predict experimentally measured resistance phenotypes on assembled genomes (Hyun et al. 2023), including in *E. faecium* (∼1300 genomes) and *E. coli* (∼2600 genomes) (Fig. 2E). Accuracy was comparable to test accuries on raw reads and correlated strongly (adj r2=0.95). On average 1 (*E. faecium*) to 1.5 (*E. coli*) unique targets per feature cluster were never seen in the training data (Methods), again evidence the model generalizes prediction. In FLASH models trained on Illumina data from a different groupa predicted vancomycin-resistance of isolates profiled with long Nanopore reads with 100% accuracy (Islam et al. 2023). This suggests that the use of deep learning embeddings and models trained on robust short-read data could be extendable for prediction directly in the clinic where real-time sequencing is in increasing use (Supp. File S1).

**Fig 2E.**
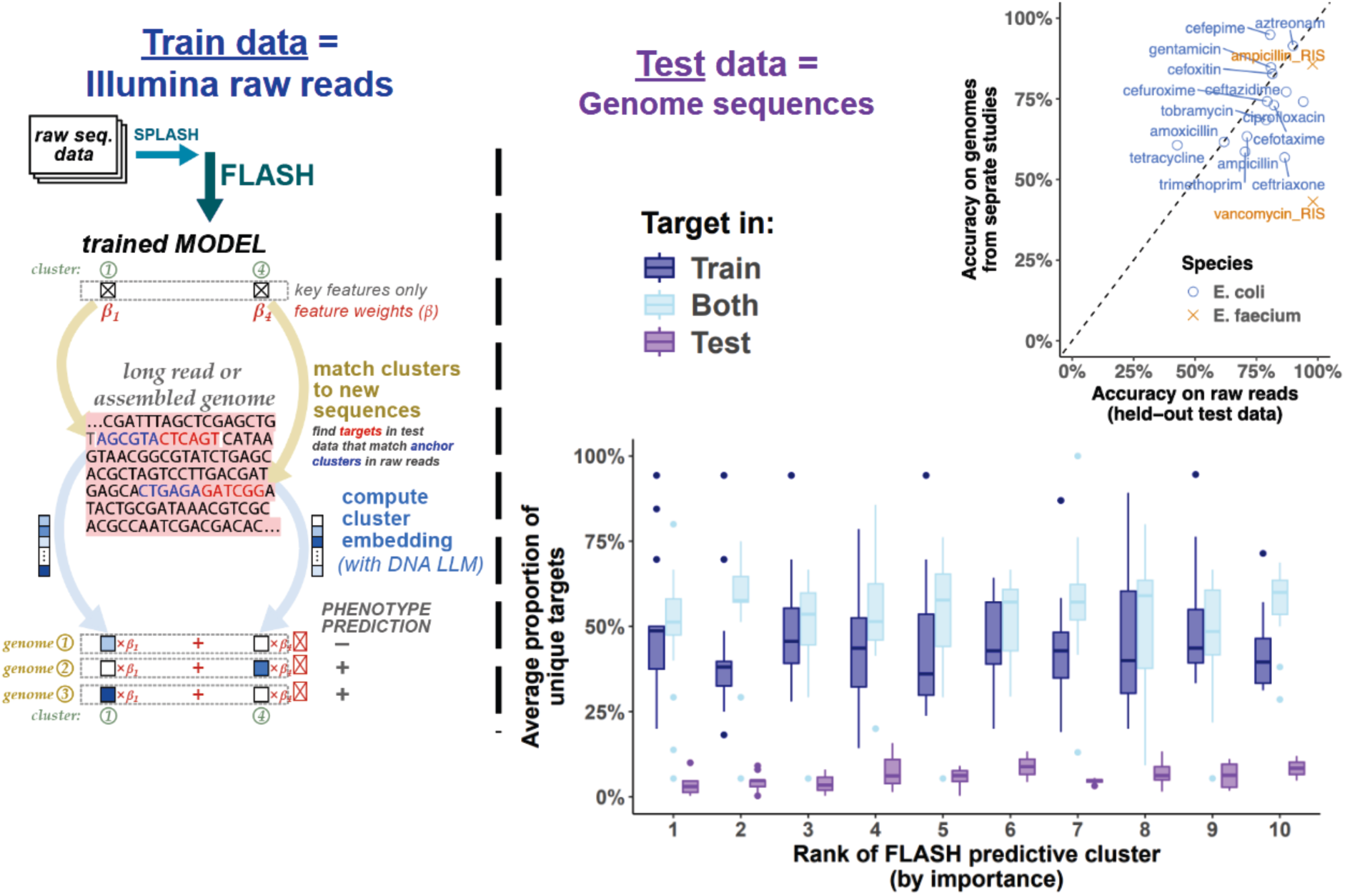
Generality of FLASH. FLASH models predict on assembled genomes of isolates not seen by training data. FLASH predictions can be made on held out sets composed of completely different data types. FLASH is fit on short Illumina raw reads splitting the data in half and predicting on raw reads using a held out test set. Then, genomes are used as an additional held-out test set by sliding along their length and grabbing sequence variation associated with the anchor clusters used to fit the original model, extracting overlapping clusters with targets never-before-seen by FLASH. In *E. coli* (blue) and *E. faecium* (orange), accuracies in genomes are comparable to accuracies in the raw read model (adjusted R^2 = 0.95; accuracy here for held out test data). 3.3% to 8.8% of the targets used for prediction in the top 10 predictive clusters were never seen by FLASH when training the model (box plots, purple boxes).

### FLASH detects functional genotypes across multiple different bacterial species

In its unsupervised mode, FLASH predicts and identifies existing structure in various microbial communities. In both *S. agalactiae* and *S. pyogenes* it recaptures patterns of serotype and Emm type (Fig. 3A). Across the cases we tested, this clustering suggests that FLASH is truly picking up on the underlying variation represented in these species. We went on to further examine which features FLASH discovers in its supervised mode.

**Fig 3A.**
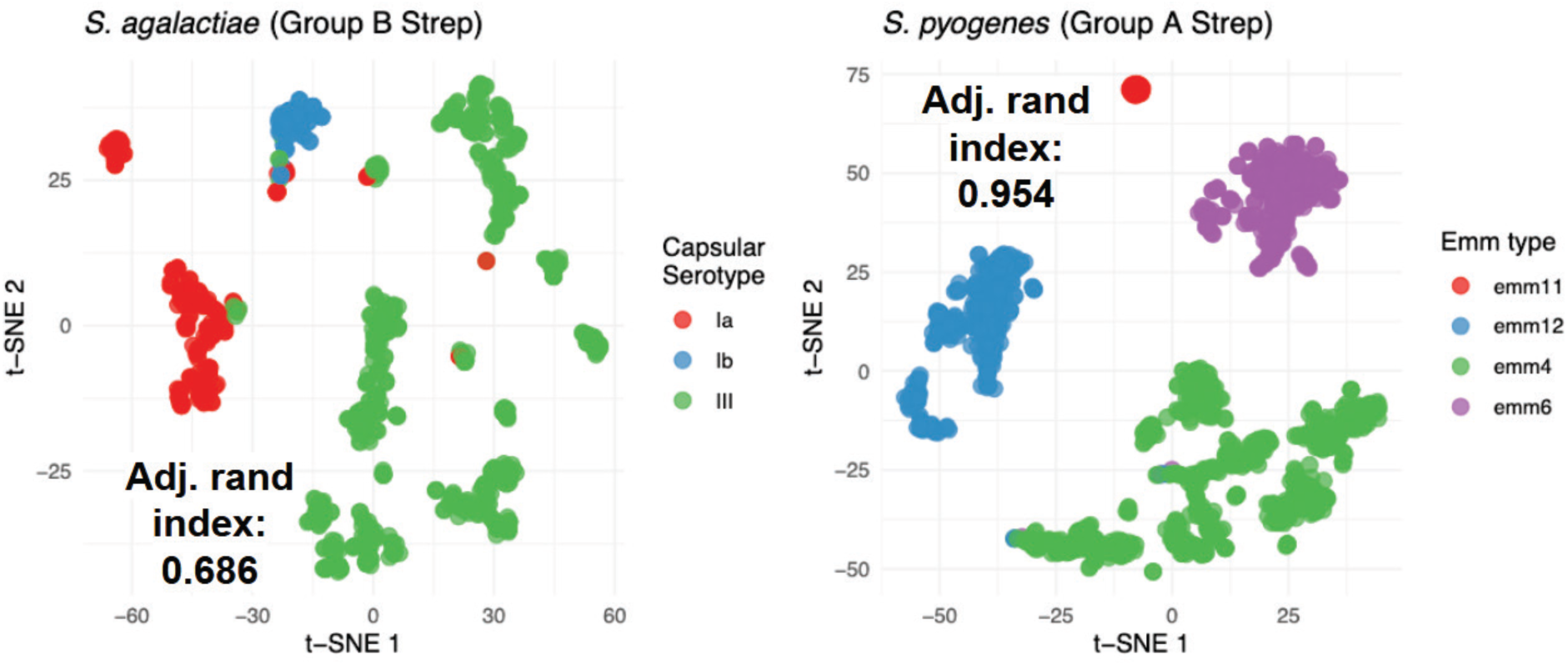
Unsupervised clustering rediscovers common sequence clusters. t-SNE clustering of sequence variation in 1,338 samples of Group B strep (or *S. agalactiae*; left) and 1,575 samples of Strep A (or *S. pyogenes*; right). Sequence variation is identified using SPLASH before these features are clustered and embedded (see methods). Using the matrix composed of samples (with classes that have >=30 samples) by one embedding dimension per cluster. We first performed principal component analysis on the embedding matrix and then visualized the top 10 principal components with t-SNE. Adjusted rand indices for the clustering shows that FLASH recaptures serotype and Emm type (0.686 for Group B strep and 0.954 for Group A strep).

Mycobacterium tuberculosis is one of the best studied microorganisms, and clinical isolates have well-known resistance mechanisms. It also has a very stable genome characterized by a lack of horizontal gene transfer and interstrain recombination (Gagneux 2018). We tested if FLASH could identify the target of pyrazinamide (Whitfield et al. 2015), a medication commonly given after an isolate displays resistance to first-line anti-TB drugs (Supp. File S2). Resistance to PZA was predicted with accuracy > 92.2%, higher than 79.5%, reported in a recent study of pyrazinamide resistance prediction (Carter et al. 2024).The three highest predictors of resistance were rpoB and katG, both known resistance genes in *M. tuberculosis* (Torres et al. 2015; Jose Vadakunnel et al. 2025). The fourth, the known direct target of the drug, is the gene pyrazinamidase (PncA). Many targets used for prediction had significant sequence variation or were missing, types of variants that would likely be missed in a GWAS. This is consistent with FLASH identifying the implied obligatory resistance acquired by resistant strains that were exposed to other first-line antibiotics, and that FLASH features include the direct drug target supporting its ability to detect direct resistance mechanisms.

We additionally tested FLASH prediction accuracy on a large and diverse set of ∼1900 isolates of *Mycobacterium abscessus* by mining multiple studies in the short read archive and annotating resistance based on known mutations in the 16S rRNA. FLASH predicted amikacin resistance with 99% accuracy, re-identifying this canonical target of amikacin without prompting. To explore correlated features associated with multidrug resistance, we removed any cluster in the 16S rRNA (achieving prediction accuracy of 96%; Supp. File S2). The top hits were enriched for associations with drug resistance and virulence. These included hits to MarR, TetR, and two different type VII secretion proteins (Bar-Oz et al. 2022; Rodriguez et al. 2023). This approach underlines the ability of FLASH to identify genes synergistic with one resistance phenotype, either due to a clinical history of exposure, or a general virulence phenotype.

In *E. faecium* and *S. pneumoniae,* the top predictors of ampicillin and penicillin resistance (respectively) include penicillin binding proteins– with target variation in *E. faecium* mapping to the C-terminal region of the gene in the binding pocket of penicillin–effectively re-discovering this pocket ab initio (Fig. 3B, left). In *S. pneumoniae,* the top hit is in the penicillin binding pocket penicillin-binding protein 2B. While these genes are well-established in the field as correlated with ampicillin and beta-lactam resistance (Hunashal et al. 2023), they are not sufficient (Fig. 3B). In *E. faecium* and *S. pneumoniae,* FLASH accuracy is driven by other predictors, including various IS transposeases, transporters and hits to other known antibiotic resistance genes (Supp. Files S2&S3). In the aminoglycoside gentamicin, FLASH’s driving feature is GNAT family N−acetyltransferase/aminoglycoside phosphotransferase (APH), a known driver of resistance (Fig. 3C) (Garneau-Tsodikova and Labby 2016) (Supp. File S3). Likewise, for vancomycin, 6 of the top 10 predictors are to various vancomycin resistance genes (Fig. 3C, right). In both cases these predictions are driven by the absence of the target variation in the susceptible samples. Among others, FLASH identifies gyrA for ciprofloxacin resistant *E. coli* (94% accuracy), Erm(B) for erythromycin resistant *S. pneumoniae* (95% accuracy), and Tet(M) for tetracycline resistant E. faecium (96% accuracy). These are all known drivers of the corresponding resistance phenotypes (Supp. Files S2&S3).

**Fig 3B.**
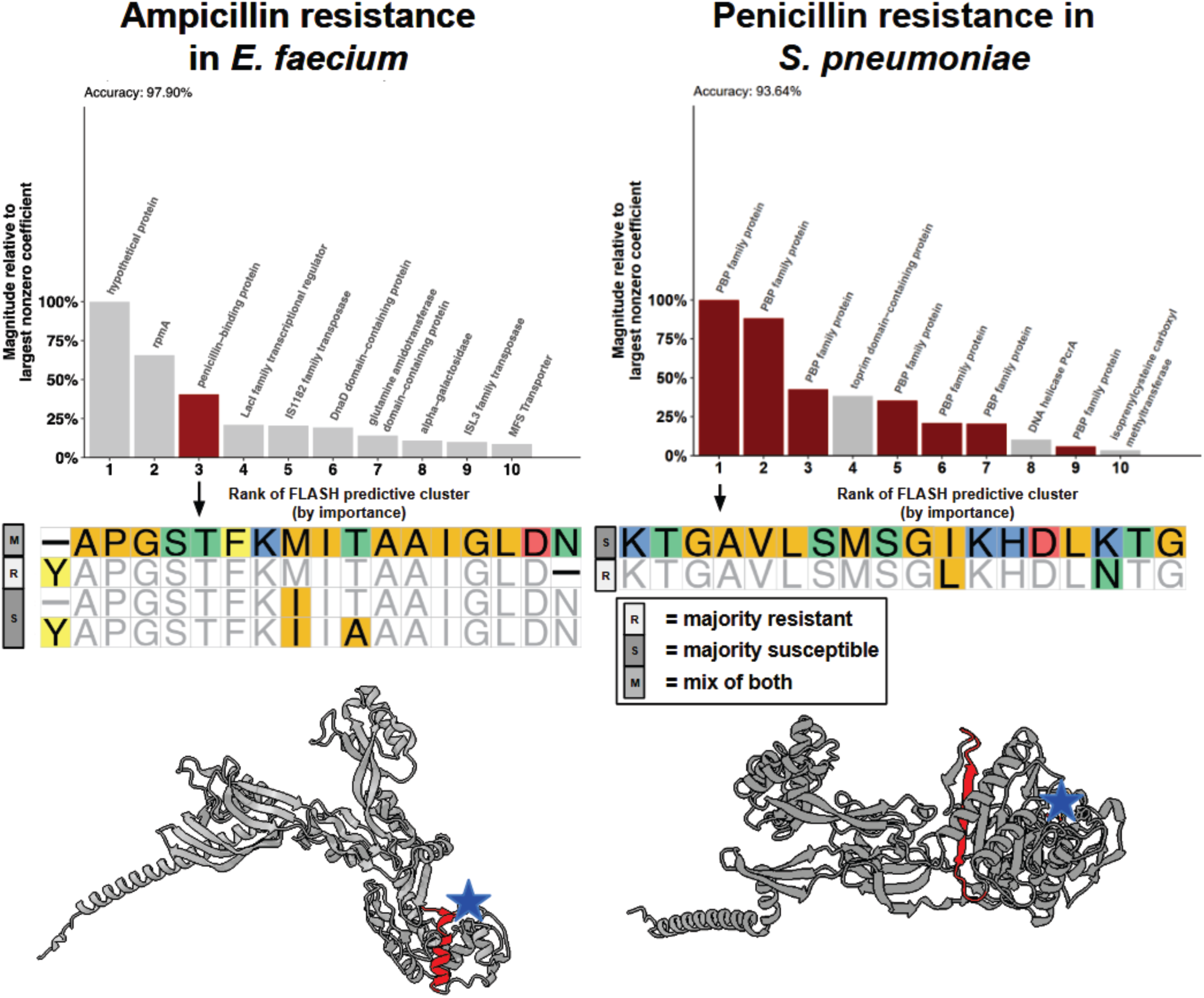
Penicillin binding proteins rank highly in predictions of beta-lactam resistance in *E. faecium* and *S. pneumoniae*. Penicillin binding protein has some of the highest nonzero coefficients in predicting ampicillin resistance in *E faecium* (left) and *S. pneumoniae* (right). The target variation is represented, translated into amino acids, for the most abundant variants (AA in majority resistant samples indicated by R, AA in majority susceptible samples indicated by S, AA with a mix of resistant and susceptible samples indicated by M). The structure of PBP5 on the left (A0A1L8WSI8) includes the annotated target region (red) and a star indicating the binding pocket of penicillin (blue). On the right, penicillin resistance is driven by variation in two different penicillin binding proteins (6 hits to Pbp2b and 1 to Pbp1a) with multiple FLASH hits in different domains the first feature illustrated on the protein structure A0A0H2ZQ75 (Red indicated the target region and the blue star indicates the binding pocket of penicillin).

**Fig 3C.**
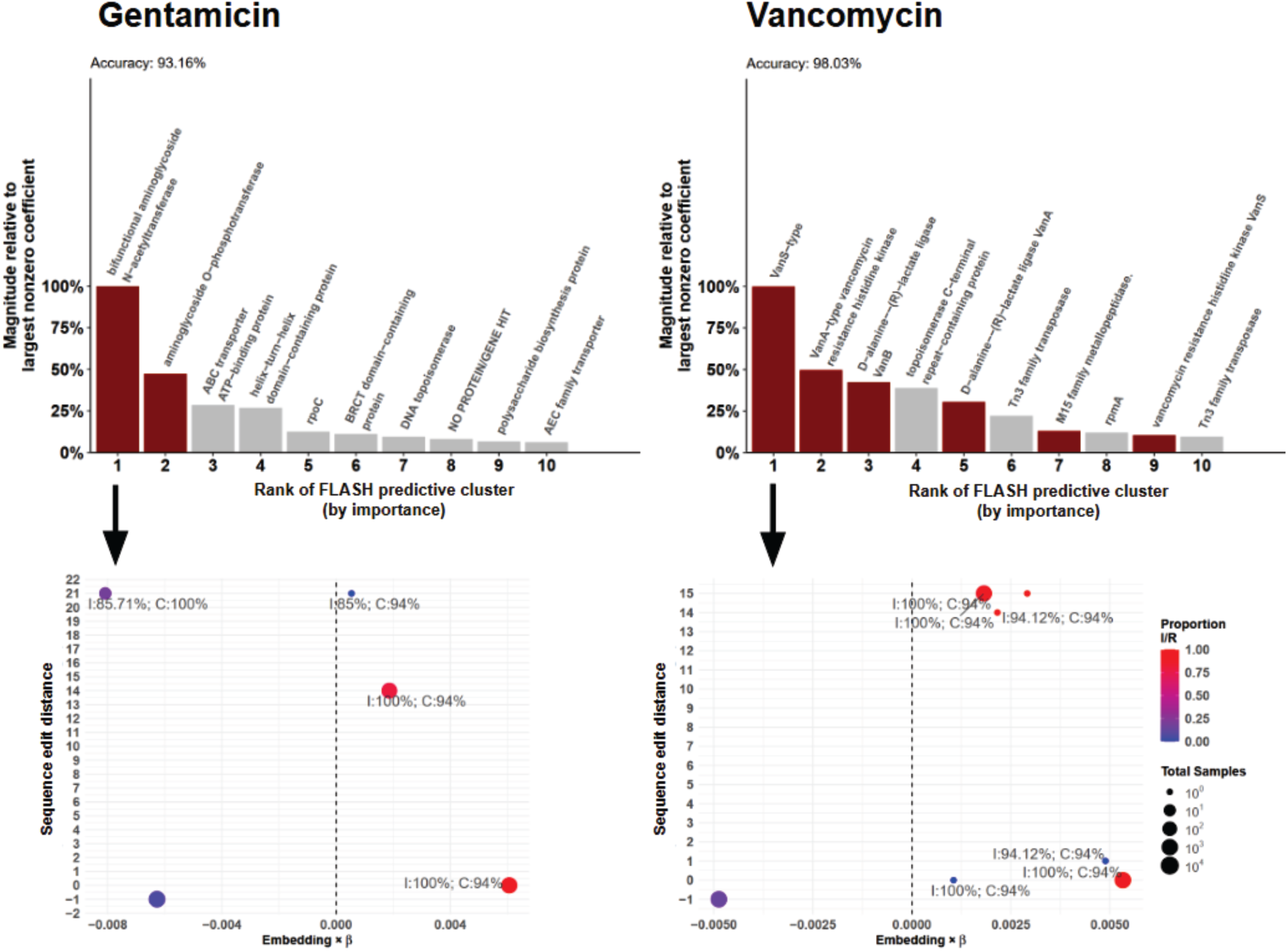
The top hits for gentamicin resistance and vancomycin resistance in *E. faecium* are driven by detecting absence of canonical resistance genes in susceptible isolates. In *E faecium*, the top 2 predictors for gentamicin resistance are hits to aminoglycosides, the main target of drugs of its class (left, dark red bars). The prediction is predominantly driven by a missing feature in susceptible samples indicated by the pure blue (all susceptible) point at the “-1” position in the scatter plot. For vancomycin (right), the prediction is driven by presence/absence of vancomycin resistant genes VanS, VanA, VanB (colored dark red). As with gentamicin, this prediction is driven by missing features in the susceptible samples (scatter plot). (I) refers to the percent identity of the closest blast hit. (C) refers to the percent coverage.

### FLASH predicts new pan-species predictors of resistance through protein homology

We analyzed how amino acid substitutions segregated with resistance in the strongest statistical predictors of resistance across 6 species of bacteria, and performed statistical tests of whether identified hits were drivers. We annotated predictive nucleotide features using protein homology, independent of genome alignment. First, the number of protein families hit in more than one species was highly significant, p<.005 (Methods). Next, we examined partitioning of resistance phenotype by amino acid change, as measured by normalized mutual information (NMI). The strongest value was perfect, in the 30S ribosomal protein S6, in prediction of resistance to gentamicin in *E. coli*, the known target of the drug.

The protein family with highest representations by number of species and the highest NMI was a family including GyrA, a topoisomerase and known target of quinolone antibiotics. It was found in *E. faecium, E. coli, M. tuberculosis, and S. pneumoniae* (Fig. 3D; Supp. File S4). The list also included DNA helicase PcrA – a protein known to regulate mobile genetic elements and several ABC transporters, previously implicated in resistance (Petit and Ehrlich 2002). Acetolactate synthase large subunits were found as predictors in 3 or more species (p< 0.005; Methods). This suggests a functional role for this protein or direct transfer by a mobile genetic element. Many other families were enriched and their multiple species hits are not explainable by chance including porins, penicillin binding proteins, and multidrug efflux pumps (Fig. 3D; Supp. File S4). Features related to mobile genetic elements were also top predictors of resistance such as mobilization proteins and the ISL3 family of transposases (Razavi et al. 2020), not previously reported to impact resistance in multiple species.

**Fig 3D.**
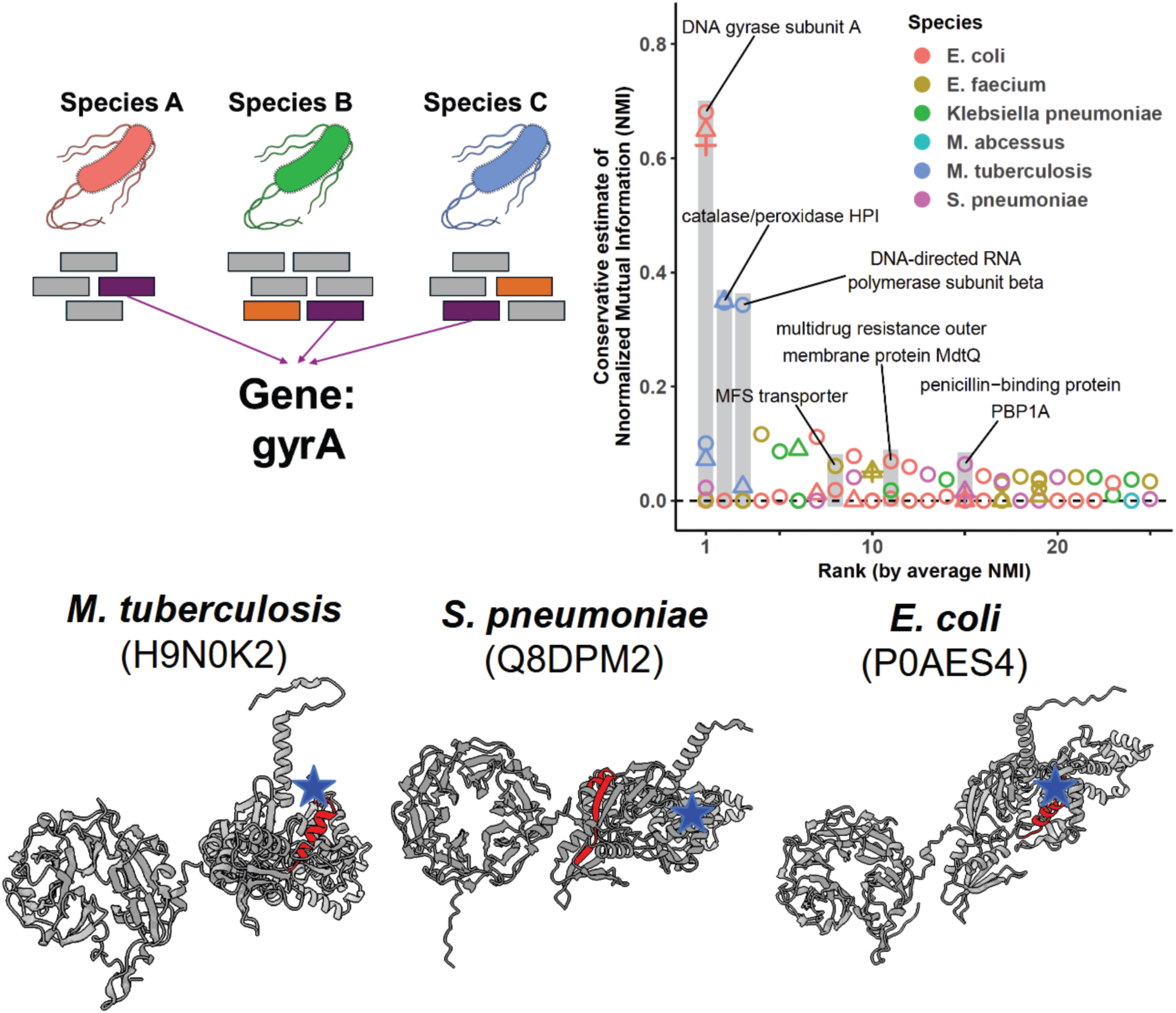
FLASH discovers shared drivers of antibiotic resistance pathways across six different bacterial species. Sets of proteins are arranged by their average normalized mutual information (NMI), that is, the measure of how well the metadata explain the amino acid translations in each FLASH predictive cluster (i.e. if a species can be partitioned into resistant and susceptible based on nonsynonymous amino acid mutations within a cluster it will have a high NMI). Each point corresponds to a FLASH predictive cluster for a unique species-metadata pair (different shapes in each column). Each point maps to a single protein which is grouped into an orthologous set across species. The set with the highest average NMI across predictive clusters occurs in four different species and corresponds to mutations in gyrA. The FLASH prediction is shown (bottom) for the top three species imposed on the respective protein structure (blue star indicates ciprofloxacin binding; red region is the location of the anchor identified by FLASH).

CD3337/EF1877 family mobilome membrane protein was among the top 5 predictors in different species for drugs sharing a mechanism of blocking protein synthesis (through targeting distinct rRNA subunits; *S. pneumoniae,* erythromycin and *E. faecium,* tetracycline). This approach is a framework to prioritize genes for further study and that the power of this approach will be increased with even more massive data mining.

### FLASH predicts drivers of clinical phenotypes including disease severity in Group B Streptococcosis

To better understand more complex phenotypes, we investigated clinical phenotypes without a clear monogenic cause, specifically asymptomatic carriage versus invasive disease (pyomyositis) in *S. aureus* and virulence phenotypes in *S. pneumoniae* and Group B streptococcus (GBS). In GBS, disease onset and isolation tissue are both difficult to predict clinically (Nusman et al. 2023; Chaguza et al. 2022). For the joint prediction of disease onset and isolation tissue (null accuracy: 25%, FLASH 4-class prediction accuracy of 49%), the top predictors included serC, an ABC transporter, nadR, which has been previously associated with virulence in GBS (Chen et al. 2024), and ardA also associated with multi drug resistance and the regulation of horizontal gene transfer (Supp. File S2). FLASH also predicted asymptomatic carriage versus symptomatic pyomyositis in *S. aureus* (accuracy 74%); the top driver of this prediction was the gene BrxA which has been shown to relate to the infectivity of *S. aureus* in vivo and contributes to the recovery of oxidative stress under infectious conditions (Linzner et al. 2019) (when excluding clusters with high missingness, Supp. File S5).

### FLASH predicts known and novel resistance mechanisms in human pathogenic fungi

We tested the generality of FLASH by applying it to pathogenic fungi. Azoles are first-line and commonly used antifungals (including fluconazole, itraconazole, voriconazole, posaconazole, and isavuconazole). They inhibit lanosterol 14ɑ-demethylase, the gene product of ERG11*/CYP51*, involved in the synthesis of ergosterol, required for fungal cell membrane integrity (Shafiei et al. 2020). However, mutations in ERG11 are not sufficient to predict resistance, and markers beyond mutations in ERG11*/CYP51* do not exist, to our knowledge, that have any individual azole class member-specific differentiation (Bader et al. 2013; Yu et al. 2020; Nishimoto et al. 2020; Whaley et al. 2016). As an initial check of its applicability to fungi, FLASH predicted mating type in *Aspergillus fumigatus* with 100% accuracy, identifying the variants in the mating type locus, MAT1-2-1, as the only predictor (Fig. 4A).

**Fig 4A:**
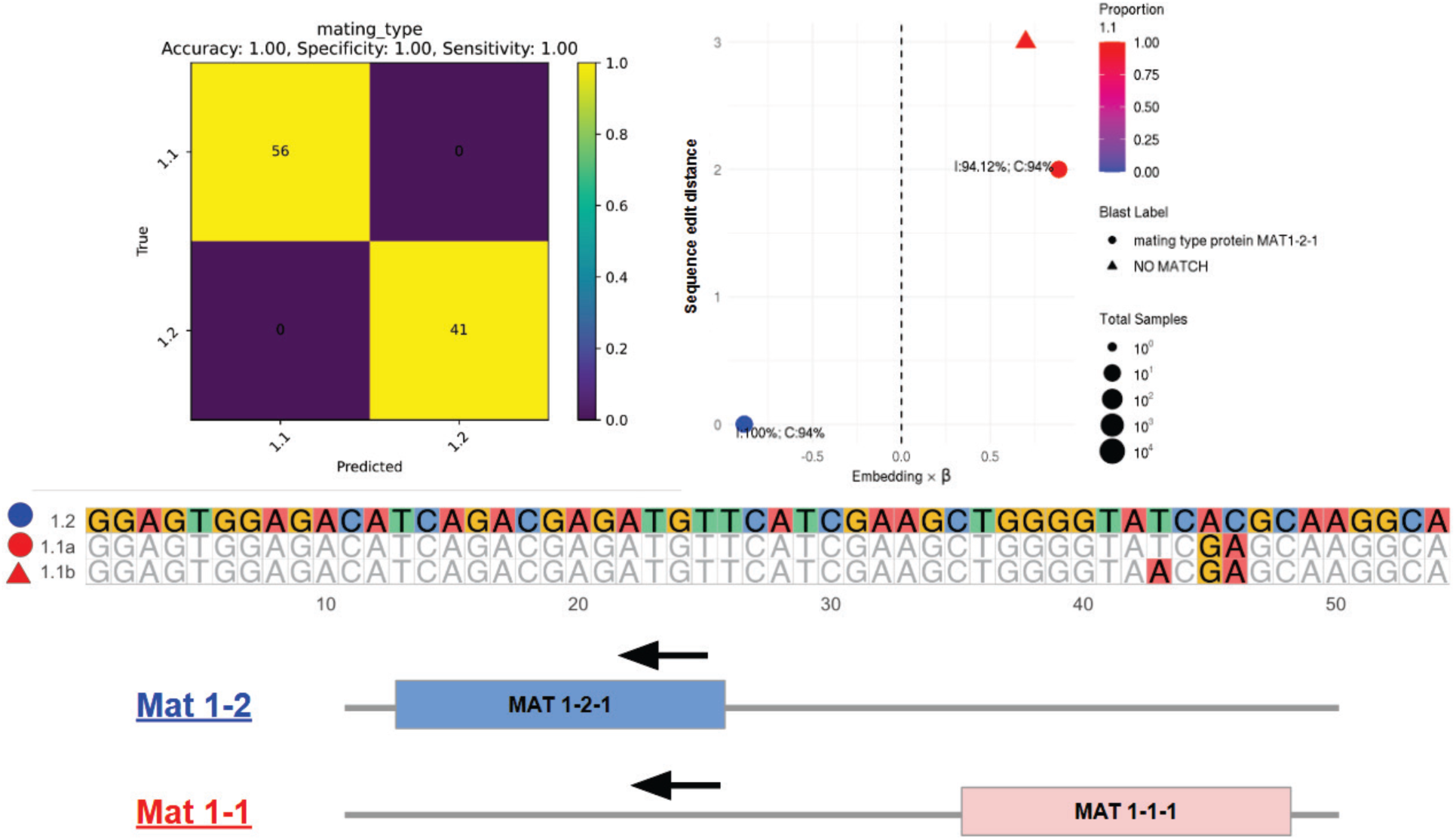
FLASH predicts mating type in *Aspergillus fumigatus* with 100% accuracy with ab initio rediscovery of the mating type locus. FLASH predicts mating type in a held out test set with 100% accuracy and perfectly recovers MAT1-2 as the only predictor consistent with known biology (scatter plot and confusion matrix). Variation in three positions distinguishes types 1.1a and 1.1b. FLASH predicts with a target mapping to variation in the C-terminal side of the highly-conserved HMG box in the Mat1-2 samples, a highly disordered region of the protein. In the Mat 1-1 samples the variation is in a homologous region to MAT1-2-1 upstream of MAT1-1-1.

FLASH recovered the canonical molecular mechanism of azole resistance in two disparate studies: one in *Candida tropicalis*, where it predicts ERG11 as the most predictive feature of fluconazole resistance with 94% accuracy (Methods, Fig. 4B), the other in *Candidozyma auris*: fluconazole resistance is predicted with 70% accuracy; the third most significant predictor is also *ERG11* (Supp. File S6). The top predictor did not map directly to any known genes via BLASTN or BLASTX; however it BLASTs to an intergenic region between CJI96_0003678, a SCP2 sterol-binding domain-containing protein and CJI96_0003679, a homology-predicted cytochrome P450 enzyme (the same family of protein as ERG11). The second most significant BLASTs to TAC1b, the transcriptional activator for CDR (Candida Drug Resistance) and MDR (MultiDrug Resistance) genes in *C. auris.* In *Candida tropicalis,* the variation in ERG11 includes 3 non-wild type sequences (Fig. 4B): one is the S154F missense mutation previously described (Siqueira et al. 2025); two others are nearby mutations T152A and T157I, present in an azole resistance “hot spot” in *ERG11* but to our knowledge not previously described (Jiang et al. 2013; Paul et al. 2022). In *C. auris*, the single Y132F variant drives the prediction of resistance. This mutation has been a major driver of high-level resistance in *C. auris* (Xu et al. 2023) and another species *Candida parapsilosis* (Choi et al. 2018).

**4B:**
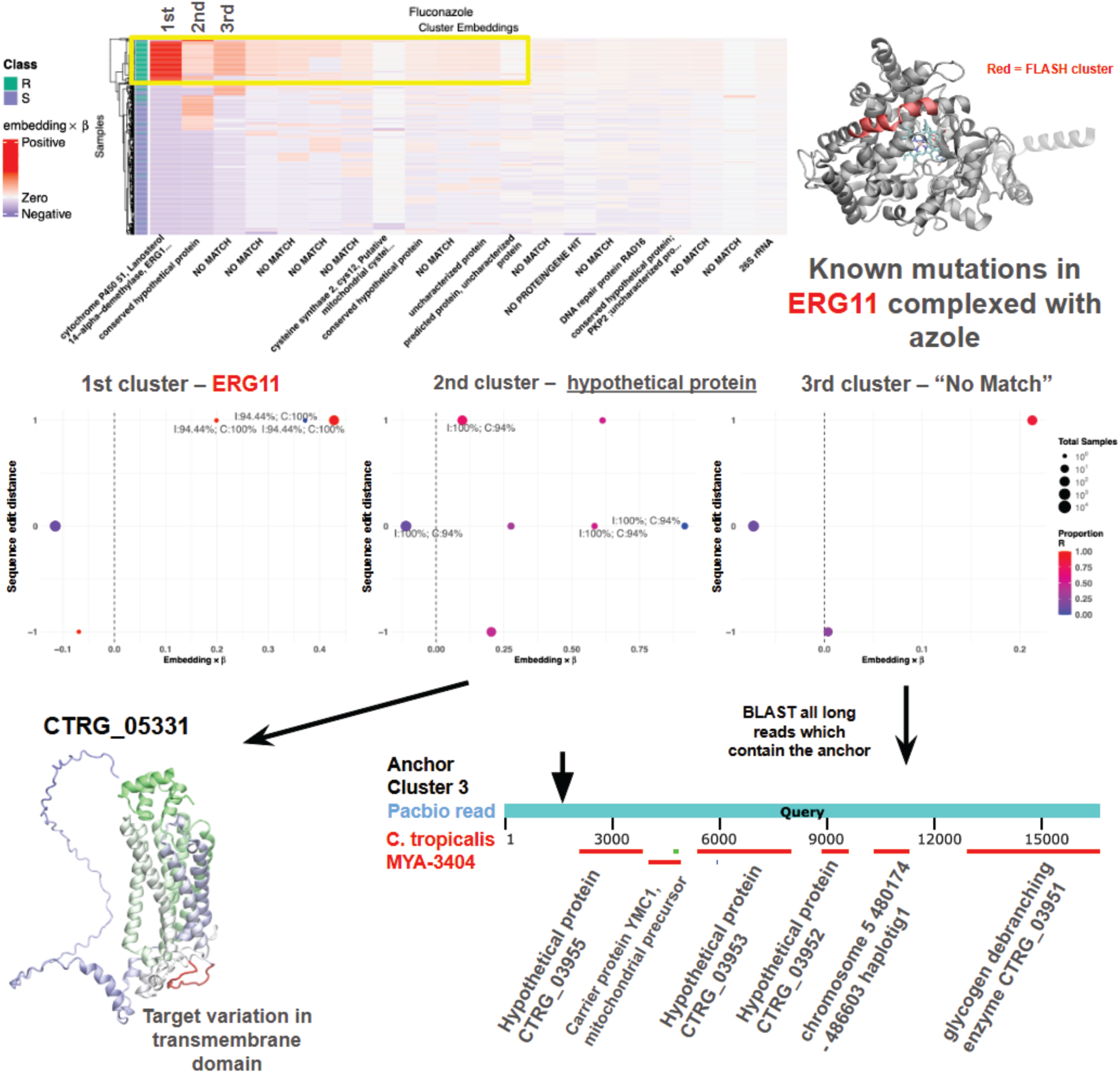
Fluconazole resistance is predicted with 94% accuracy, assigning highest importance to the canonical resistance gene and discovering novel predictors in *Candida tropicalis*. Predictors of azole resistance in clinical isolates of *C. tropicalis* are varied and many have no known blast hits to either protein RefSeq or the core nucleotide databases. The top predictor of fluconazole resistance is ERG11, the known target of fluconazole. The scatter plot shows the edit distance of anchor-targets that are involved in the prediction; variants highlighted in the sequence alignment and illustrated to be contained within a sequence fragment in a domain contacting the ligand fluconazole. Points at “-1” indicate a sample that is “missing” the cluster (i.e. it does not have any of the anchors that were used to define the cluster). Cluster 2 blasts to a hypothetical protein, CTRG_05331, whose structure matches a 7-transmembrane domain containing protein (Uniprot: C5MGX7). Cartoon representation in a color scale transiting gradually from green in the N-term to blue in the C-term. The fragment used for FLASH clustering is shown in red; the flexible low confidence portion is in the transmembrane domain; scatter plots show 7 variants detected, including absence of the anchor cluster that distinguish resistant and susceptible.

The second cluster in *C. tropicalis* mapped to a conserved hypothetical protein, CTRG_05331, whose structure (Uniprot: C5MGX7) matches a 7 transmembrane domain containing protein with structural similarity (via Foldseek) to a human GPCR chemokine receptor C-C type 9 (e-value: 6.7e-3), and to *S. cerevisiae* proteins RTA1 (e-value: 8.5e-5), RTM1 (e-value: 5.3e-3), and RSB1 (e-value: 1.7e-3) all of which have functions as transporters that export toxins from the cytoplasm. The mutations found by FLASH occur in the transmembrane domain (Fig. 4B). The third most significant cluster did not BLAST. It was, however, found in 82 PacBio long read sequences available from a subset of the isolates (Fan et al. 2023) as well as the *C. tropicalis* isolate from Y1000, and only that isolate. By BLAST, each of these long reads have segments aligning to the *Candida tropicalis* genome (GCF_000006335.3), with large deletions when mapped with BLAT (Kent 2002) (Supp. Methods). These reads have segments aligning to predicted genes identified in clinical isolates of pathogenic *Candida* (Butler et al. 2009) (Fig. 4B) and two genes containing a functional annotation: a glycogen debranching enzyme (CTRG_03951) and YMC1, involved in mitochondrial L-ornithine transmembrane transport (CTRG_03954; GO:1990575) (Fig. 4B).

The first cluster predicting posaconazole resistance BLASTed to a protein annotated tRNA methyltransferase; clusters ranked 2, 3, 5 and 6 did not BLAST (Methods). (Fig. 4C). For the 2nd predictive cluster, we analyzed 88 PacBio reads containing the anchors in this cluster; these reads contained sequences with high homology to a set of 6 known and hypothetical genes (at least 10/88 of the matching long reads; BLAST e value ∼0; Fig. 4C; Supp. Methods). The gene contained in the largest number of reads (n = 65) is a conserved hypothetical protein (CTRG_04444), predicted to be an integral membrane-bound transporter, with low complexity domains and DUF2421, a domain involved in Brefeldin A sensitivity, a fungal-produced antibiotic which has been shown to confer resistance to azoles via inducing aneuploidies (Zhang et al. 2024). Lower ranked clusters of posaconazole resistance had BLAST hits with similarity to Candida Drug Resistance (CDR) ABC transporters (Prasad et al. 2019) and to an exocyst complex component which has a previous, though indirect implication in resistance to azoles (Albehaijani et al. 2022) (Fig. 4C).

**4C:**
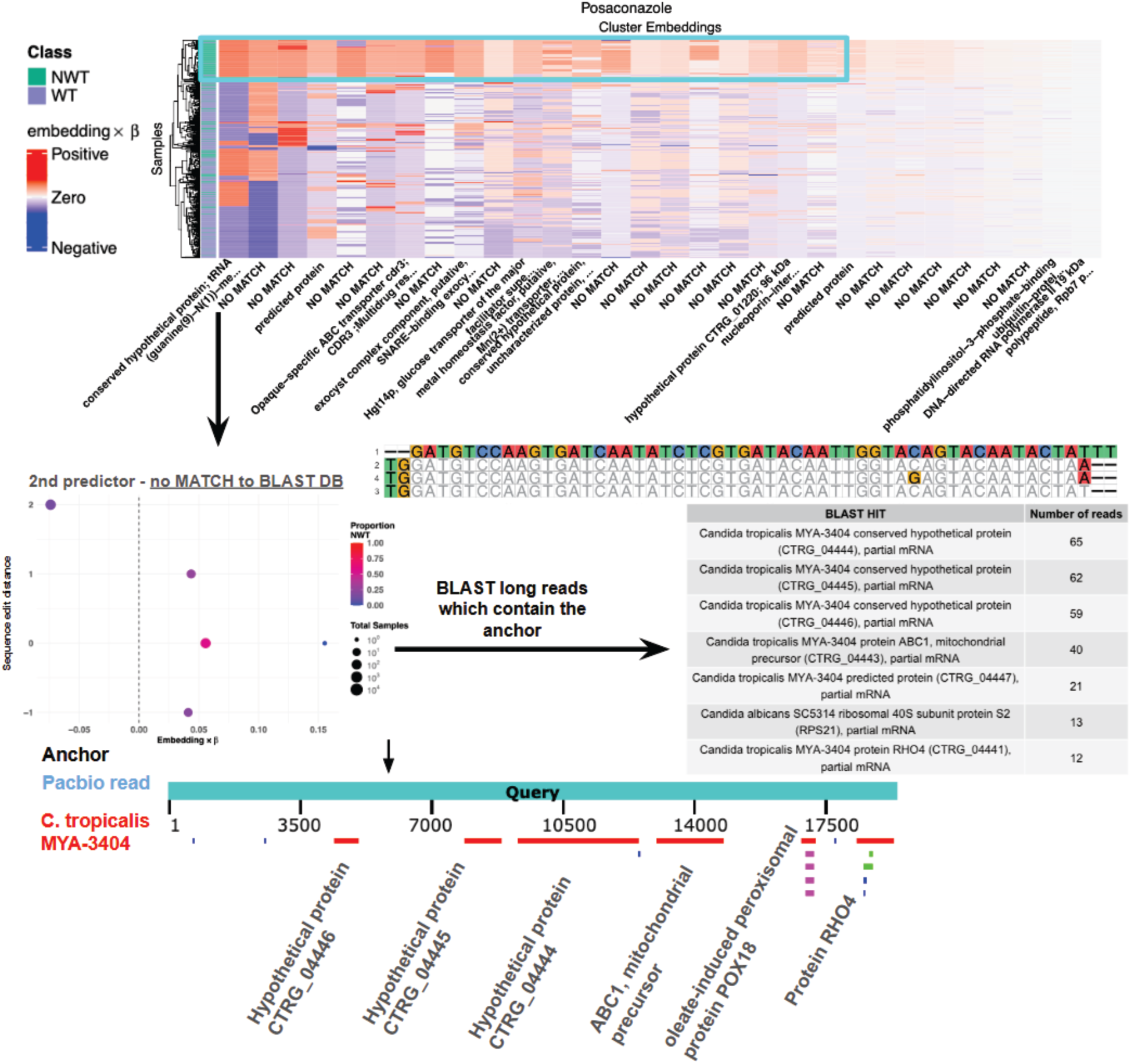
Posaconazole resistance is predicted with 81% accuracy. Resistant (non wild-type NWT) isolates in candida have a strong correlation across many of their top predictors. The top predictor blasts to a tRNA methyltransferase. BLAST of long reads containing the anchor from cluster 2 summaries: commonly hit genes are represented by counts of the number of reads supporting them, most being hypothetical proteins. An example read and the location of the anchor and other genes with high homology in it are illustrated at the bottom of the panel.

Itraconazole’s second highest predictor had homology to a hypothetical protein, MSN4, phosphatidylinositol-binding protein, a protein with plausible involvement in maintaining cell wall integrity (Schiavone et al. 2016). The most significant cluster did not have any BLAST hits, but analysis of long reads containing the clusters (Supp. Fig. S1, Supp. Methods), identified that the cluster is proximal to Candida Drug Resistance Gene CDR3 suggesting FLASH reports on a locus proximal to genes involved in resistance (Supp. Fig. S1).

### FLASH identifies a hypervariable gene cluster in pathogenic fungi associated with virulence

ERG11, the canonical resistance marker, was the fourth most predictive feature for voriconazole resistance. The most predictive feature had only partial BLAST hits, with the best match to a predicted protein, CTRG_04842, and a CFEM-domain containing protein, CTRG_04948 (in *C. albicans,* the ortholog is named Surface antigen protein 2; Uniprot: Q5A0X8). In other fungi, CFEM proteins have been shown to contribute to virulence and inhibit plant immunity (Huang et al. 2023). CTRG_04948 is annotated containing NADH-quinone oxidoreductase subunit E. CTRG_04948 is homologous to CTRG_04939, annotated as including a ribonuclease E domain and a CFEM domain.

Further analysis of these genes using SPLASH Sample Anchor Target Count (SATC) files (Methods) showed that strains have high levels of sequence variation in CTRG_04948 that segregate with resistance. One anchor in this gene had 17 variant sequences with high count (>5) with significant deviation from the reference, which contains only two copies of the anchor (Fig. 4D). Further, the diversity of the target sequences and number of variants is predicted by resistance phenotype. The variants are not clonal, ruling out the possibility of ‘passenger’ variation. Together, this suggests that the gene is part of a locus undergoing either rapid evolution and likely expansion and contraction of the repeats or copy number in resistant isolates and missed by assembly (see Supplement for further discussion).

**4D:**
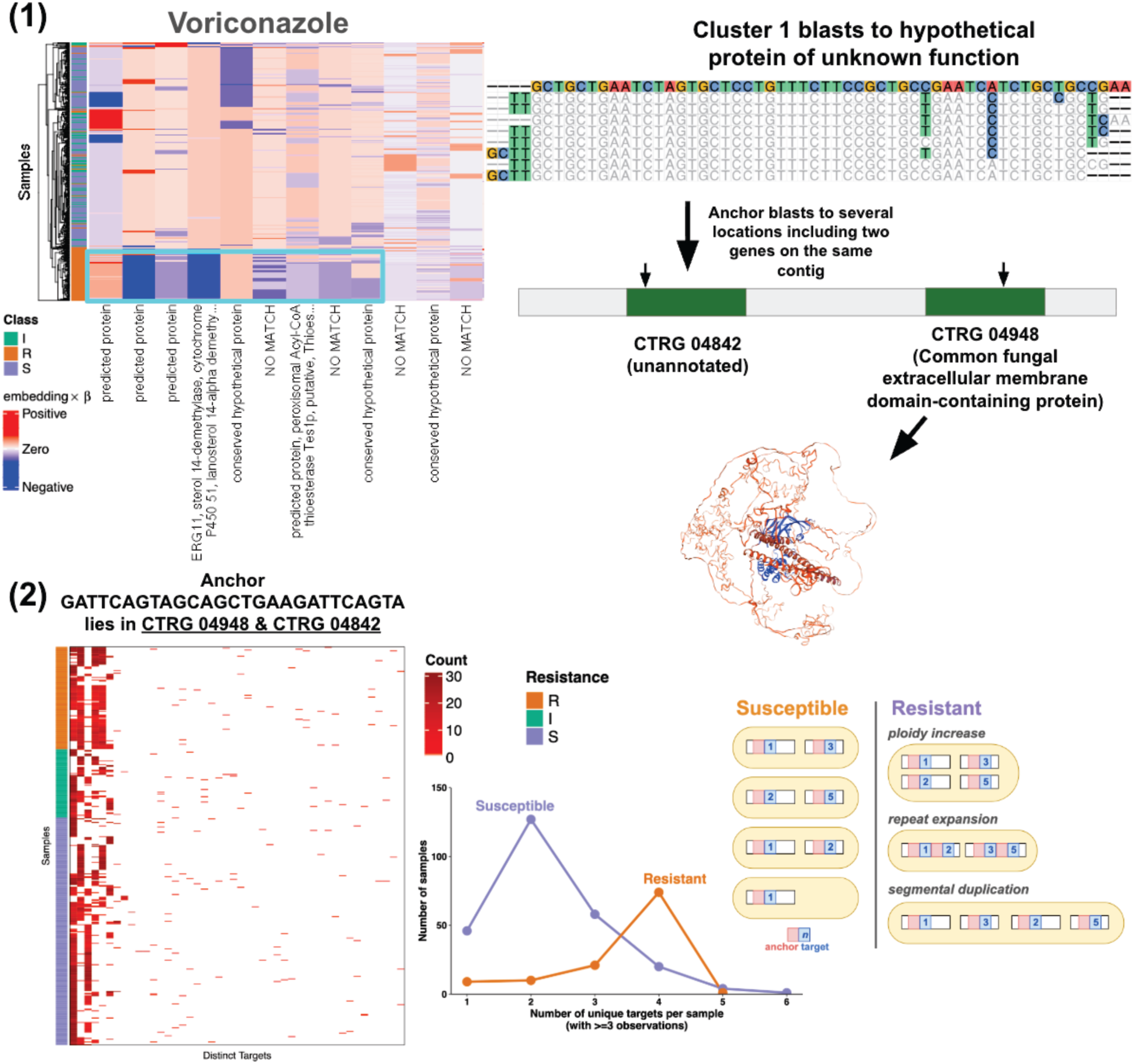
The top predictor of voriconazole resistance is in an extracellular membrane protein with the number of unique variants associated with the resistance phenotype. Heatmap of cluster contributions to the prediction of voriconazole resistance (top left) in *C. tropicalis*. The first three clusters driving the prediction are predicted proteins with poor BLAST and BLAT hits; the first hit is to hypothetical protein CTRG_04842. The fourth cluster is ERG11. Selecting all long reads containing the anchor from CTRG_04842 in the first cluster and blasting the long reads, reveals many hits to other hypothetical proteins. Proteins CTRG_06242 and CTRG_06243 appear in most reads, and within a read, have many blast hits. Multiple reads match CTRG_04948 (C5MFV5, structure from SWISS-MODEL) which has been annotated as similar to potential cell-surface antigens and a CFEM (Common fungal extracellular membrane) domain contain protein; blast hits show evidence of rearrangement or structural variation (see supplemental methods A). One anchor GATTCAGTAGCAGCTGAAGATTCAGTA lying in CTRG_04948 has many different configurations of targets and significantly higher levels of diversity. Visualizing SPLASH anchor variation in anchors with >=5 counts illustrates more unique targets occurring in resistant isolates than in susceptible or intermediate isolates (bottom). Mechanisms that explain this observation include polyploidy, gene duplication, or repeat expansion (schematic).

### FLASH, predicts resistance phenotype in yeast isolates separated by >400 million years of evolutionary divergence

We also tested FLASH in the Y1000 *Saccharomycotina* samples, species sharing a common ancestor over 400 million years ago, FLASH prediction accuracy for Flucinazole subset to only Piachales was 81.4% (Methods; Supp. File S6). FLASH identified a cluster mapping to β-1,3-glucan synthase, a key cell wall structural component in several fungi and an important clinical biomarker for invasive *Candida* infections. In severe infections, echinocandins (e.g. caspofungin), which target β-1,3-glucan (Wagner et al. 2023) are used in combination with azole therapy.

Some sequences within the FLASH-identified cluster BLASTed to predicted or uncharacterized genes annotated as β-1,3-glucan synthases and which had homology to GSC2 and FKS1, paralogs in *S. cerevisiae* which are part of the β-1,3-glucan synthase complex. This predicts a functional role for these uncharacterized sequences and shows that FLASH could aid in annotation of such uncharacterized proteins.

Further, FLASH-detected variation was not explained by phylogeny (Fig. 4E). Phylogenetic analysis of *Saccharomycetales* (*Saccharomycetes*), *Serinales*, *Pichiales*, *Alaninales* (all *Pichiomycetes*), and *Dipodascales*, shows amino acid mutations at positions 1228-1232 (Uniprot: P38631) repeatedly gained or lost in orders separated by ∼350 million years. These variants correlate significantly with resistant vs susceptible isolates (p=0.0028, Fisher’s exact test) and less significantly when comparing by order (*Pichiomycetes* vs *Saccharomycetes*; p=0.09, Fisher’s exact test). The mutations are in the glycosyl-transferase domain, which catalyze the addition of glucose to glucan chains. While β-1,3-glucan is not a direct known target of fluconazole or voriconazole, variations in the enzymes producing it may represent a genetic background that is synergistic to their targets (Fig. 4E). Clusters predicting fluconazole resistance did not include *ERG11,* possibly because FLASH did not aggregate highly evolutionarily diverged sequences into the same predictive clusters.

**Fig 4E.**
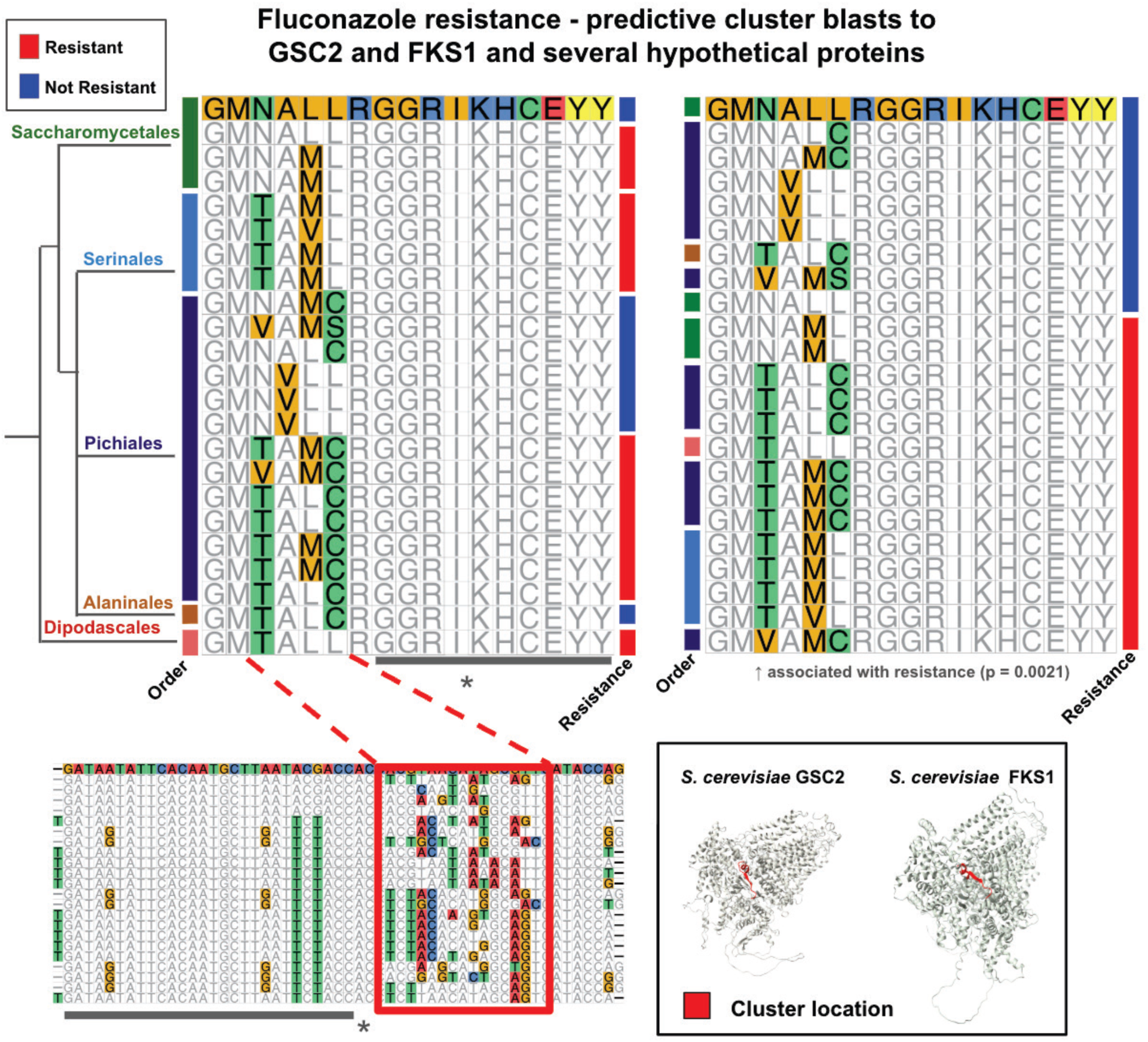
Variation in β-1,3-glucan synthases highlights high levels of resistance-associated sequence diversity across Saccharomycotina evolution. Fluconazole resistance in Saccharomycotina is predicted by sequence variation with blast hits to β-1,3-glucan synthases (top). These hits are across many different anchor-target combinations across a diverse set of Orders. At the protein level, these nucleotide changes occur in only four consecutive amino acids with certain changes enriched in resistant relative to susceptible isolates and seemingly unrelated to phylogenetic relationships (red box). The rest of the sequence has high conservation (the region indicated by the asterisk is the anchor region used for clustering and will bias the sequences here to be more conserved). The domain is observed in both FKS1 (Uniprot: P38631) and GSC2 (Uniprot: P40989) (alpha fold structure predictions in S. cerevisiae with anchor-target region highlighted in red).

### FLASH extends in generality to viral RNA and discovers interactions between bacterial hosts and infective phages

We analyzed viral RNA isolates collected during the recent to identify variation that could predict host range (Nguyen et al. 2025). All sequences mapping to a eukaryotic host were excluded. FLASH had 95% accuracy distinguishing between the viral host (chicken, turkey or cattle; Fig. 5A; Supp. File S7). The most significant clusters used for the prediction are several hits to polymerase PB2 and hemagglutinins HA1 & HA2. Anchor absence in exactly one species occurred in most predicted clusters, showing an important and common driver of FLASH prediction is presence/absence of a gene or anchor sequence (eg. Fig. 5A); that is, FLASH is sensitive to variation in the target, but variation in the constant anchor region is another important signal (see Methods).

**Fig 5A.**
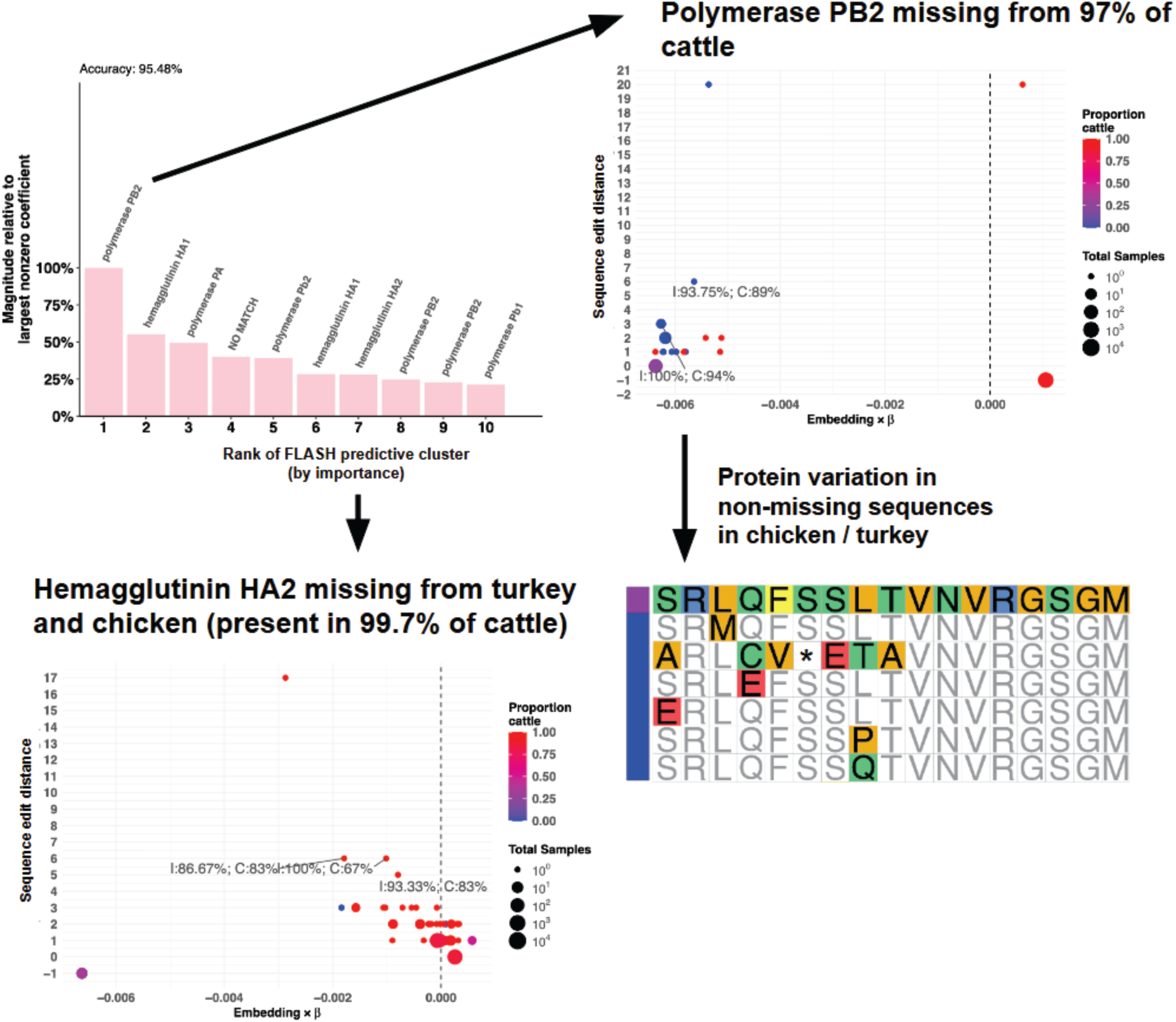
FLASH models predict host range with 96%% accuracy on RNA viruses, like H5N1, with high-mutation rates. Host (chicken, turkey, cattle) is predicted in H5N1 with 96% accuracy. The 1st top predictor indicates that in cattle, for PB2, the samples are entirely missing any anchors from this cluster. On x axis: magnitude of the embedding prediction; y axis represents sequence edit distance between each embedded sequence; color represents fraction of metadata class with that label; blast Coverage (C) and identity (I) are indicated. The value at -1 indicates the samples that are missing any target variation. The MSA represents the turkey/chicken variation (blue) and one sequence present in cattle and chicken (purple). * indicates a stop codon. Within-cattle variation that is not missing is only present in ∼30 of 950 cattle samples. In the 7th cluster, annotated hemagglutinin HA2, the prediction is also driven by an anchor which is not detected, but in this case the anchor is not detected in birds.

To further explore the capability of FLASH to detect viral host range and variation, we analyzed a study of 148 clinical isolates of *Vibrio cholerae* circulating in Bangladesh during the years 2016-2019 (LeGault et al. 2021). 44 ICP1 phage genomes isolated from water sources were also assembled and provided as orthogonal to sequenced isolates. Resistance to each phage was experimentally found to fluctuate by year (LeGault et al. 2021). We tested if FLASH could identify which phages were year-matched to circulating strains by taking all pairs of strains and phages and predicting if the two sampled in the same year, a proxy for the phage being able to successfully infect and replicate (LeGault et al. 2021). Marginal predictors using phage or bacterial features alone had accuracy of 74%. Models with features including bacterial sequences, phage sequences and their interactions selected only features that were interactions between phage and bacterial host, and accuracy improved by 12% to 86% (Fig. 5B, Supp. File S8). The most predictive features included transporters, porins and type IV toxin-antitoxin system AbiEi family antitoxin, a bacterial abortive infection system recently shown to be a widespread regulator of phage infection (Dy et al. 2014). This is evidence that FLASH independently identifies temporal genetic variation in phage defense systems (LeGault et al. 2021). In addition, genes of unknown functions including those that did not BLAST and a protein with an annotated COG3014 domain are significant predictors, suggesting they may be involved in phage defense.

**Fig 5B:**
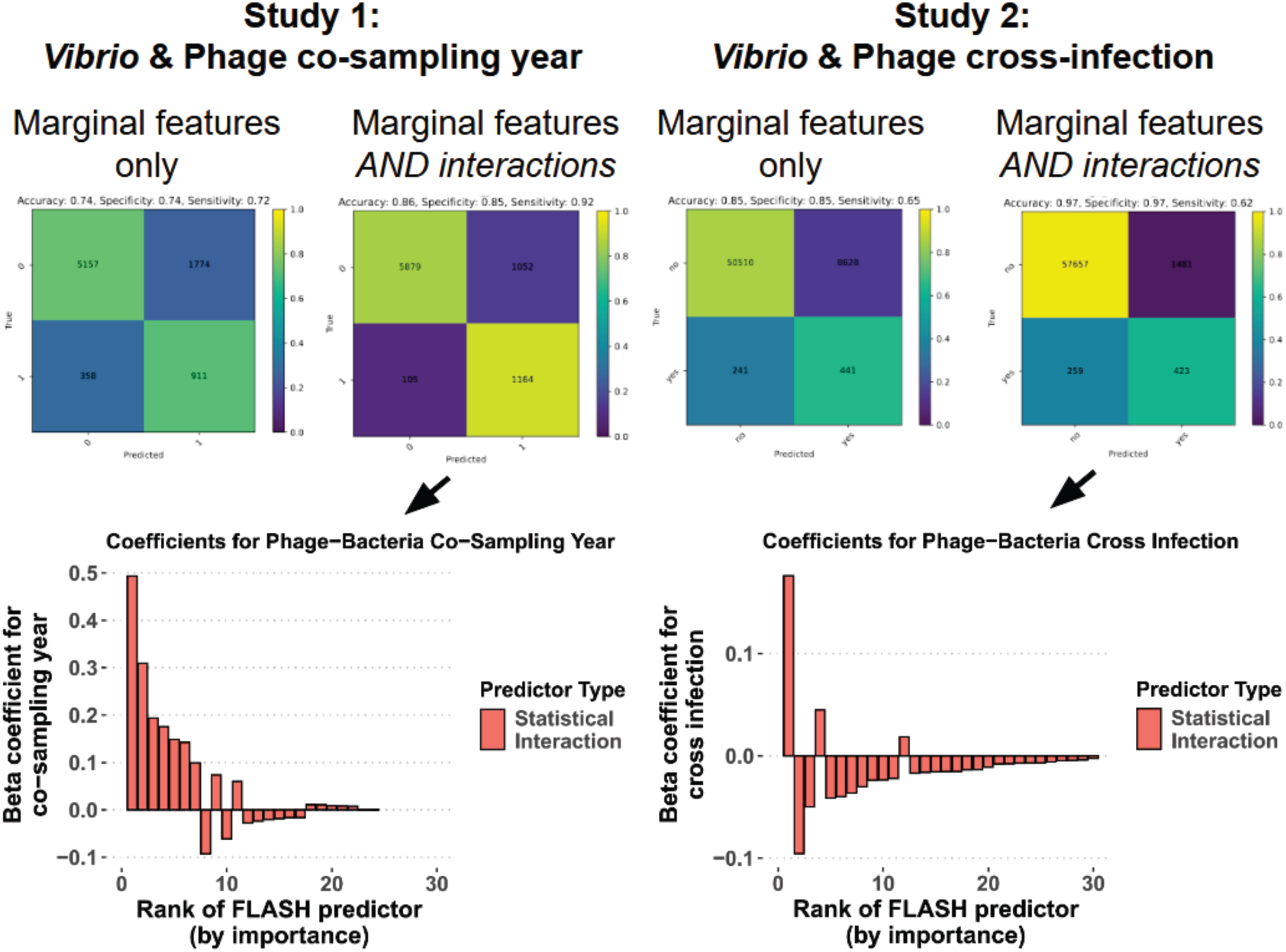
FLASH models discover biological interactions and predict phage-host infections with 86% and 97% accuracy in two distinct studies of phage host (*Vibrio) range*. *Vibrio* and phage pairs of samples were generated and used to predict co-sampling in the same year (LeGault et al. 2021)(left) and cross-infection studied experimentally (Kauffman et al. 2022)(right). In both cases, the inclusion of genetic bacteria-phage interactions (in addition to the marginal features) increased the prediction accuracy by >10%. Statistical interactions dominated the predictions of co-samping year and for cross-infection (barplots).

To test the generality of this approach, we used FLASH to predict phage-bacterial interactions by analyzing experimentally measured host ranges of 239 phages and 256 *Vibrio* species (Kauffman et al. 2022). The steps for the prediction followed from above (see methods). Marginal prediction accuracy, using only the initially significant clusters found independently in bacterial and phage sequences, was 85%. Adding the interaction terms between all bacteria and phage clusters to the prediction increased the accuracy to 97%, with the majority of the selected features as interactions (Fig. 5B, Supp. File S8). One of the top interactions was between a cell surface ABC transporter and a phage hypothetical protein (KMD65_gp02) (Kauffman et al. 2018).

## Discussion

Historically, assembled genomes, genes within them and their assigned functions have been a prerequisite to predicting phenotype. FLASH bypasses these steps while identifying canonical and novel determinants of phenotype (such as in antibiotic and antifungal resistance) by efficient statistical algorithms that run on raw sequencing data, and indeed reverses the order: predicting genotype from phenotype. FLASH is very simple to run, modular, and highly efficient and achieves blind ‘universal’ phenotypic prediction accuracy that matches or exceeds custom bespoke studies. Further, it begins to open analysis that could not be performed with our previous work, SPLASH, as it provides phenotype prediction from single-samples, bypassing the need for comparative analysis, a previous requirement.

To our knowledge, FLASH recovery of known resistance is more consistently powerful—across datasets and the tree of life—and more parsimonious than any other available methods. In addition to identifying known and plausibly novel genes in the antibiotic and antifungal applications, FLASH can pinpoint known mechanistically important domains, such as binding and catalytic sites. We hypothesize that this performance will significantly improve when its sequences are integrated into large language models, and extendable to human drug targets, both a subject of current work. The parsimonious, interpretable attributions provided will likely enable analysis covariation, that is conditional epistatic interactions, in the future.

FLASH may be especially impactful for discovery in microbes, tumors and non-model organisms where reference genomes are poor approximations or unavailable; or, in genetic loci with high variability that cannot be captured by a reference genome (such as our findings in *Candida*). Due to space, we have not presented an analysis of other applications (such as single cell RNA-seq data), but we believe FLASH models will provide significant insight in this realm as well: it remains difficult to find SNPs, splicing or other sequence variants correlated with cell type or state with current methods. In the clinical realm, we have shown a proof of principle that FLASH could be usable with point of care diagnostics by directly parsing raw reads, including out-of-species or error-prone long-reads. Moreover, we believe that optimizing feature clustering, especially amino-acid-based clustering, will enable more robust predictions across more divergent organisms, a proposition to be tested in future work.

FLASH also is able to nominate genes with new potential biological roles in phenotypes, (e.g. the virulence phenotypes we explored here). We speculate that our findings of a potentially new diversifying locus in *C. tropicalis*, through prediction of voriconazole resistance, are indirect with a large portion of the effect due to a virulence phenotype versus a direct interaction with the drug. Further work on FLASH, including its integration with multimodal deep learning models will refine predictions aiming to distinguish features predicted to be direct and mechanistic versus confounded with other phenotypes. As this is a fundamental and longstanding goal of trait and phenotype prediction, it may not be possible. Finally, FLASH can predict phenotypes not possible to predict with GWAS, such as phage host range.

In summary, FLASH provides a new approach to linking phenotype and functional prediction of genome-feature type (such as genotype) by generating a simple alternative approach that is based on direct statistical analysis of raw sequencing data. A critical step in FLASH missing from SPLASH is similar in spirit, but an alternative to multiple sequence alignment: to group similar sequences to make them comparable so that they can be used as a single statistical predictor. To our knowledge, that has not been previously suggested. Many parameters, such as how clusters are defined, and the length and or character (nucleotide or amino acid) can be explored to improve performance. In doing so, it provides a “lower bound” for prediction accuracy without using higher dimensional multivariate modeling. We anticipate that with larger training data, use of deep learning will enhance the ability of FLASH to predict phenotypes and assign functions to DNA, RNA and protein sequences.

### Limitations

There is no explicit causality in the prediction between clusters that FLASH produces and biological causality. Similarly, FLASH, by design, could and does aggregate different genes into the same cluster. This could be considered a limitation or an advantage. A current limitation is that to show a proof of principle, and save computational costs, FLASH currently limits itself to 50,000 feature clusters as it prepares data to be embedded. It also does not incorporate any information about the reverse complement of the DNA or paired-end information. Increasing efficiency and adding more contextual information are both extensions that will increase the power of this method. Similarly, the parameters have not been optimized to pick a best value, especially the kmer length. Our choice of 54 (27 anchor and 27 target) is based on previous work, but it is possible that a different kmer length would increase accuracy of the predictions.

In our analysis, we have also not generally discriminated between amino acid changes and or feature presence/absence. As in any computational prediction, we also cannot exclude that the difference in some predictions, such as those made for H5N1, could be influenced by inconsistencies in data collection across species and tissue types (Good et al. 2024). Finally, a subject of future work is to discriminate between active mechanistic interactions with drugs and genetic events that may modify these interactions.

## Supporting information

Combined Supplemental Info

## Acknowledgments

We thank members of the Salzman lab for useful discussions, and Gavin Sherlock, Chester Yu, Zhiyong Zhang, Elysse Grossi-Soyster, Aaron Behr, Jeremy Rock, James Hickling and Leonor Garcia-Bayona for comments on the manuscript. We thank Mu Gao for providing some AlphaFold alignments and mapping FLASH sequences to structures.

J.S. is supported by the National Institute of General Medical Sciences grant R35 GM139517. M.K. and S.D. are supported by the National Science Center, Poland (project no. DEC-2022/45/B/ST6/03032). A.F.C. is supported by the National Institute of Allergy and Infectious Diseases grant DP2 AI171122.

## Author Contributions

Conceptualization: JS; Data curation: DJC, MCH; Formal Analysis: DJC, MCH, PLW, JS; Funding acquisition: JS; Investigation: DJC, MCH, PLW, AR, JS; Methodology: DJC, MK, SD, JS; Project administration: JS; Software: DJC, MK, SD; Supervision: SD, JS; Validation: DJC, MCH, PLW; Visualization: DJC, PLW; Writing and editing manuscript: DJC, MCH, AR, PLW, MK, AFC, SD, JS

## Methods

FLASH is an interpretable, scalable machine-learning method that operates on k-mer (nucleotide sequences of length k) representations of each sample to jointly analyze sample phenotypes and genetic features (Fig. 1). The k-mer inputs to FLASH can be directly extracted from raw sequencing data: here, we use anchors found by SPLASH (Chaung et al. 2023). Anchors are k-mers in sequence that evidence sample-dependent variation in downstream sequence, which we have shown contain rich regulatory information called targets (Chaung et al. 2023; Kokot et al. 2024). In principle, the method has no reliance on SPLASH, and any set of kmers could be used as input. FLASH has three key steps.

In step 1, FLASH performs “anchor clustering”, an efficient, parameter-free and scalable step that can be thought of as analogous to multiple sequence alignment. In this step, FLASH groups anchors that are similar and within a defined edit distance.

This step guarantees that anchors in distinct clusters will be at least a fixed distance apart—here 5 (user defined parameters). It also allows for minor variation (eg. SNPs) between strains or species to be grouped together without pairwise or multiple sequence alignment, allowing anchors to be ‘aggregated’ when they differ with a small number of variants. We implemented and explored clustering in both edit distance in nucleotide space and translated nucleotides that converted anchors into amino acid sequences in silico. In general, both methods produced similar results (Supp. Fig. S2).

In step 2, FLASH creates a per-sample representation of sequencing reads into clusters: when using SPLASH input, it ranks each anchor in each cluster based on the effect size (Chaung et al. 2023; Kokot et al. 2024). FLASH processes samples independently: for a) each sample and b) each cluster, it chooses the anchor with the highest effect size and then records the most abundant target for that anchor as the representative for the anchor cluster. If no such anchor is found in the sample, the sequence is defined as the highest effect size anchor in the cluster followed by Ns to represent the absence of a target. Step 2 can be used to produce a map from a different sample or an assembled genome using a new functionality called FAFQ filter (Methods) that allows phenotype prediction without running SPLASH.

In step 3, for each sample, FLASH maps sequences of identified clusters to numbers. FLASH either maps sequences to an indicator variable—“one hot encoding” or OHE—by encoding per-sample presence or absence of a specific anchor-target pair or it embeds data in a nucleotide language model, HyenaDNA, trained on a large set of prokaryotic sequencing data (Nguyen et al. 2023) tokenized by splash anchor-targets. FLASH summarizes embeddings by averaging across nucleotides, further reducing dimensionality by choosing the embedding dimensions with the highest variability (Methods). In zero shot prediction, FLASH uses principal component analysis of the embedding representation of sequence clusters (see Fig. 3A). This clustering is done without prior knowledge of the samples’ identities.

In the FLASH analog of fine tuning, when phenotypic metadata is available, FLASH can run in supervised mode, dually predicting phenotypes and the features responsible for prediction via a cross validated multinomial generalized linear model with elastic net (Yang and Hastie 2024). Throughout, prediction accuracies are reported on test data completely held out of training. The model assigns variable importance to clusters; the larger the magnitude of the coefficient beta the more important the sequence cluster is in the prediction. The cluster which has the highest magnitude contribution to the prediction is called the first cluster throughout the manuscript, the second cluster has the second highest, and so on.

### SPLASH

FLASH is built on top of SPLASH, an unsupervised, reference-free, algorithm that discovers regulated sequence variation through statistical analysis of *k*-mer composition (Chaung et al. 2023; Kokot et al. 2024; Baharav et al. 2024). We use SPLASH as the starting point to identify statistically significant kmers that are variable across a given set of samples. Previous work (Chaung et al. 2023) showed that how well samples and targets can be partitioned into disjoint classes is an effective metric (the effect size) to prioritize functionally important anchors. We used anchors with high effect size and with length 27, again a convention based on previous work (Chaung et al. 2023), as features input to the first step in FLASH. In FLASH, anchor absence is encoded (as Ns following the anchor, see below), and is also used as a predictor. We note that shorter anchor lengths can capture important genes and predictors missed by longer anchors (See discussion below and Supp. Fig. S3). SPLASH was run with an anchor length of 27, a target length of 27, a poly ACGT length of 8 and a gap length of 0 for all data (except in some cases discussed below when testing anchor length).

### Select anchors

FLASH uses SPLASH anchors to predict patterns either unsupervised or supervised using metadata such as antibiotic resistance. It does this both to predict these metadata categories in a held-out test set and to assess which kmers have the highest effect in contributing to a given metadata category.

First, SPLASH results are filtered to only include anchors from the set of samples which have an effect size above 0.6. Anchors in the bottom 10th percentile by their total number of samples are also removed to eliminate sparsity from the dataset.

### Step 1. Cluster anchors to align features across samples

Anchors are clustered using one of two methods outlined below. They are then ordered within each cluster in decreasing order arranged by effect size. Clustering eliminates anchors that come from similar parts of the genome and would be too correlated, thus, each cluster represents a distinct genetic feature that will be used for the prediction of metadata downstream. For downstream analyses, we use the top N=50,000 of these clusters.

There are two different clustering approaches used for aligning anchors into feature clusters: ***1) Shift Distance:*** Pairwise distances are calculated between all anchors and edges are drawn between anchors that are connected with a shift distance of five or less. After drawing an adjacency network connecting all anchors that fit this criteria, connected components are labeled as clusters. To ensure that the anchors within each cluster are not too distant from one another, they are further filtered by calculating the Levenshtein (edit) distances to a single anchor representative (chosen as the highest SPLASH effect size anchor) and removing anchors greater than 5 edit distance away. *This results in some anchors and anchor types (such as highly polymorphic ones) not being included as inputs to the model. **2) Amino-acid-based approach***: For the amino-acid based approach, each anchor sequence is translated into six reading frames (three forward and three reverse) using the specified translation table. Translations shorter than seven characters (using anchors of length 27) are discarded. Anchors are grouped by masking m=3 random characters from the translated sequence, N=300 times and checking for matches in a lookup table. Additional trimming at the beginning and end of sequences (up to j=2 characters) accounts for shifts. If a match is found, the anchor is assigned to the matched cluster. If no match is found, a new cluster is created, and all masked or trimmed variants are added to the lookup dictionary for future comparisons. These clusters (as their nucleotide sequences) are also filtered to ensure they are within 5 edit distance away from the highest effect size anchor in the cluster.

### Step 2. Post-process anchor clusters

To prepare sequences for downstream processing, one anchor+target is selected from each feature cluster for each sample so that each sample is represented only once in the feature cluster. To do this, FLASH works down the list of anchors (ordered by effect size continuous, a SPLASH statistic) and identifies, for each sample, the highest abundance *target* for that anchor. If the first anchor is not present in a sample, it attempts to find targets for the second anchor in the cluster and so on. If, for a given sample, there are no targets for any of the anchors, the first anchor (highest effect size) is selected and a target composed of all Ns is added (indicating missingness). The chosen anchor+target is the “cluster representative” for that sample.

Note, that the metric for ordering clusters, effect size continuous, used throughout, is especially useful when using large datasets with many isolates. For smaller datasets, an alternative metric, effect size binary can be used to better identify anchors which are significant in the raw data. We use effect size binary, primarily for the processing of the *Candida tropicalis* data and other fungal data after comparing it and the effect size continuous metrics (Baharav et al. 2024).

For each feature there is a set of sequences consisting of one anchor followed by its single target that had the highest abundance in each sample. Here, the choice to use just a single target was made as it simplifies the analyses as they were conducted across mostly haploid bacterial species. For high effect size SPLASH anchors, the highest abundance target usually makes up ∼95% of all targets observed for that anchor (Supp. Fig. 4A&B). Even for diploid organisms, such as *Candida tropicalis*, the distribution of target abundances skews highly in favor of the most abundant target representing most of the variation (Supp. Fig. S4C). This is likely driven by the way SPLASH prefers partitions of samples that are distinct between two organisms, driving down the frequency of heterozygosity in high effect size anchors.

These anchor-target pairs are arranged as a matrix where the first column in the matrix is composed of the anchor target pairs for the first cluster across all samples, the second column is the second cluster (anchor + target) and so on for the remaining feature clusters. These anchor-targets are then either one hot encoded or transformed into embeddings using HyenaDNA trained on bacterial DNA.

### Filtering initial cluster features

In ∼64% of clusters in E. faecium, 20% of samples are encoded with NNNNs as missing targets. We provide a method to post process clusters as they are prepared to discard any that are missing greater than N% of samples. We ran this regime by filtering for clusters missing >5% of samples and included the results in Supp. Table S5.

### Step 3. Embed or One Hot Encode Anchor clusters

Next, the sequences from each feature cluster are prepared for downstream prediction in one of two ways: 1) Using embeddings from a DNA-language model to represent each sequence numerically or 2) by one-hot encoding the data or representing the presence or absence of each unique sequence in a cluster as an indicator variable.

### Embeddings

For language model embeddings, each unique sequence (anchor-target pair) in the matrix is embedded in a pre-trained DNA language model, HyenaDNA (Nguyen et al. 2023) in order to have a numerical representation of the sequences. HyenaDNA uses a nucleotide vocabulary A, C, T, G, and N as well as several special tokens. A HyenaDNA model with 8 layers and 128 dimensions was pre-trained on ∼50,000 bacterial isolates (Supp. Table S4) formatted by concatenating SPLASH anchors with their most abundant targets similar to the way the sequences are encoded in the pipeline. Our model performed comparably to a model already pre-trained on humans (<u>huggingface.co/LongSafari/hyenadna-medium-160k-seqlen</u>; Supp. Fig. S5).

The sequences were embedded in this model to produce a 256-long vector per nucleotide. By averaging over each nucleotide in the sequence, this step produces one vector of length 256 that represents the entire anchor target. Using these embedding representations, the feature matrix was then created for downstream prediction of metadata. The feature matrix has rows equal to the number of samples and columns equal to the number of features multiplied by the number of dimensions in the model (256 dimensions).

### One-hot encoding

One-hot encoding (OHE) can also be used to transform the data for prediction by representing the presence or absence of each possible cluster representative as an indicator variable. The cluster representatives for all samples were determined in Step 2. For each feature cluster, there will be a set of columns in the vector representing each of the cluster representatives seen in the data. For each cluster, every sample will have a 1 in the column corresponding to its cluster representative and 0s in the other columns. The resulting feature matrix has rows equal to the number of samples and variable columns depending on the number of unique cluster representatives. If a cluster representative is detected in the training data, but there are no samples with that representative in the test data, the column is still there, just encoded entirely with 0s.

### Predict phenotypes using a generalized linear model

To reduce the feature space that is used in the prediction step, the embeddings for each feature cluster are filtered, selecting only the top 100 highest variance columns. These embeddings can be either normalized (centered and scaled) or left unnormalized prior to prediction. For OHE, the feature matrix is not further filtered. Then a model is fit one by one for each metadata category on this feature matrix X using a grouped multinomial elastic net implemented in adelie (Yang and Hastie 2024) using cv_grpnet and the parameters *min_ratio = 0.1*, *lmda_path_size = 100*, *n_folds = 5*. The elastic net groups correspond to each of the clusters (such that all embedding dimensions for a cluster share a grouped penalization). Training data uses 80% of the samples from the least represented class and balances samples for each remaining class; the remaining samples are the test set. This approach does not suffer from the biased estimates of test error introduced by approaches that use cross-validation (Bates et al. 2024).

Throughout the paper, unless otherwise stated, we fit a model and calculate accuracies using the steps described above and 1) clustering using the nucleotide-based approach, 2) using the embeddings from the HyenaDNA model for prediction, and 3) normalizing the embeddings prior to prediction. Train and test set sizes are as specified in the prior paragraph.

### Accuracy calculations

In all cases, accuracy is calculated on all available test data after subsetting out the balanced training classes. Accuracy is calculated on the testing set as the number of true positives plus the number of true negatives all divided by the total size of the testing set.

In figures reporting performance of accuracy across models, in order to facilitate comparison between datasets, we drop any data points whose prediction is not on binary phenotypes. Throughout the main text, accuracy is calculated on all classes who have more than 15 observations in the data set.

In comparing accuracies to previously-published work, for *E. faecium (Coll et al. 2024)* and for Y1000 *Saccharomycotina* (Opulente et al. 2024; Harrison et al. 2025), we follow slightly different procedures depending on each study. For *E. faecium*, the study calculates performance measures on the whole dataset since it uses a genotype-database to predict resistance. We compare their numbers (tabulated in Table 3 and Appendix 2) by calculating their accuracies and comparing to our test accuracies calculated in the held out test set. To compare our accuracies to those calculated for Y1000, we compare against the antibiotic resistance accuracies using the same approach. Accuracies for Y1000 (resistance) are calculated by disincluding clusters with > 5% missing data across all samples. Accuracy for Fluconazole is restricted and calculated exclusively in Pichiales before comparing it to the original study. See Fig. 2A and Supp. Table S2 for the accuracy measurements reported in these papers.

### Anchor attribution

Each multinomial elastic net that is trained using adelie identifies several anchor clusters (as described above) that are nonzero in their contributions to the model prediction. Because the feature matrix that is the input to adelie contains multiple embedding vectors for each cluster of sequences, the output is filtered to identify which clusters had were attributed the highest nonzero beta coefficients by the elastic net (max across the absolute value of all multinomial coefficients per cluster).

Sequences in each of the anchor clusters are Each of these sequences is then subject to a homology search either using BLAST or using a lookup table from SPLASH (https://github.com/refresh-bio/SPLASH) to identify any antimicrobial features from the megaRes database v3 (Bonin et al. 2023). Because a cluster can have multiple sequences, we report all annotations for each sequence in a cluster that has a nonzero coefficient. For example, if a cluster contained sequences that each BLASTed to two separate genes, each gene is reported for the cluster. When plotting these results, the information is collapsed down to generate one label per cluster containing information about all unique homology hits.

For nucleotide homology, BLASTN (with parameters -db core_nt -evalue 0.1 -task blastn -dust no -word_size 24 -reward 1 -penalty -3 -max_target_seqs 5 and filtering for taxids given in Table S1) is run against the core_nt database before using efetch to grab all overlapping gene or feature names of the blast hit from its NCBI record.

For protein results, annotations are generated using BLASTX (with parameters -evalue 0.1 -max_target_seqs 10 and specifying -query_gencode with the correct translation table of the species or set of species and filtering for taxids given in Table S1). This search is performed for all 54mers from nonzero coefficient clusters against the refseq_protein database reporting the title of the protein hit as the annotation. Each of these analyses are restricted to a taxon filter in order to improve performance. For individual species, the taxon is the genus and for larger subsets the taxon is one that is representative of all species.

### Datasets

Details on 16 datasets analyzed using FLASH can be found in Supp. Table S1 (Mets and Morin 2024; Coll et al. 2024; Heinz et al. 2024; Young et al. 2019; Lo et al. 2019; Southon et al. 2020; Chaguza et al. 2022; Kim et al. 2023; Etienne et al. 2021; Fan et al. 2023; Misas et al. 2024; Harrison et al. 2025; Shen et al. 2018; Nguyen et al. 2025; LeGault et al. 2021; Kauffman et al. 2022). To train the HyenaDNA nucleotide embedding model, data in Supp. Table S4 were run through SPLASH and compiled into sequences of anchors and targets. We report the size of the data and the time it took to analyze these samples through FLASH. For all paired-end data, we used R1 for each sample when running the initial SPLASH computation. All accessions used for FLASH can be found in Supp. File S9.

### Model and cluster comparisons

To compare the difference in performance between different model types (bacterial hyena model with nucleotide embeddings normalized versus other model choices and OHE) and different clustering approaches (nucleotide shift distance or amino acid based), all pairwise comparisons between each of these different choices are aggregated across all metadata categories, filtering for only metadata categories that are binary.

For each dataset and for all data aggregated together, a two-sided Wilcoxon signed rank test is used to see if the difference in accuracies across all metadata is statistically significant. p-values are corrected using a Benjamini–Hochberg correction.

This process was used to compare model choice. In addition to hyena and OHE, an amino-acid based model is also compared in addition to choices about whether to normalize the embeddings prior to prediction. Across all datasets, hyena embeddings (normalized) and one-hot encoding performed similarly but better than the other methods on average (Supp. Fig. S1). For this reason, the analyses were focused on hyena nucleotide embeddings (normalized) and one-hot encoding-based models.

We also compared between our pre-trained model and a model already trained on the human genome by HyenaDNA (Supp. Fig. S5). We see that our models perform comparably to other pre-trained models and have slightly higher performance when restricting to predictions with higher accuracy. We use the HyenaDNA model we pre-trained for all embeddings of data unless otherwise stated.

This process was also used to compare choices for clustering. As mentioned above, there are two main choices made for clustering the data: one nucleotide-based approach and one amino-acid based approach. These choices were compared with an additional nucleotide-based approach (masked nucleotide clustering) which uses the same procedure described for the amino acids but on untranslated nucleotide anchors and setting m=5, j=5, and N=1000. The amino acid approach and the shift distance nucleotide approach performed similarly (Supp. Fig. S1).

### Number of model features

For each dataset, the number of input features and the total number of features selected by the model were determined by grabbing the number of columns from the input feature matrices used for the prediction and by counting the number of nonzero features in the final adelie predictions. The accuracy of each metadata prediction (subsetting for metadata with binary categories only) versus the number of input features to the model were compared (Fig. 2B). Similarly, accuracies and the number of model features *selected* by the elastic net were compared (Fig. 2B)

### Analyzing anchors within a dataset that have not been seen by FLASH

We subset the *E. faecium* dataset by specifically selecting a single study from a compilation of data from 21 different studies (Coll et al. 2024). We ran SPLASH and FLASH on exclusively the data isolates from this study to artificially limit the number of features in the training set. The rest of the isolates were then post-processed to extract overlapping clusters and format them for prediction using the FAFQ filter package (see below). In this way, we ensure that the test data FLASH is seeing contain sequences in their clusters that were never seen when fitting the FLASH model with the embeddings and elastic net. The accuracy of this prediction and the number of clusters never seen before testing are reported in Fig. 2D.

### FAFQ Filter package

The SATC format is a binary file format used in SPLASH. It encodes, for each <u>S</u>ample, <u>A</u>nchor, and <u>T</u>arget, the related occurrence <u>C</u>ount. Originally designed as a SPLASH internal format, SATC can also be used as one of SPLASH’s outputs. For such a use case we designed a set of helper utilities. An overview of their high-level functionality is presented in Supp. Fig. S6. Internally FAFQ filter utilizes a custom hash table combined with Bloom filter to represent a set of anchors for which the most frequent targets are to be identified. Each input sample is processed independently (possibly by different threads). Processing a single sample works as follows. For each read its anchors are extracted, and only those that are part of a predefined set are stored in a standard C++ hash table. For each such anchor a separate hash table is maintained to count targets. Once all reads have been processed, for each anchor (iterating in lexicographical order) targets are sorted in a decreasing order of frequency. The most abundant target(s) per anchor are then written to the output SATC file.

### FLASH unsupervised clustering

Using FLASH in an unsupervised manner, we can identify existing clusters of samples (such as known sequence clusters or serotypes) across different bacterial species without the need for assembly or alignment. To do this we run FLASH through Step 3, embedding feature clusters and formatting the data for supervised prediction. Using normalized embeddings, we grab the embedding with the highest variance per cluster to format a matrix that is *n* samples x *p* clusters (only keeping the highest variance column per cluster). We filter out samples that belong to a cluster with <30 samples (<75 for GBS). We perform scaled and centered principal component analysis (PCA) on this filtered matrix and then perform t-SNE on the top 10 PCs (theta =0.2, max_iter=2000). For Fig. 3A, we color the metadata classes which have the highest number of observations in each dataset. We also illustrate *S. pneumoniae* and *K. pneumoniae* in Supp. Fig. S7.

To verify the quality of this clustering, and to make predictions of clusters for which we have no metadata, we used Hierarchical DBSCAN (Hahsler et al. 2019) to generate artificial cluster labels. We then calculated the adjusted rand index (Rand 1971) between the assigned labels and the known labels as a measure of similarity between the two clusterings (between 0 and 1 where 0 is no similarity and 1 is exact similarity, accounting for stochasticity). Adjusted rand indices (0.954 for group A strep, 0.686 for group B strep, 0.827 for S. pneumoniae, and 0.887 for K. pneumoniae), where 0 is poor clustering and 1 is a perfect match to known clusters, performed comparably or markedly better in some cases than existing k-mer based approaches (Lees et al. 2019).

### Resistance annotations in bacterial species

We analyze the performance of FLASH on antimicrobial phenotypes in six species: *E. faecium, E. coli, S. pneumoniae, M.tuberculosis, M. abscessus* and *K. pneumoniae*. This includes both exploring the accuracy metrics as well as the investigation of the clusters with the highest predictive value for a phenotype. Each feature cluster is assigned gene identity via protein and nucleotide homology searches with BLAST (See above).

Each bacterial resistance dataset was also annotated with the MegaRes database using SPLASH lookup table (Bonin et al. 2023). Any hit to a sequence in this database was recorded for a given antibiotic and all predictive clusters across all antibiotics were aggregated (Supp. File S10).

Additionally, InterProScan was run on all hits that successfully had a blastx hit to the RefSeq protein database (Jones et al. 2014). Superfamily annotations (Wilson et al. 2009) were collated into the heatmap shown in Supp. Fig. S8.

For each dataset, we use the HyenaDNA nucleotide language model (with normalized embeddings) that was trained on bacterial sequences. We use nucleotide clustering for all species. All results reported are for these choices of model and clustering. To better compare accuracies, antibiotic sensitivity phenotypes were binarized by lumping Intermediate and Resistant labels into a single category.

### Identifying genes that predict resistance in multiple antibiotic-resistant species

Protein blast annotations are aggregated for any cluster that is non-zero for an antibiotic resistance phenotype across the datasets (*E. coli, E. faecium, S. pneumoniae, K. pneumonia, M. tuberculosis, M. abscessus*) for which we have experimental antibiotic resistance phenotypes. We query all RefSeq protein sequences for the anchor targets that were predictive for each antibiotic. To identify proteins that are similar across species, we tabulate a pairwise similarity matrix by conducting a pairwise alignment (using BLOSUM62; gap opening = -10; gap extension = -0.5), between every nucleotide sequence with a BLASTX hit across these six bacterial datasets. Before performing pairwise alignments, we substitute the nonstandard amino acid U for C in places that it appears.

After pairwise alignment, we identify sets of features that have alignment scores greater than 500 and then assemble a graph, and choose connected components as the clusters for downstream analysis (the cutoff of 500 was chosen after testing the sensitivity of the procedure to produce non-singleton sets of proteins). For each feature in a set, we calculate the entropy of its amino acid sequences to score the number of nonsynonymous mutations that appear approximately for a given feature. Entropy is computed as the relative probabilities of every observed amino acid and will be 0 if there is only one translated AA per cluster. We also calculate normalized mutual information (NMI) for each predictive cluster (per species-metadata category pair) using the metadata labels (R or S) for each amino acid sequence to construct a contingency table defining the probability of resistance or sensitivity given a specific translated sequence (If there is only one amino acid translation in the cluster, the value will be 0). The metric is calculated as the mutual information of the contingency table divided by the square root of the product of the marginal entropies (Supp. File S4). Using this clustering and these metrics, orthologous sequences are identified across species and aggregated into sets. A set can include proteins from ≥ 1 species but here we are especially interested in sets of proteins that have representation from multiple species. This clustering and annotation is conservative and only includes a sequence or an amino acid translation if it had a blast hit. Sequences in the cluster are absent if they did not have a blast match under our parameter choices. This represents a set of sequences that have successfully matched a protein in protein_refseq via blast and as such missing information for a given species or cluster does not mean the sequences are non-existent. Full nucleotide sequences for the top 10 hits for each prediction are present in Supp. File S3.

To explore the probability of such sets being found, we consider the generalized birthday problem where k is the number of species-metadata pairs and N is the number of possible sets in the dataset. We want to consider the probability a species-metadata pair occurs with another in the same set using predictors of at most rank r. In Supp. File S4, we report the sets for all species-metadata pairs with any rank of predictor. There are k=41 different species-metadata pairs across 6 species and N=1438 unique sets in this table.

Across sets, there are 17 which contain at least 2 distinct species predicted with rank less than or equal to 20. For i species, this event is Poisson approximated with lambda = ((k choose 2) - sum_i (k_i choose 2))* r/N with k =41, i=6, r=20 and N = 1438. The values for k_i here are 26,13,6,1,4, and 1. The probability of seeing at least 8 sets with at least 2 distinct species in our dataset overall is statistically significant at p = 0.033.

In one of these sets, we have 3 different species mapping to an acetolate synthase gene with all predictors in the top 3. We ask what is the probability that there is a set with at least 3 species with rank 3 or less. Following from above, this event is Poisson approximated with lambda ((k choose 3) - sum_i(k_i choose 3)) * (r^2/N^2) with k = 41, r=3 and N=1438 and the same values for i and k_i as above. The probability of seeing at least one such event is statistically significant (p = 0.0037).

We note that this approximation is conservative as it specifically includes only proteins which were already found as hits across all species and metadata categories

### Using different length k-mers to define SPLASH anchors

SPLASH and FLASH both allow for the specification of different length kmers (although 27 is the default used here). As in previous work, we chose k=27; compared to k=8 in Supp. Fig. S3.To explore the effect of different-kmer lengths, FLASH was also run after setting the anchor length, k, to be 8 and the target length to be 31. There are a few benefits for using short 8mers to represent the data for downstream analysis. One motivation for doing this is that it will allow the discovery of highly abundant, replicative mobile elements that are under selection. These are known to confer antibiotic resistance in escape pathogens. If using splash with 8mer anchors, the abundance of the targets could be driven by higher copy number of mobile elements which are often associated with phenotypes like antibiotic resistance (Partridge et al. 2018).

To explore this, after running SPLASH on 8-mers, FLASH was run leaving everything the same but modifying the parameters for kmer size. Instead of attempting to cluster the 8mer anchors (as there are many fewer possible anchors when k=8) they were simply ordered by effect size, and similarly the top 50,000 clusters were selected for downstream analysis. Results are reported in Supp. File S11. The accuracy of the predictions using this approach was tested in *E. faecium* and *E. coli* against the accuracies from predictions based on the analysis with anchor and target length both set to 27 (Supp. Fig. S3).

### Generating scatter plots of edit distance

Scatter plots are generated for each of the clusters using a summary of the BLAST hits for each cluster as well as which samples each sequence in a cluster was present in. The color of the point indicates the proportion of the samples that contain a specific sequence in the plot. The shape indicates a shared blast hit. The x axis indicates the embedding for that sequence multiplied by the beta coefficient for the current metadata class (seen in legend). The y axis indicates the edit distance to the most abundant anchor-target sequence (after aligning the clusters with MSA and trimming the edges to avoid inflating edit distance due to shifted anchors). This visualizes the extent of the variation in each cluster, how the prediction was influenced by it, and how split different sequences are between samples.

### Generating heatmaps of FLASH nonzero coefficients

The heatmap representation aggregates this information onto one plot by plotting each sample per row and each cluster per column. The color of a cell indicates the embedding value that a given sample had in that cluster multiplied by the beta coefficient for that cluster from the elastic net. This is used to visualize correlation across different clusters and across groups of samples.

### Visualizing cluster location in proteins

We searched for AA locations in the BLASTX hits of several proteins and downloaded their structures from AlphaFold, or their structures with small-molecule binding via AlphaFill. We then used ChimeraX (Meng et al. 2023) to color the locations of these hits. We also had assistance in generating these figures using AF2Complex (Gao et al. 2022).

### Predicting phenotypes in fungal data

FLASH produced exact (100%) prediction accuracy in held-out test samples for *Aspergillus fumigatus* when predicting mating type (1-1 vs 1-2). We analyzed this prediction by exploring the single cluster that was deemed significant for the prediction. We compared this cluster the MAT 1-2 gene in the *A. fumigatus* reference genome (GCF_000002655.1) and observed that the sequence present in isolates with the 1-2 phenotype had 100% identity to this gene while the sequences in this cluster from the other mating types had slightly different accuracy.

Because many of the antifungal datasets are small or very sparse across phenotypes, throughout our analysis of fungi we chose to use a different method for reordering clusters that is better at identifying larger effects in the input anchors (see above section on cluster reordering (Supp. File S6).

FLASH had consistently high prediction accuracy for azole resistance in *Candida tropicalis*: fluconazole (94%), itraconazole (90%), posaconazole (81%) and voriconazole (81%). These accuracies were calculated using Resistant and Susceptible phenotypes in fluconazole, itraconazole, and posaconazole. In voriconazole, when calculating accuracies, intermediate phenotypes were lumped in with resistant ones to create a binary phenotype on which to train a model and evaluate performance. When discussing attribution, we used the 3 category model and did not lump intermediate and resistant phenotypes. Discussion of attribution followed the methods described for bacteria, using BLAST and BLASTX annotations to assign homology to different feature clusters. One key change is that many more of these clusters were labeled as hypothetical or did not have a BLAST or BLASTX hit to the databases we queried. In order to explore features of these hits we used long read data that was available from the same *Candida tropicalis* study where these isolates came from (Fan et al. 2023). We searched each long read for the presence of an anchor from the cluster of interest (Supp. Methods). We then BLASTed these long reads (which had the anchors) against the *C. tropicalis* reference and explored which genes had significant hits (e<0.001). We further explore genes that had BLAST hits in multiples of these reads. In one case, in voriconazole resistance, we also visualized the target distribution of an anchor present in one of these genes. We did this by subsetting the SPLASH SATC file for presence of the anchor of interest and then plotting the counts of each of its targets in each sample as a heatmap. We plotted the sum of targets that were present in >=3 samples across all samples as a distribution (Fig. 4D).

In the Y1000 dataset, we analyzed predicted features for the amino-acid-based clustering, hypothesizing that the results would be more interpretable by capturing increased rates of synonymous nucleotide mutations. We explored β-1,3-glucan (which was a top cluster in fluconazole) by translating the sequences for each sample and aligning them in an MSA. We ordered this MSA by phylogeny and by phenotype and compared how the mutation was associated with either resistance or with order by using Fisher’s exact test. For organisms, we grouped the N to T AA mutation (position 3 in Fig. 4e) into T and not T as well as resistant or not resistant. We analyzed the resulting 2x2 contingency table with Fisher’s exact test (p= 0.0021). Using the contingency table for T and not T versus order—collapsed into *Pichiomycetes* (*Serinales*, *Alaninales*, *Pichiales*) vs *Saccharomycetes* (*Saccharomycetales*)—we also tested for the relationship to the phylogeny (p=0.09, Fisher’s exact test).

### Predicting resistance in genomes never seen by SPLASH or FLASH

Step 2 of FLASH produces a map from raw sequence measurement such as a fastq file from a sample to a short representation which is one 54mer per anchor cluster. If SPLASH were not run, it is possible to hash all kmers from a sample, such as through KMC, and search for an anchor matching the anchor cluster, and define the most abundant target to plug into the above procedure. It is also possible to specifically search for all anchors in the cluster a priori through a tool that will be introduced here called FAFQ filter. FAFQ filter enables efficient retrieval of most frequent targets for a given set of predefined anchors in the set of input samples (FASTA/FASTQ files). It is a lightweight tool working in a streaming fashion. Note that many other choices in this procedure are possible, including the use of multiple targets or their quantification, different sequence or statistical embedding models, all a subject of further work.

We used FLASH and FAFQ Filter to predict resistance on genomes after fitting a balanced model on raw read data processed by SPLASH, so that under random guessing each sample has a 50/50 chance of classification into each category. We first trained a FLASH model on raw reads using antimicrobial phenotypes collapsed into binary categories: Intermediate/Resistant (I/R) versus Susceptible (S). Using *E. faecium* and *E. coli* genomes from (Hyun et al. 2023) we first identified any common isolates (those sharing a BioSample identifier) and dropped them from the data. We then used the tool FAFQ filter (see above) to extract the anchor clusters deemed significant by the FLASH model and following Step 2 used the predefined list of anchors from the FLASH run, formatting them and embedding them separately in the nucleotide language model. We then used these genome embeddings as the test set to predict resistance (Fig. 5A). We did this using clusters defined using the amino acid clustering approach as we hypothesized it may allow for better identification of anchors with more divergence from the original training data. The same analysis followed for the nanopore *E. faecium* reads from (Islam et al. 2023) (Supp. File S1).

### Viral analyses – H5N1

The viral analysis proceeded following the same methods as the other analyses except we further filtered the original anchors with any 12mer that was present in cattle (bosTaur9) or chicken (galGal6) genomes (to avoid calling host genetic data in the analysis). We then predicted on several metadata categories, including host, and visualized the results in the same manner as above (Supp. File S7).

### Bacteria-virus infection and interaction

We analyzed a study of 148 clinical isolates of Vibrio cholerae circulating in Bangladesh during the years 2016-2019 (LeGault et al. 2021). 44 ICP1 phage sequences isolated from water sources were also assembled and provided as orthogonal to sequenced isolates. Resistance to phages were experimentally found to fluctuate by year. We tested if FLASH could identify which phages were year-matched to circulating strains by taking all pairs of strains and phages and predicting the binary phenotype: were these two collected in the same year. First, we used *V. Cholera* sequences to predict collection year; in parallel, we identified all clusters of sequences in the phage genomes. We then incorporated the interactions between all phage and all marginal bacteria features by taking the rowwise dot product between the embeddings for each feature cluster. Using the matrix of marginal features and interactions, we then predicted accuracy using a grouped elastic net using 1) only the marginal features and 2) using the marginal features and the interactions, treating each interaction as its own group.

We followed the same methods for analyzing the data on co-infection above. Here, we marginally predicted bacteria on each phage and phage on each bacteria to identify a set of initial features. We then created the extended matrix of infection or not where each bacteria-phage was treated as a sample. (For phage we included the marginal features as well as the next 200 features to increase the space on which we were testing for interactions). Again, we predicted on the marginal grouped embeddings as features then on the matrix composed of those features plus their interaction dot products. In both cases, we used blast and blastx to annotate the significant features driving the predictions.

### Software used

FLASH is currently implemented as a Snakemake pipeline (version=9.6) (Mölder et al. 2021). Plots are generated using ggplot2 (Wickham 2016). Additional plotting was done using ComplexHeatmap (Gu 2022) and ggmsa (Zhou et al. 2022). Alpha fold structures are used directly from their database as well as through Alpha Fill for checking binding pockets (Hekkelman et al. 2023) and AF2Complex for looking at assemblies of multiple proteins (Gao et al. 2022). Additionally, ChimeraX was used to aid in visualization of the target regions in some protein structures (Meng et al. 2023). Foldseek was used to identify proteins with similar structures (van Kempen et al. 2024).

