## Supplementary material for "FunctionaL Assigning Sequence Homing (FLASH) maps phenotype to sequence with deep and machine learning": Combined Supplemental Info

#### Supplemental Material

##### Supplemental Methods

**A. Analyzing unblasted reads and predicted proteins in *C. tropicalis.***

##### Supplemental Tables

**Table S1.** Datasets analyzed using FLASH.

**Table S2.** Accuracies from other studies.

**Table S3.** Timing  and resource usage of FLASH Snakemake pipeline.

**Table S4.** Data used to train nucleotide language model.

##### Supplemental Figures

**Fig S1.** Itraconazole resistance in *Candida tropicalis* is driven by several correlated feature clusters including unannotated and hypothetical proteins.

**Fig S2.** Comparisons of clustering choices made when testing FLASH.

**Fig S3.** 8mers perform comparably to 27mers in *E. faecium* and *E. coli* but offer the opportunity for discovery of different genetic features.

**Fig S4.** In general, the most abundant target represents nearly 100% of each sample's variation, with the exception of *C. tropicalis* which has a slight decrease below 100%.

**Fig S5.** Hyena DNA LLM pre-trained on bacterial kmers performs similarly to a hyena model previously trained on hg38.

**Fig S6.** SATC (Sample Anchor Target Count) Utilities can be used to process SPLASH output or produce inputs for analyses without running SPLASH.

**Fig S7.** UMAP clustering of *S. pneumoniae* and *K. pneumoniae*.

**Fig S8.** FLASH discovers shared signatures of antibiotic resistance pathways across different bacterial species.

#### Supplemental Files

**File S1.** Prediction results in vancomycin-resistant *E. faecium* sequenced via nanopore and trained on held-out short reads.

**File S2.** All BLAST (protein and nucleotide) summaries by cluster for each metadata category across species.

**File S3.** Top 10 clusters and sequences for each metadata category across bacteria.

**File S4.** Cross-species sets of proteins predictive for antibiotic resistance across six different species.

**File S5.** After removing missing clusters, All BLAST (protein and nucleotide) summaries by cluster for each metadata category across species.

**File S6.** All fungal results.

**File S7.** H5N1 results after removing cattle and chicken genome contamination.

**File S8.** Predictors and accuracies for both *Vibrio*-phage studies.

**File S9.** All accessions used for analyses.

**File S10.** All MegaRes and Interpro SUPERFAMILY annotations for *E. faecium, E. coli, S. pneumoniae, K. pneumoniae, M. tuberculosis,* and *M. abscessus.*

**File S11.** All aggregated data for 8mer analyses of *E. faecium*, *E. coli* and Y1000 Sacchoromycotina data.

#### Supplemental Methods

**A. Analyzing unblasted reads and predicted proteins in *C. tropicalis*.**

In order to explore unblasted or hypothetical clusters in *C. tropicalis* we analyze long reads, searching for only reads which contain the sequence of an anchor we are interested in. There are seven accessions from (Fan et al. 2023) that have had PacBio sequencing done: SRR23943247, SRR23943248, SRR23943253, SRR23943252, SRR23943251, SRR23943250, and SRR23943249. The anchors we are interested in from each antifungal are laid out below:

- Fluconazole's third cluster doesn't blast (Cluster 3296). We use anchor ATCAATCCAACGGTTTTACTGTTTATG
- Posaconazole's second cluster doesn't blast (Cluster 7444). We use anchor TGGATGTCCAAGTGATCAATATCTCGT
- Itraconazole first cluster doesn't blast to our DBs (Cluster 2584). We use anchor AAAAGCCAGTGTTGCACCTCGTCAGTT
- Voriconazole's first cluster is hypothetical. We are interested in finding out more about it. We use a truncated anchor from Cluster 12142: GCTGCTGAATCTAGTGCTCCTG.

We use grep to extract all reads which contain these anchors from the long reads. We then independently BLAST (paramaters: -db core_nt -evalue 0.001 -task blastn -dust no -word_size 28 -reward 1 -penalty -2 -max_target_seqs 4) the resulting files for each of these four antifungals. By aggregating the number of reads on which different BLAST hits appear, we see common genes that are present near these clusters. In Fluconazole, there are **82** total long reads with the anchor. They BLAST to several hypothetical proteins given below:

```
69 Candida tropicalis MYA-3404 hypothetical protein (CTRG_03955), partial mRNA
65 Candida tropicalis MYA-3404 hypothetical protein (CTRG_03956), partial mRNA
52 Candida tropicalis MYA-3404 conserved hypothetical protein (CTRG_03953), partial mRNA
40 Candida tropicalis MYA-3404 hypothetical protein (CTRG_03958), partial mRNA
31 Candida tropicalis MYA-3404 carrier protein YMC1, mitochondrial precursor (CTRG_03954), partial mRNA
22 Candida tropicalis MYA-3404 glycogen debranching enzyme (CTRG_03951), partial mRNA
16 Candida tropicalis MYA-3404 hypothetical protein (CTRG_03959), partial mRNA
11 Candida tropicalis MYA-3404 chorismate mutase (CTRG_03957), partial mRNA
6 Candida tropicalis MYA-3404 hypothetical protein (CTRG_03960), partial mRNA
4 Candida tropicalis MYA-3404 conserved hypothetical protein (CTRG_03961), partial mRNA
4 Candida tropicalis MYA-3404 conserved hypothetical protein (CTRG_03952), partial mRNA
2 Candida tropicalis MYA-3404 hypothetical protein (CTRG_03963), partial mRNA
1 Sinorhizobium medicae strain T10 plasmid unnamed1
1 PREDICTED: Pangasianodon hypophthalmus vesicle amine transport 1 (vat1), mRNA
1 Candida tropicalis MYA-3404 hypothetical protein (CTRG_03950), partial mRNA
1 [Candida] subhashii YMC2 (J8A68_001144), partial mRNA
```

When grabbing one of these reads and using BLAT against the *C. tropicalis* reference genome, GCF_000006335.3, there are large deletions:

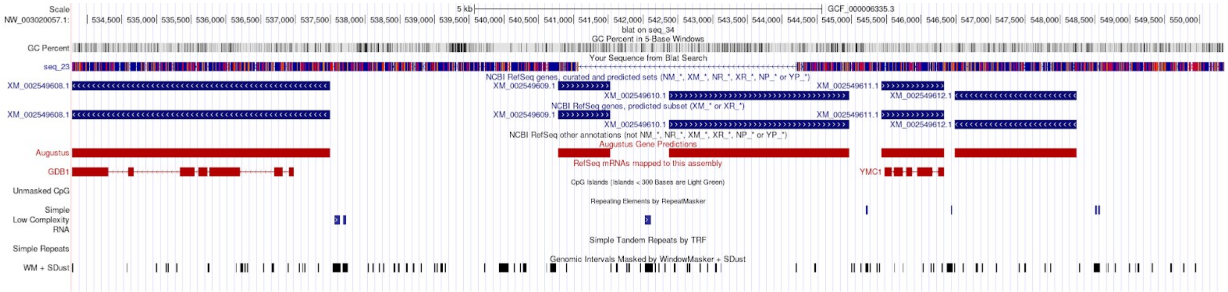

In Posaconazole, there are 88 long reads with the anchor we are interested in. They BLAST to several hypothetical proteins given below:

```
65 Candida tropicalis MYA-3404 conserved hypothetical protein (CTRG_04444), partial mRNA
62 Candida tropicalis MYA-3404 conserved hypothetical protein (CTRG_04445), partial mRNA
59 Candida tropicalis MYA-3404 conserved hypothetical protein (CTRG_04446), partial mRNA
40 Candida tropicalis MYA-3404 protein ABC1, mitochondrial precursor (CTRG_04443), partial mRNA
21 Candida tropicalis MYA-3404 predicted protein (CTRG_04447), partial mRNA
13 Candida albicans SC5314 ribosomal 40S subunit protein S2 (RPS21), partial mRNA
12 Candida tropicalis MYA-3404 protein RHO4 (CTRG_04441), partial mRNA
8 Candida tropicalis MYA-3404 conserved hypothetical protein (CTRG_04440), partial mRNA
5 Candida dubliniensis CD36 40S ribosomal protein S2 (CD36_01360), partial mRNA
1 Wickerhamomyces ciferrii 40S ribosomal protein uS5 (RPS2), partial mRNA
1 Spathaspora passalidarum NRRL Y-27907 40S ribosomal protein S2 (SPAPADRAFT_57745), mRNA
1 Saccharomycodes ludwigii uncharacterized protein (SCDLUD_001000), partial mRNA
1 Pungitius pungitius genome assembly, segment: ctg7180000009036
1 PREDICTED: Scylla paramamosain uncharacterized LOC135106467 (LOC135106467), transcript variant X6, ncRNA
1 PREDICTED: Pangasianodon hypophthalmus uncharacterized protein LOC619199 homolog (zgc:112334), mRNA
1 PREDICTED: Microplitis demolitor uncharacterized LOC103580284 (LOC103580284), transcript variant X6, mRNA
1 PREDICTED: Megalobrama amblycephala ADP-ribosylation factor-like 6 (arl6), transcript variant X3, mRNA
1 PREDICTED: Anser cygnoides trichohyalin-like (LOC136790603), mRNA
1 Populus EST from leave
1 Monocercomonoides exilis putative ubiquitin carboxyl-terminal hydrolase FAF-X (MONOS_3137), partial mRNA
1 Escherichia coli strain SX45 plasmid pSX45-3-mcr-1, complete sequence
1 Escherichia coli strain OXEC-27 plasmid unnamed
1 Candida tropicalis MYA-3404 superoxide dismutase, mitochondrial precursor (CTRG_04448), partial mRNA
1 Candida tropicalis MYA-3404 predicted protein (CTRG_04453), partial mRNA
1 Candida tropicalis MYA-3404 conserved hypothetical protein (CTRG_04452), partial mRNA
1 Candida tropicalis MYA-3404 conserved hypothetical protein (CTRG_04451), partial mRNA
1 Candida pseudojiufengensis uncharacterized protein (KGF55_000655), partial mRNA
1 Candida pseudojiufengensis RPS2 (KGF55_000652), partial mRNA
1 Candida oxycetoniae RPS2 (KGF56_001857), partial mRNA
1 Candida metapsilosis 40S ribosomal protein uS5 (I9W82_001140), partial mRNA
1 Candida margitis RPS2 (CANMA_003767), partial mRNA
1 Candida jiufengensis NAB2 (KGF54_001264), partial mRNA
1 Candida albicans SC5314 uncharacterized protein (CAALFM_C101510WA), partial mRNA
```

Similarly, using BLAT to map one of these long reads against the C. tropicalis reference genome, we see evidence for at least one potential large deletion.

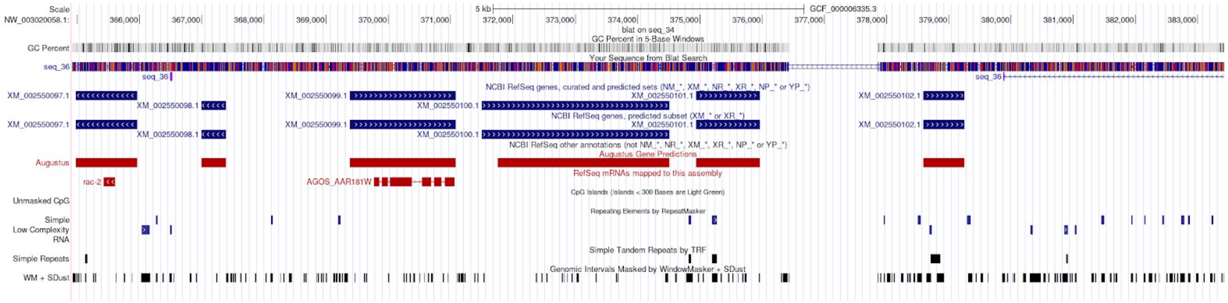


In Itraconazole, there are 154 long reads with the anchor. They BLAST to several hypothetical proteins given below:

```
105 Candida tropicalis MYA-3404 predicted protein (CTRG_00296), partial mRNA
104 Candida tropicalis MYA-3404 hypothetical protein (CTRG_00294), partial mRNA
100 Candida tropicalis MYA-3404 hypothetical protein (CTRG_00301), partial mRNA
90 Candida tropicalis MYA-3404 predicted protein (CTRG_00297), partial mRNA
```

```
82 Candida tropicalis MYA-3404 predicted protein (CTRG_00295), partial mRNA
49 Candida tropicalis MYA-3404 conserved hypothetical protein (CTRG_00303), partial mRNA
20 Candida tropicalis MYA-3404 predicted protein (CTRG_00291), partial mRNA
14 Candida tropicalis MYA-3404 predicted protein (CTRG_00298), partial mRNA
13 Candida tropicalis MYA-3404 predicted protein (CTRG_00299), partial mRNA
8 Candida tropicalis MYA-3404 predicted protein (CTRG_00302), partial mRNA
7 Candida tropicalis MYA-3404 predicted protein (CTRG_01926), partial mRNA
6 Candida tropicalis MYA-3404 predicted protein (CTRG_00304), partial mRNA
2 Candida tropicalis MYA-3404 predicted protein (CTRG_00292), partial mRNA
2 Candida tropicalis MYA-3404 conserved hypothetical protein (CTRG_00290), partial mRNA
1 PREDICTED: Vulpes lagopus basic proline-rich protein-like (LOC121501784), partial mRNA
1 PREDICTED: Salvelinus namaycush basic proline-rich protein-like (LOC120046039), partial mRNA
1 PREDICTED: Salvelinus namaycush basic proline-rich protein-like (LOC120022283), partial mRNA
1 Human herpesvirus 1 strain 5-5-2, partial genome
1 Candida tropicalis MYA-3404 predicted protein (CTRG_00300), partial mRNA
1 Candida tropicalis MYA-3404 opaque-specific ABC transporter CDR3 (CTRG_00286), partial mRNA
1 Candida tropicalis MYA-3404 conserved hypothetical protein (CTRG_00287), partial mRNA
1 Candida albicans SC5314 uncharacterized protein (CAALFM_C404010WA), partial mRNA
```

We also take one of these long reads and BLAT it, showing interesting patterns and poor mapping:

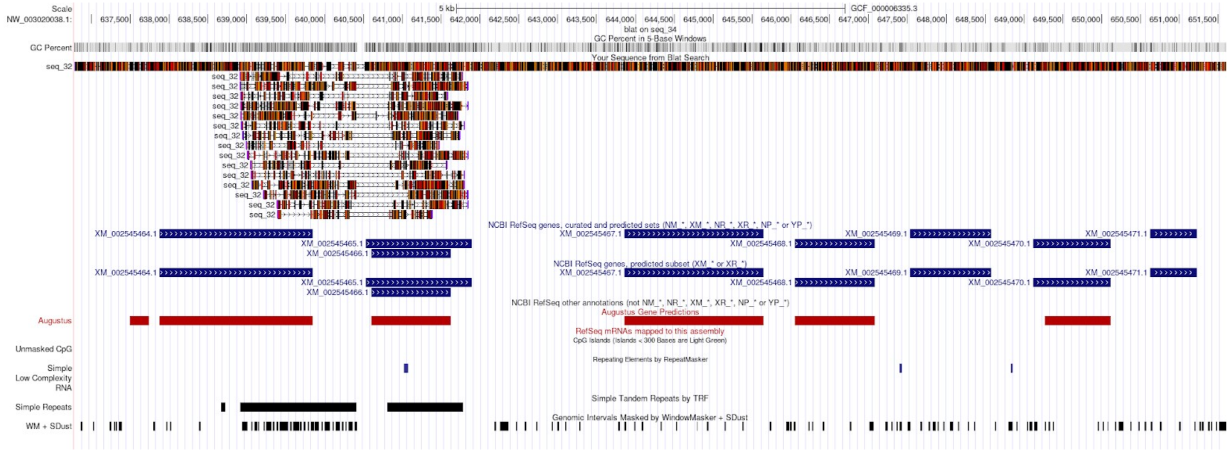


For Voriconazole, there are 116 reads with the truncated anchor given above. Counting their occurrences on long reads, we get the following breakdown of reads with BLAST hits to genes:

```
82 Candida tropicalis MYA-3404 predicted protein (CTRG_04841), partial mRNA
61 Candida tropicalis MYA-3404 predicted protein (CTRG_04840), partial mRNA
61 Candida tropicalis MYA-3404 hypothetical protein (CTRG_04839), partial mRNA
51 Candida tropicalis MYA-3404 predicted protein (CTRG_04842), partial mRNA
19 Candida tropicalis MYA-3404 conserved hypothetical protein (CTRG_04838), partial mRNA
17 Candida tropicalis MYA-3404 predicted protein (CTRG_06242), partial mRNA
14 PREDICTED: Culex pipiens pallens uncharacterized LOC128092361 (LOC128092361), partial mRNA
14 Hahella chejuensis KCTC 2396, complete genome
14 Candida tropicalis MYA-3404 hypothetical protein (CTRG_04948), partial mRNA
12 Candida tropicalis MYA-3404 predicted protein (CTRG_06243), partial mRNA
 7 Culex quinquefasciatus cell surface glycoprotein 1-like (LOC119771006), mRNA
 6 Suhomyces tanzawaensis NRRL Y-17324 uncharacterized protein (CANTADRAFT_7189), partial mRNA
 6 Candida tropicalis MYA-3404 predicted protein (CTRG_04837), partial mRNA
 3 PREDICTED: Sabethes cyaneus protein TsetseEP-like (LOC128739554), partial mRNA
 3 PREDICTED: Culex pipiens pallens uncharacterized LOC128092780 (LOC128092780), partial mRNA
 3 Culex quinquefasciatus vegetative cell wall protein gp1-like (LOC119766921), mRNA
 2 Culex quinquefasciatus vegetative cell wall protein gp1-like (LOC119765508), mRNA
 2 Candida tropicalis MYA-3404 conserved hypothetical protein (CTRG_04279), partial mRNA
 2 [Candida] subhashii PHO87 (J8A68_005413), partial mRNA
```

```
2 Candida dubliniensis CD36 inorganic phosphate transporter PHO87, putative (CD36_05600), partial mRNA
1 Urechis unicinctus clone scaffold A homeobox protein region genomic sequence
1 PREDICTED: Prionailurus bengalensis basic proline-rich protein-like (LOC122466907), partial mRNA
1 PREDICTED: Macaca thibetana thibetana basic proline-rich protein-like (LOC126950815), partial mRNA
1 PREDICTED: Bombyx mori uncharacterized LOC134200883 (LOC134200883), transcript variant X3, ncRNA
1 Kaburagia rhusicola voucher ZMIOZ15700 mitochondrion, partial genome
1 Culex quinquefasciatus vegetative cell wall protein gp1-like (LOC119769315), mRNA
1 Candida tropicalis MYA-3404 conserved hypothetical protein (CTRG_04278), partial mRNA
1 Candida albicans SC5314 SPX domain-containing inorganic phosphate transporter (PHO87), partial mRNA
1 3_Tms_b3v08
```

We also use BLAT to explore one of the sequences in this cluster and observe the following indicating possible rearrangements or poor mappability to the reference.

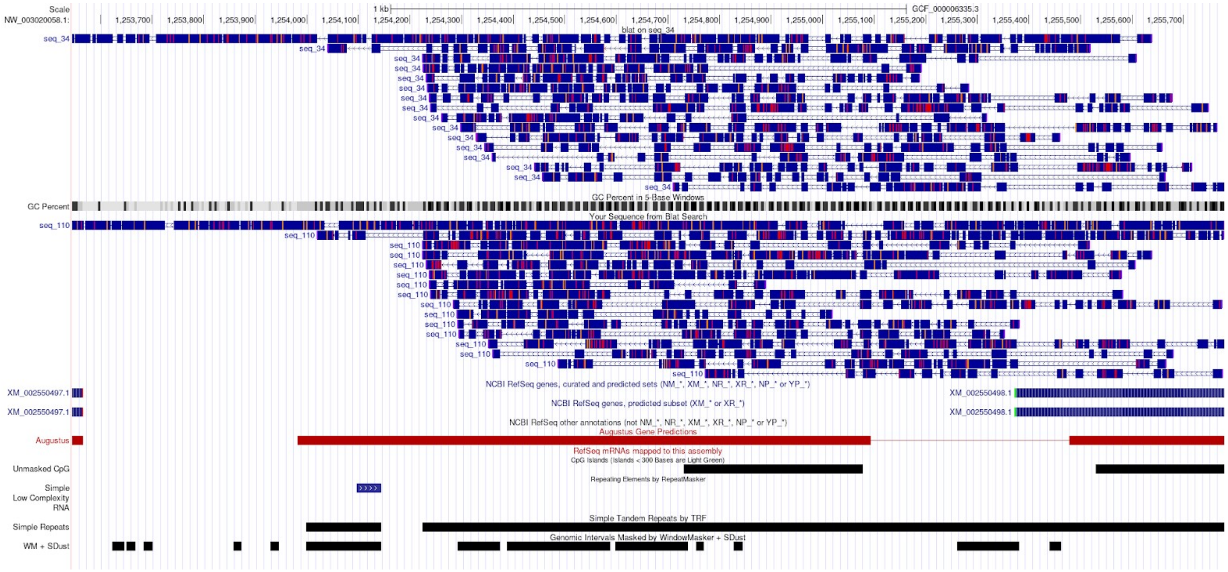


A set of commonly represented genes on these long reads, CTRG_06242 and CTRG_06243, hypothetical proteins, had the unusual quality of having multiple BLAST hits to portions of the same gene, indicative of structural rearrangement or a high degree of polymorphism (Supp. Methods). CTRG_06243 is annotated as containing a FLO11-domain, and a signal peptide. In *S. cerevisiae*, FLO11 regulates morphological developmental switches and has been reported to have mutagenic processes that create cell surface variation (Halme et al. 2004), consistent with large target diversity. The same Flocculin type 3 repeat– though this repeat was not assigned to a single gene– was the most important predictor for voriconazole resistance identified in a previous study (Harrison et al. 2025). The FLO11 domain has been reported to undergo structural rearrangements (Fidalgo et al. 2006). Together with the high correlation between voriconazole and targets of the anchor in CTRG_04948 identity, as well as the high target diversity in CTRG_04948 anchors, we predict a new link between CTRG_04948 variation and virulence.

The anchor technically blasts into CTRG_04948 and 04842 but it also blasts just downstream of this FLO11 gene discussed:

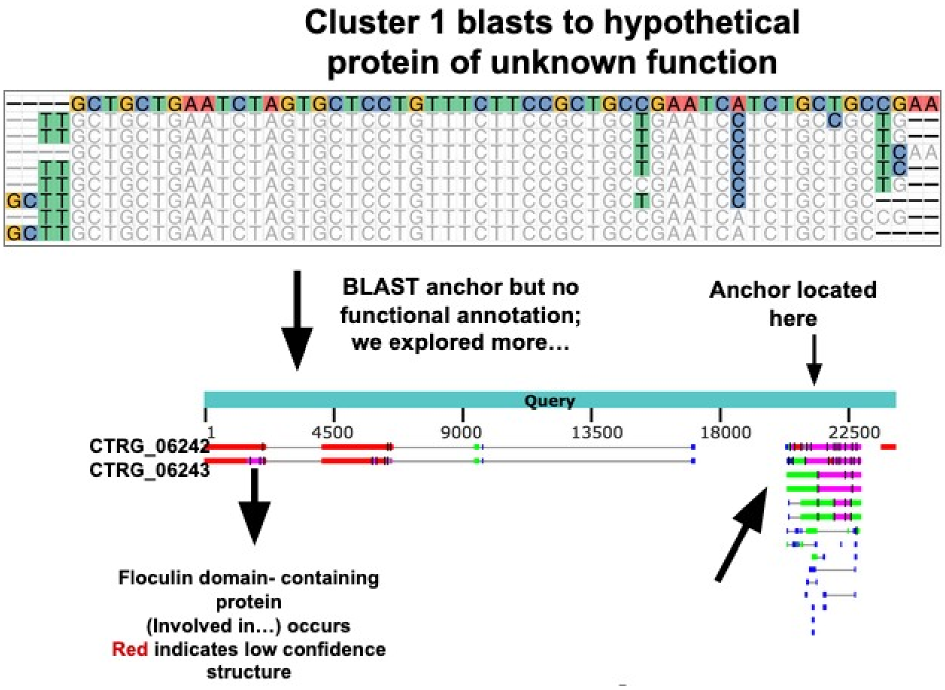


Figure 4D Illustrates an analysis further exploring the relationship between CTRG_mmm and drug resistance. We tiled genes CRTG_04948 and CTRG_04842 for all 27mer anchors and then analyzed their Sample Anchor Target Count tables as heatmaps. Anchor GCTGCTGAATCTAGTGCTCCTG shows a relationship between the unique number of downstream targets and the resistance status of the fungal samples.

### Supplemental Tables

**Table S1. Datasets analyzed using FLASH.** Datasets analyzed when running FLASH. SPLASH was run on R1 samples for the accessions identified in Supp. File S8. The translation tables and taxon filters used are indicated as well as any information about compiling the data.

| ***Category*** | ***Data*** | **metadata** | **samples** | **Bio Project** | **Paper(s)** | **Additional Metadata / Notes** | **Translation Table** | **Taxon Filter Used** |
|---|---|---|---|---|---|---|---|---|
| ***Bacterial*** | ***E. coli*** | Antibiotic resistance | 7,057 | See notes | (Mets and Morin 2024) | Dataset compiled from many different sources. See paper for details. | 11 | 561 |
| ***Bacterial*** | ***E. faecium*** | Antibiotic resistance | 4,382 | See notes | (Coll et al. 2024) | Dataset compiled from many different sources. See paper for details. | 11 | 1350 |
| ***Bacterial*** | ***K. pneumoniae*** | Sequence type and antibiotic resistance | 1374 | PRJEB 42462 | (Heinz et al. 2024) | - | 11 | 570 |
| ***Bacterial*** | ***S. aureus*** | Carriage of pyomyositis | 518 | PRJNA 418899 | (Young et al. 2019) | - | 11 | 1279 |
| ***Bacterial*** | ***S. pneumoniae*** | GPSC cluster / antibiotic resistance phenotypes | 3232 | ERP00 1505 | (Lo et al. 2019) | Data from Global Pneumococcal Sequencing Project; subset from paper<br><br>1,494 Gbases<br><br>1h (96-core machine), $10 computing cost for SPLASH | 11 | 1301 |
| ***Bacterial*** | ***S. pyogenes (Strep A)*** | Serotype / emm type | 1515 | SRP25 3833 | (Southon et al. 2020) | - | 11 | 1301 |
| ***Bacterial*** | ***S. agalactiae (Group B strep)*** | Sequence type / disease onset / tissue | 1,512 | ERP01 5737 | (Chaguza et al. 2022) | - | 11 | 1301 |
| ***Bacterial*** | ***M. tuberculosis*** | PZA resistance | 10,725 | See notes | (Kim et al. 2023) | Compiled across various studies. Refer to Table S1 from the original paper. | 11 | 1773 |
| ***Bacterial*** | ***M. abscessus*** | Computation ally characterized amikacin resistance | 1,937 | See notes | See notes | Compiled from the SRA after searching for *M. abscessus* and quantifying resistance via presence of known mutations | 11 | 670516 |
| ***Fungal*** | ***Aspergillus fumigatus*** | Triazole resistance and mating type | 187 | SRP26 1459 | (Etienne et al. 2021) | - | 4 | 5052 |

| | | | | | | | | |
|---|---|---|---|---|---|---|---|---|
| ***Fungal*** | ***Candida tropicalis*** | Azole resistances | 459 | SRP428185 | (Fan et al. 2023) | Use only new isolates sequenced in the study | 12 | 147537 |
| ***Fungal*** | ***Candidomyza auris*** | Azole resistances | 181 | PRJNA1003896 | (Misas et al. 2024) | - | 12 | 498019 |
| ***Fungal*** | ***Y1000 / Saccharomycotina*** | Azole resistance | 1088 | PRJNA736342 | (Opulente et al. 2024; Harrison et al. 2025; Shen et al. 2018) | - | 1 | 147537 |
| ***Viral*** | ***H5N1 in Birds and Dairy Cattle*** | Host/collection source/collection date | 1227 | SRP503016 | (Nguyen et al. 2025) | https://www.ncbi.nlm.nih.gov/bioproject/PRJNA1102327 *Use isolates released before 8-11-24* | 1 | 10239 (after filtering for contamination) |
| ***Phage interactions*** | ***V. cholerae time series*** | Year of sampling | 244 bacteria 44 phage | & See notes | (LeGault et al. 2021) | *Phage sequences were drawn directly from genomes and assembled without SPLASH | 11* | 0 |
| ***Phage - Interactions*** | ***Vibrio-phage co-infection*** | Co-infection | 256 bacteria 239 phage | | (Kauffman et al. 2022) | *Phage sequences were translated using table 1 | 11* | 0 |

**Table S2. Accuracies from other studies.** FLASH accuracy was compared against accuracies from two other studies which report accuracy. For Y1000, the accuracies are compiled from tables and figures from the corresponding studies. Accuracies here are calculated on a balanced test set and use 90% of individuals for training. We do the same for our comparisons. The Lancet accuracies are calculated across all of the data using presence / absence of genes from a genotype database. We compare our standard accuracies to these.

| **Study** | **Metadata Category** | **Accuracy from Paper** |
|---|---|---:|
| Y1000 (Harrison et al.) | Fluconazole_resistance | 0.749 |
| Y1000 (Harrison et al.) | Voriconazole_resistance | 0.659 |
| Y1000 (Harrison et al.) | Terbinafin_resistance | 0.675 |
| Y1000 (Harrison et al.) | Caspofungi_resistance | 0.668 |
| Y1000 (Harrison et al.) | Amphotericin_B_resistance | 0.638 |
| Y1000 (Harrison et al.) | Itraconazole_resistance | 0.531 |
| Y1000 (Harrison et al.) | Posaconazole_resistance | 0.648 |
| *E. faecium* (Lancet) | ampicillin_RIS | 0.9533 |
| *E. faecium* (Lancet) | ciprofloxacin_RIS | 0.9866 |
| *E. faecium* (Lancet) | clindamycin_RIS | 0.9022 |
| *E. faecium* (Lancet) | daptomycin_RIS | 0.88 |
| *E. faecium* (Lancet) | erythromycin_RIS | 0.7914 |
| *E. faecium* (Lancet) | gentamicin_RIS | 0.9234 |
| *E. faecium* (Lancet) | kanamycin_RIS | 0.9106 |
| *E. faecium* (Lancet) | linezolid_RIS | 0.8812 |

**Table S3. Timing and resource usage of FLASH Snakemake pipeline.** FLASH timings (post-SPLASH analysis) for 12 of the datasets mentioned above. Time measures time to prediction and not annotation. Datasets are benchmarked using normalized nucleotide embeddings and nucleotide-based shift distance clustering and capped at 20,000 features per analysis.

| **Dataset** | **Num Samples** | **Num Metadata categories** | **Total Time (H)** | **Max Memory PSS (Gb)** | **Max Cpus** |
|---|---|---|---|---|---|
| *S. agalactiae* (Group B Strep) | 1338 | 9 | 1.92 | 150.18 | 32 |
| H5N1 | 1226 | 10 | 0.77 | 78.27 | 32 |
| *A. fumigatus* | 187 | 22 | 0.71 | 20.84 | 32 |
| *C. tropicalis* | 459 | 31 | 1.14 | 44.03 | 32 |
| *E. coli* | 7026 | 51 | 6.59 | 484.91 | 32 |
| *E. faecium* | 4730 | 16 | 3.82 | 286.27 | 32 |
| K. pneumoniae | 1373 | 48 | 2.35 | 137.3 | 32 |
| *S. pneumoniae* | 3233 | 24 | 5.84 | 367.09 | 32 |
| *S. aureus* | 518 | 5 | 0.69 | 39.84 | 32 |
| *S. pyogenes* (Strep A) | 1515 | 11 | 1.76 | 145.08 | 32 |
| Y1000 (Sacchoromycotina) | 516 | 18 | 3.57 | 77.73 | 32 |
| *M. tuberculosis* | 10725 | 2 | 4.09 | 644.37 | 32 |

**Table S4. Data used to train nucleotide language model.** FLASH embeddings are generated using a pre-trained nucleotide language model built off of HyenaDNA. The data used to train this language model were pre-processed with SPLASH. Data size and speed/cost of SPLASH analysis are indicated below.

| **Data Source** | **Identifiers** | **Size of Data** | **Speed of SPLASH analysis** |
|---|---|---|---|
| bacteria2 | SRP348579, SRP325909, SRP372864, SRP139780, SRP139779, SRP094837 | 986 Gbases | 14h (32- and 64-core machines), $31 computing cost |
| bacteria3 | SRP131361, SRP301889, SRP254676, SRP457417, SRP303098, SRP186318, SRP325909, SRP262345, SRP273998 | 798 Gbases | 4h (32-core machine), $8 computing cost |
| pneumo* | 3233 runs from ERP001505, based on samples analyzed in https://pmc.ncbi.nlm.nih.gov/articles/PMC7641901/#ack3 | 1,494 Gbases | 1h (96-core machine), $10 computing cost |
| ERP001505 | ERP001505, 204 jobs; up to 200 runs per job | 16,309 Gbases | 177h (8-core machines), $103 computing cost |

#### Supplemental Figures

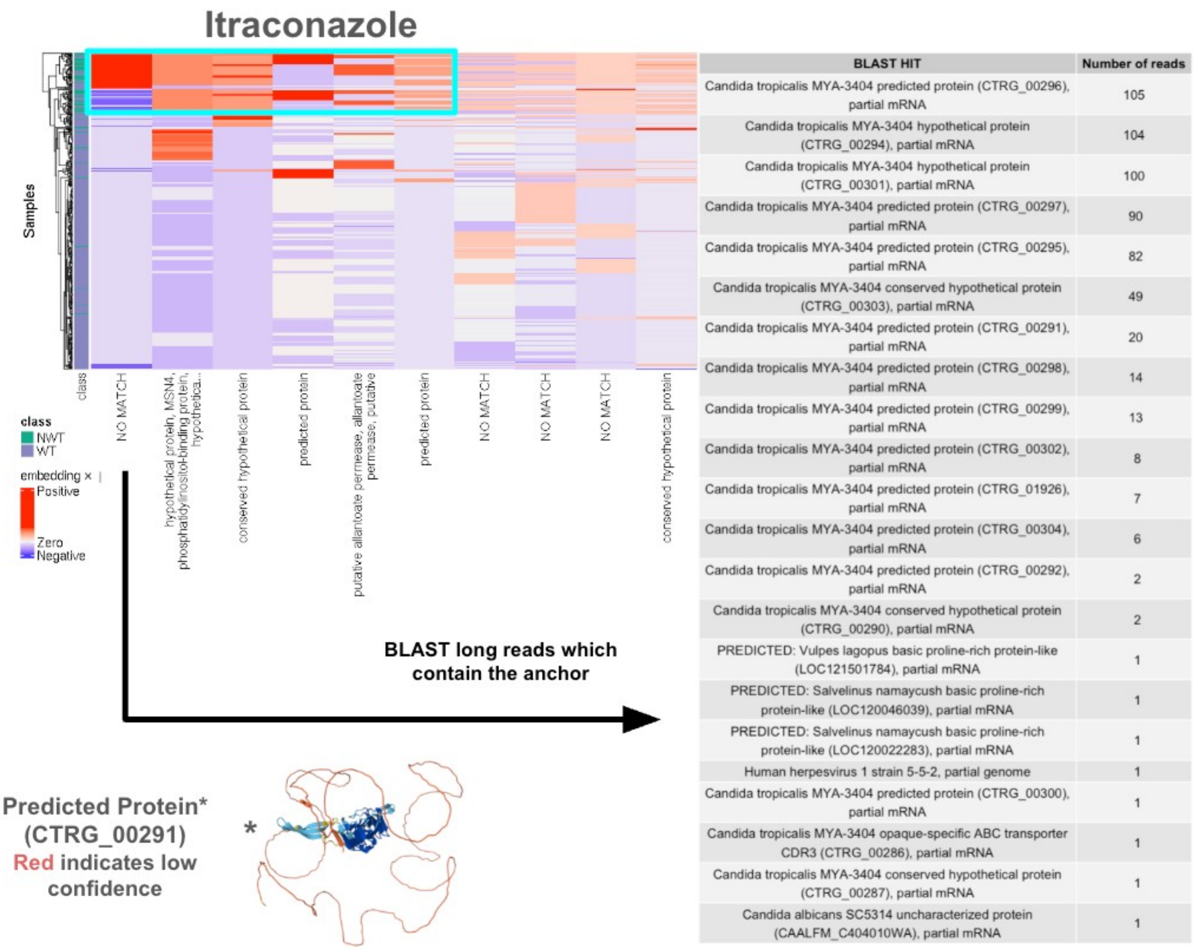


| BLAST HIT | Number of reads |
|---|---|
| Candida tropicalis MYA-3404 predicted protein (CTRG_00296), partial mRNA | 105 |
| Candida tropicalis MYA-3404 hypothetical protein (CTRG_00294), partial mRNA | 104 |
| Candida tropicalis MYA-3404 hypothetical protein (CTRG_00301), partial mRNA | 100 |
| Candida tropicalis MYA-3404 predicted protein (CTRG_00297), partial mRNA | 90 |
| Candida tropicalis MYA-3404 predicted protein (CTRG_00295), partial mRNA | 82 |
| Candida tropicalis MYA-3404 conserved hypothetical protein (CTRG_00303), partial mRNA | 49 |
| Candida tropicalis MYA-3404 predicted protein (CTRG_00291), partial mRNA | 20 |
| Candida tropicalis MYA-3404 predicted protein (CTRG_00298), partial mRNA | 14 |
| Candida tropicalis MYA-3404 predicted protein (CTRG_00299), partial mRNA | 13 |
| Candida tropicalis MYA-3404 predicted protein (CTRG_00302), partial mRNA | 8 |
| Candida tropicalis MYA-3404 predicted protein (CTRG_01926), partial mRNA | 7 |
| Candida tropicalis MYA-3404 predicted protein (CTRG_00304), partial mRNA | 6 |
| Candida tropicalis MYA-3404 predicted protein (CTRG_00292), partial mRNA | 2 |
| Candida tropicalis MYA-3404 conserved hypothetical protein (CTRG_00290), partial mRNA | 2 |
| PREDICTED: Vulpes lagopus basic proline-rich protein-like (LOC121501784), partial mRNA | 1 |
| PREDICTED: Salvelinus namaycush basic proline-rich protein-like (LOC120046039), partial mRNA | 1 |
| PREDICTED: Salvelinus namaycush basic proline-rich protein-like (LOC120022283), partial mRNA | 1 |
| Human herpesvirus 1 strain 5-5-2, partial genome | 1 |
| Candida tropicalis MYA-3404 predicted protein (CTRG_00300), partial mRNA | 1 |
| Candida tropicalis MYA-3404 opaque-specific ABC transporter CDR3 (CTRG_00286), partial mRNA | 1 |
| Candida tropicalis MYA-3404 conserved hypothetical protein (CTRG_00287), partial mRNA | 1 |
| Candida albicans SC5314 uncharacterized protein (CAALFM_C404010WA), partial mRNA | 1 |

**Fig S1. Itraconazole resistance in *Candida tropicalis* is driven by several correlated feature clusters including unannotated and hypothetical proteins.** Heatmap of cluster contributions to the prediction of Itraconazole resistance (left) and Voriconazole resistance (right) in C. tropicalis. The first predictive cluster for Itraconazole has no blast hit. Grabbing all pacbio reads and blasting them against C. tropicalis, there are many common hypothetical genes that show up (table). One, https://alphafold.ebi.ac.uk/entry/C5M2K2, is partially annotated as containing a hyphally-regulated cell wall protein. Much of alphafold's prediction of the structure is low confidence (orange regions).

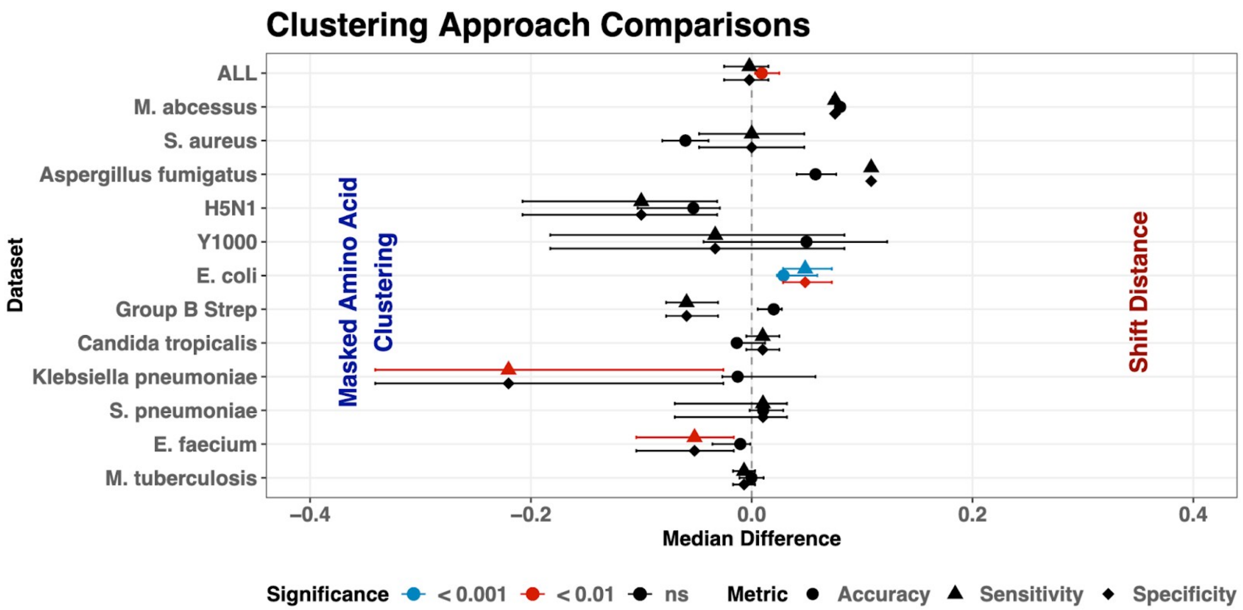


**Fig S2. Comparisons of clustering choices made when testing FLASH.** Wilcoxon signed rank test comparing accuracies across different cluster approaches (right). The pseudomedian and confidence intervals are reported for each study and all studies aggregated together. Shift Distance refers to using shift distance to define networks and then using disconnected components from these networks to define clusters (see methods). Masked Amino Acid clustering refers to translating sequences, masking them, and then clustering them if their masks match one another in either the nucleotide or amino acid space (see methods). The shiftDist-levFilter and masked aa cluster both perform similarly in accuracy across all data, though in the aggregate shift distance performs significantly better (with only a 0.9% increase in accuracy).

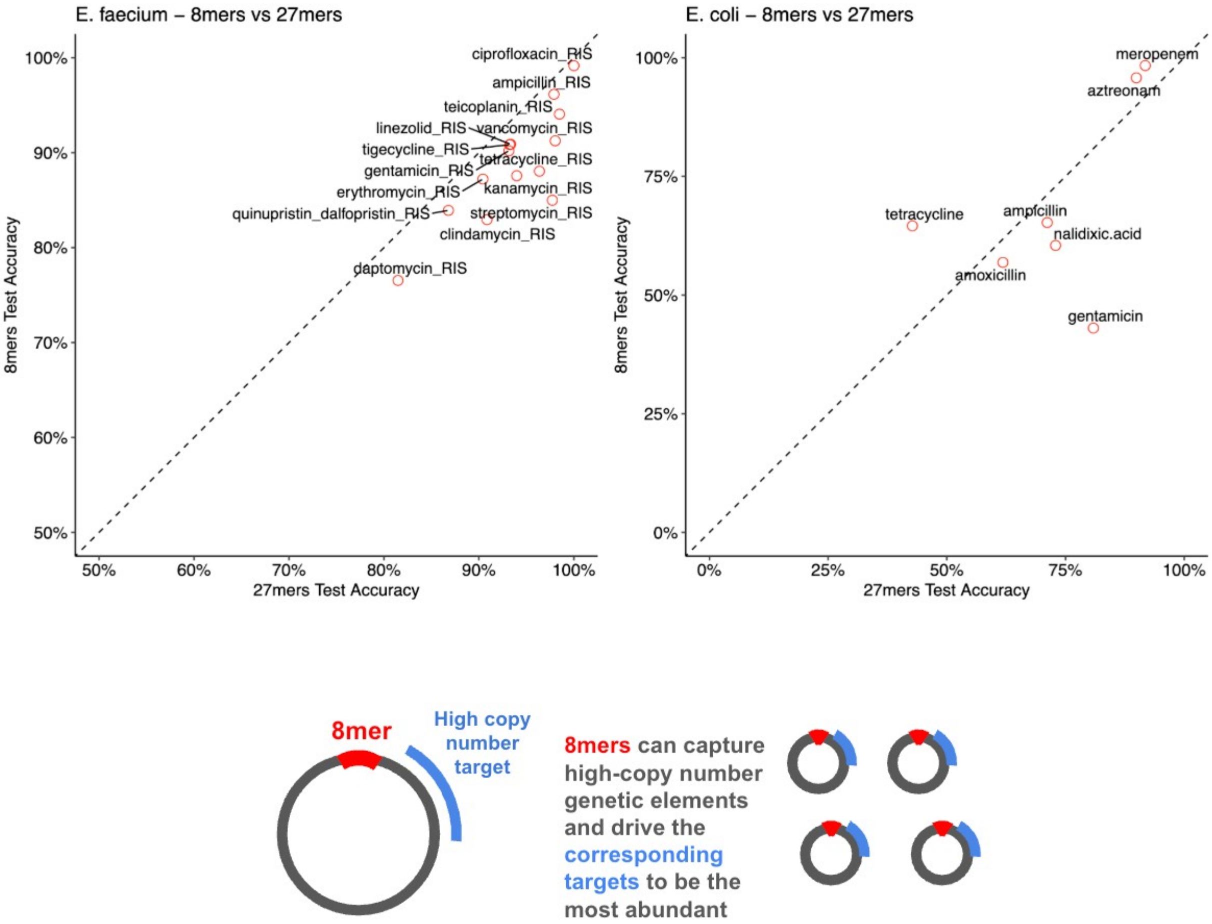


**Fig. S3. 8mers perform comparably to 27mers in *E. faecium* and *E. coli* but offer the opportunity for discovery of different genetic features.** Predictions made using hyena embeddings and one-hot encoding calculated on 8mers (with 31mer targets) vs 27mers (with 27mer targets) in antibiotic resistance data from *E faecium* (top left) and *E. coli* (top right). 8mers perform slightly better than 27mers for some metadata categories but performance is similar in general. For some specific cases, we hypothesize that 8mers can capture certain signatures missed by 27mers especially when it comes to higher copy number or identification of repeats (schematic).

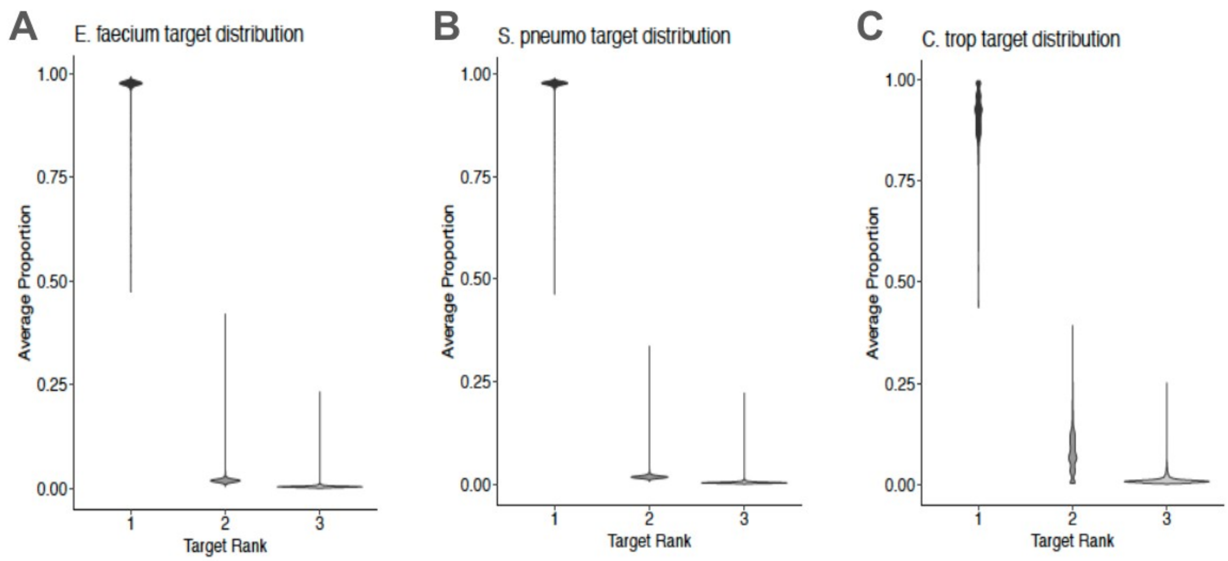


**Fig S4. In general, the most abundant target represents nearly 100% of each sample's variation, with the exception of *C. tropicalis* which has a slight decrease below 100%.** Average proportions of the top 1st, 2nd, and 3rd targets (by abundance) for each of the top effect size anchors called when using SPLASH and FLASH on (A) *Enterococcus faecium*, (B) *Streptococcus pneumoniae*, and (C) *Candida tropicalis*.

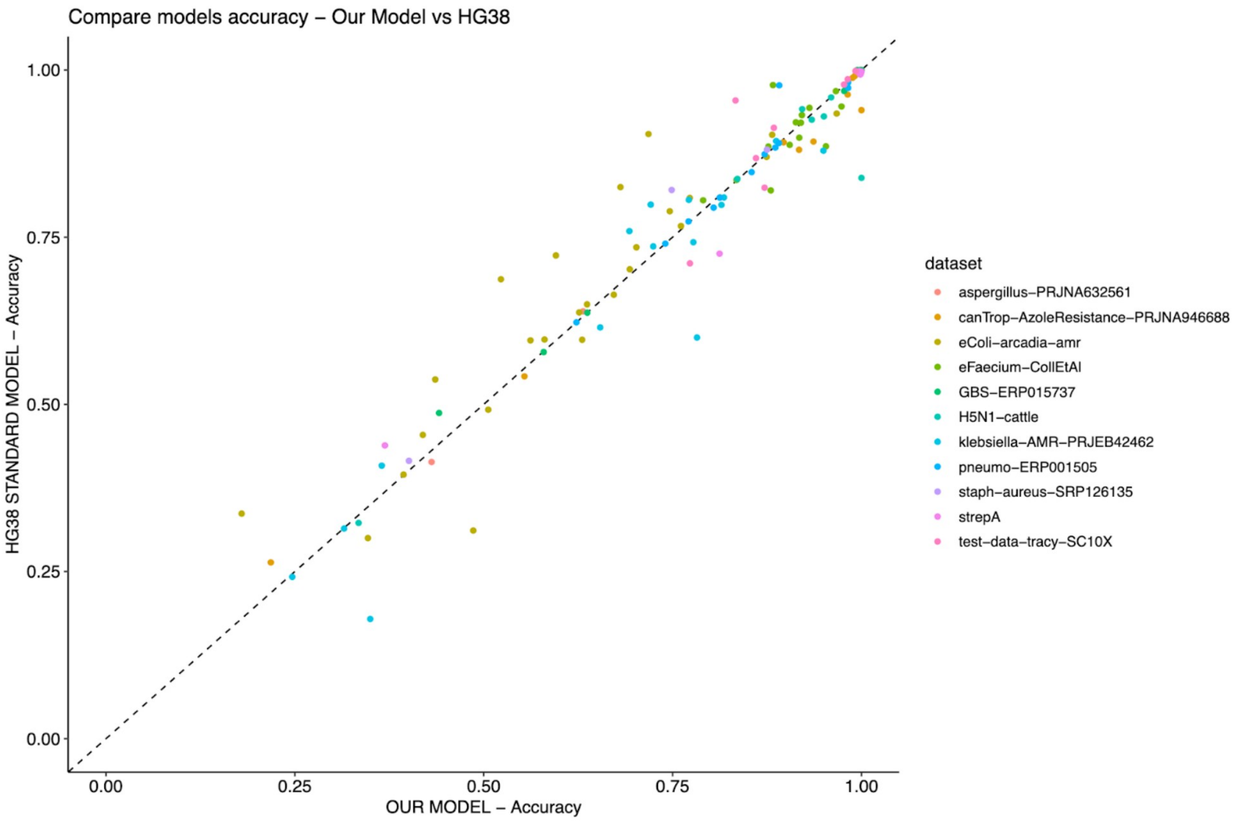


**Fig. S5. Hyena DNA LLM pre-trained on bacterial kmers performs similarly to a hyena model previously trained on hg38.** Plotting the accuracy across all datasets we explored was calculated in a model downloaded and already pre-trained (x axis) and a model we trained on data composed of anchors and targets collected from many different bacteria. The model we compare to can be found here: huggingface.co/LongSafari/hyenadna-medium-160k-seqlen

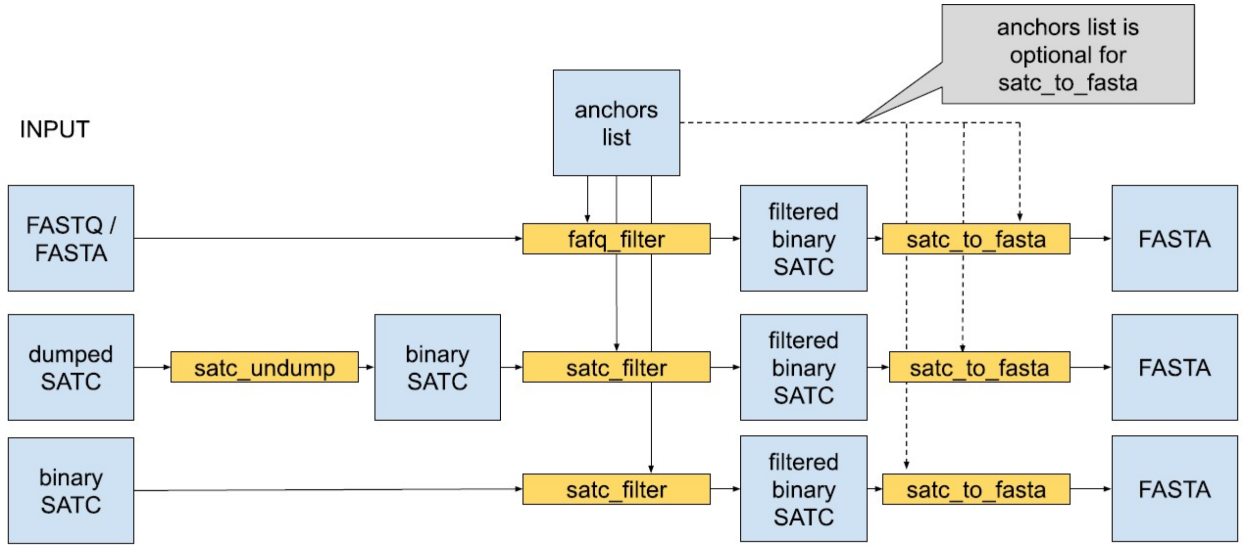


**Fig. S6. SATC (Sample Anchor Target Count) Utilities can be used to process SPLASH output or produce inputs for analyses without running SPLASH.** High level overview of SATC (Sample, Anchor, Target, Count) file helper utilities. Input can range from raw read files (FASTA/FASTQ) to SPLASH SATC output files. These utilities can be used to select anchors and abundant targets from hashed kmers calculated from fastq files (fafq_filter) or directly from satc files (satc_filter). The utility satc_to_fasta can also be used to unwrap a given sample's anchor and target variation into a sequence like those we are using for calculating embeddings in FLASH.

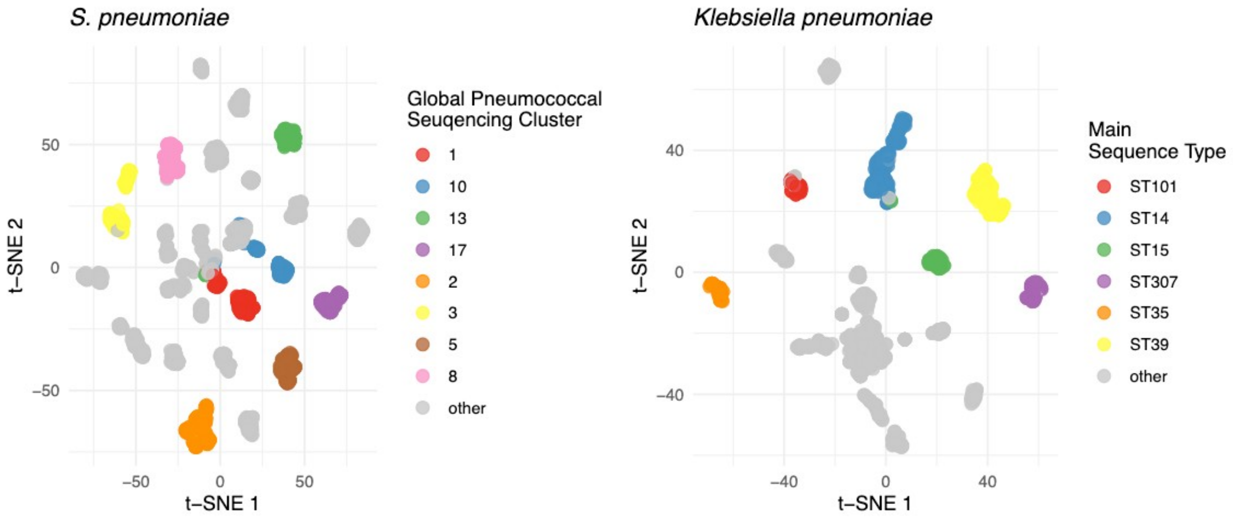


**Fig. S7. UMAP clustering of *S. pneumoniae* and *K. pneumoniae*.** t-SNE clustering of sequence variation in 1,338 samples of 3,233 samples of *S. pneumoniae* (left) and 1,313 samples of *K. pneumoniae* (right). Sequence variation is identified using SPLASH before these features are clustered and embedded in DNA Hyena (see methods). We cluster the matrix composed of samples (with >=30 samples) by one embedding dimension per cluster by first performing principal component analysis on this matrix and then visualizing the top 10 principal components with t-SNE. Colors here correspond only to the most abundant clusters per category (Global Pneumococcal Sequencing Clusters and Sequence Types). Adjusted rand indices indicate clusters which match labeled phenotypes: 0.827 for *S. pneumoniae*, and 0.887 for *K. pneumoniae.*

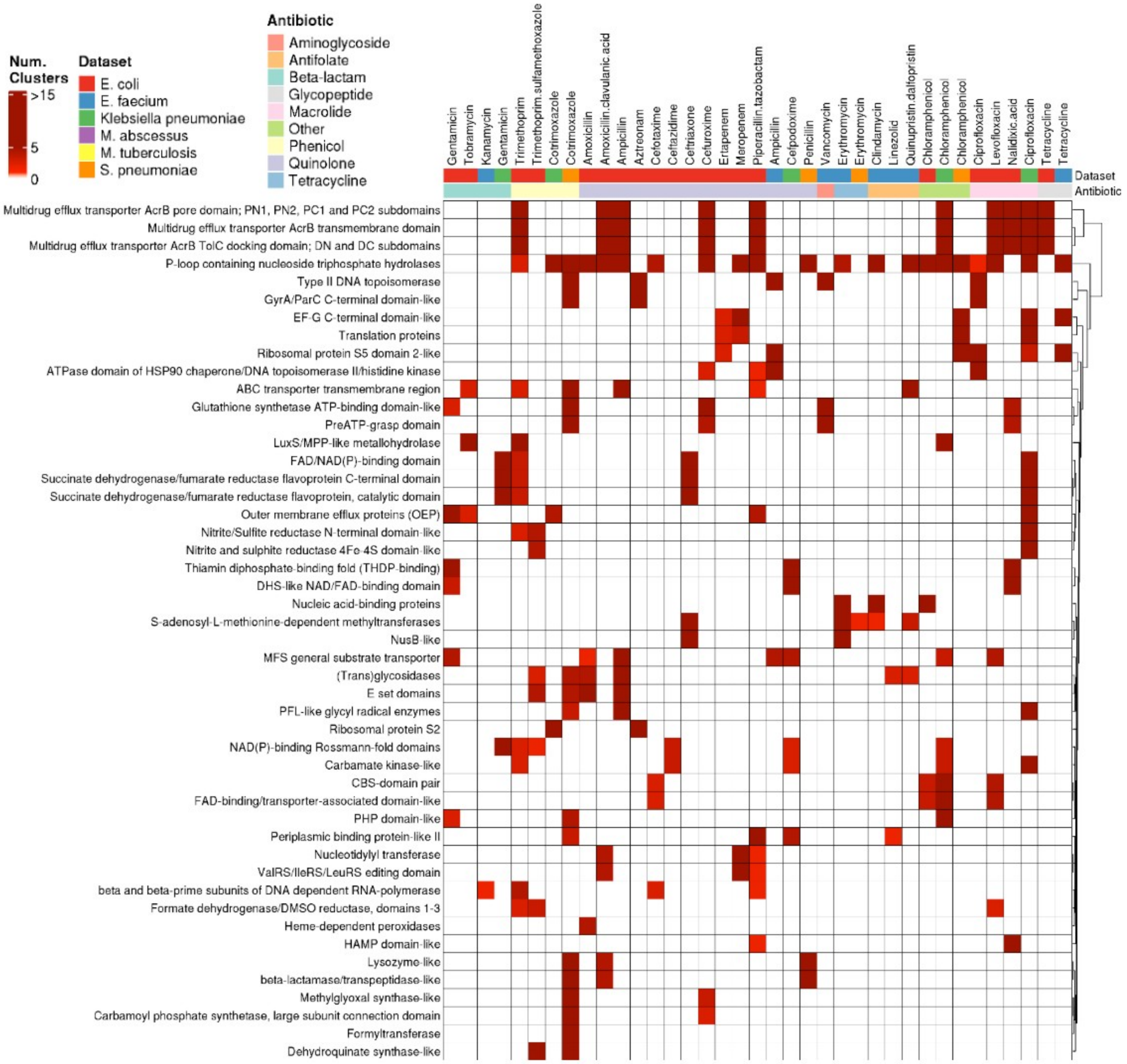


**Fig. S8. FLASH discovers shared signatures of antibiotic resistance pathways across different bacterial species.** Antibiotic annotations from InterProScan (SUPERFAMILY) database [cite] are conserved across antibiotic phenotypes in six different bacterial species: *E. Coli*, *E faecium*, *S. pneumoniae*, *K. pneumoniae, M. tuberculosis*, and *M. abscessus*. Each nonzero coefficient that contributes to a resistance phenotype and which maps to RefSeq via blast (row) is assigned an antibiotic annotation (column) using interpro scan. Rows with less than 2 different antibiotics are dropped from the visualization. The total count of different hits to a shared annotation is summed across phenotypes and the resulting heatmap is hierarchically clustered. Efflux pumps, topoisomerases and penicillin-binding proteins / other beta lactamases are seen most often across all of the antibiotic resistance phenotypes in the six species.

#### Supplemental Files

**File S1. Prediction results in vancomycin-resistant *E. faecium* sequenced via nanopore and trained on held-out short reads.** This file provides the features and their nonzero coefficients that were used to make the prediction in the *E. faecium* long nanopore reads from (Islam et al. 2023)

**File S2. All BLAST (protein and nucleotide) summaries by cluster for each metadata category across species.** This file provides all of the BLASTX and BLASTN annotation labels for every sequence in a cluster and ranked by the magnitude of that cluster in predicting the metadata category. Each sheet corresponds to a different dataset.

**File S3. Top 10 clusters and sequences for each metadata category across bacteria.** Clusters and their full set of sequences are reported for the top 10 predictive clusters for every metadata category in each species. Each sequence is joined onto the distribution of samples by metadata class given that they had that sequence in the cluster. This can be used to explore sequences which are specifically present in one phenotype and absent in another.

**File S4. Cross-species sets of proteins predictive for antibiotic resistance across six different species.** This table represents sets of proteins coming from all predictive features in *E. faecium, E. coli, S. pneumoniae, K. pneumoniae, M. tuberculosis,* and *M. abscessus* for antibiotic resistance phenotypes. Any predictive feature that successfully had a BLASTX hit in the RefSeq protein database was "clustered" with all other proteins via a pairwise alignment similarity matrix grabbing connected components of the matrix when the pairwise alignment score was above a certain threshold (methods). These clusters are labeled "sets". The remaining columns summarize properties of the features within the sets and the metadata corresponding to each unique sequence in the table. Perplexity was calculated on each short protein sequence using ESMfold (Lin et al. 2023). Entropy and Normalized Mutual Information NMI were calculated on each individual feature and averaged within each set for ranking purposes.

**File S5. After removing missing clusters, All BLAST (protein and nucleotide) summaries by cluster for each metadata category across species.** This file provides all of the BLASTX and BLASTN annotation labels for every sequence in a cluster and ranked by the magnitude of that cluster in predicting the metadata category. This was done for the predictions made after removing clusters with many missing targets across samples. Each sheet corresponds to a different dataset.

**File S6. All fungal results.** This table represents the results of running all fungal data using the effect size bin metric of reordering the clusters. The Y1000 results presented here contain the masked-aa clustering method rather than the nucleotide shift distance clustering method.

**File S7. H5N1 results after removing cattle and chicken genome contamination.** All BLAST (protein) summaries by cluster for each metadata category.

**File S8. Predictors and accuracies for both *Vibrio*-phage studies.** Contains columns corresponding to the two clusters used to create the interaction term and the annotations (if available) for each of these clusters.

**File S9. All accessions used for analyses.** Accessions listed out for each piece of data we analyzed.

**File S10. All MegaRes and Interpro SUPERFAMILY annotations for *E. faecium, E. coli, S. pneumoniae, K. pneumoniae, M. tuberculosis,* and *M. abscessus*** FLASH clusters are reported for each species as well as their annotations to the MegaRES antibiotic resistance gene database. Columns are included for each predictive cluster for each metadata category. Annotations are provided for clusters which had SPLASH Lookup Table hits to the database. The final two columns summarise the count and proportion of clusters per metadata category which had an AMR annotation.

**File S11. All aggregated data for 8mer analyses of *E. faecium*, *E. coli* and Y1000 Sacchoromycotina data.** This table represents the information from File S2 presented for 8mers in *E. coli*, *E. faecium*, and Y1000. The 8mer clusters are also ranked by magnitude.
